# Performance evaluation of RNA purification kits and blood collection tubes in the Extracellular RNA Quality Control (exRNAQC) study

**DOI:** 10.1101/2021.05.11.442610

**Authors:** The exRNAQC Consortium, Jasper Anckaert, Francisco Avila Cobos, Anneleen Decock, Philippe Decruyenaere, Jill Deleu, Katleen De Preter, Olivier De Wever, Jilke De Wilde, Bert Dhondt, Thibaut D’huyvetter, Celine Everaert, Carolina Fierro, Hetty Hilde Helsmoortel, An Hendrix, Eva Hulstaert, Jan Koster, Scott Kuersten, Tim R Mercer, Pieter Mestdagh, Annelien Morlion, Nele Nijs, Justine Nuytens, Annouck Philippron, Thomas Piofczyk, Franco Poma-Soto, Kathleen Schoofs, Gary P. Schroth, Olivier Thas, Eveline Vanden Eynde, Jo Vandesompele, Tom Van Maerken, Ruben Van Paemel, Kimberly Verniers, Jasper Verwilt, Nurten Yigit

## Abstract

The use of blood-based extracellular RNA (cell-free RNA; exRNA) as clinical biomarker requires the implementation of a validated procedure for sample collection, processing, and profiling. So far, no study has systematically addressed the pre-analytical variables affecting transcriptome analysis of exRNAs. In the exRNAQC study, we evaluated ten blood collection tubes, three time intervals between blood draw and downstream processing, and eight RNA purification methods using the supplier-specified minimum and maximum biofluid input volumes. The impact of these pre-analytics on deep transcriptome profiling of both small and messenger RNA from healthy donors’ plasma or serum was assessed for each pre-analytical variable separately and for interactions between a selected set of pre-analytics, resulting in 456 extracellular transcriptomes. Making use of 189 synthetic spike-in RNAs, the processing and analysis workflow was controlled. When comparing blood collection tubes, so-called preservation tubes do not stabilize exRNA well, and result in variable RNA concentration and sensitivity (i.e., the number of detected RNAs) over time, as well as compromised reproducibility. We also document large differences in RNA purification kit performance in terms of sensitivity, reproducibility, and observed transcriptome complexity, and demonstrate interactions between specific blood collection tubes, purification kits and time intervals. Our results are summarized in 11 performance metrics that enable an informed selection of the most optimal sample processing workflow for a given experiment. In conclusion, we put forward robust quality control metrics for exRNA quantification methods with validated standard operating procedures (SOPs), representing paramount groundwork for future exRNA-based precision medicine applications.

## Main

Biomarker studies are increasingly utilizing biofluids as an attractive resource of molecules reflecting human health or disease states. Biopsies from those human body fluids are often referred to as ‘liquid biopsies’. In contrast to tissue biopsies, they have the advantage of being minimally invasive and are compatible with serial profiling, enabling to monitor the impact of an intervention (e.g., treatment, physical exercise) over time.

Most liquid biopsy biomarker studies focus on cell-free nucleic acids as candidate biomarkers. While cell-free DNA has been studied intensively and found its way in daily clinical practice for non-invasive prenatal testing^1^, as well as for mutation and methylation detection in cancer^2^, extracellular RNA (exRNA) is relatively new in the biomarker field. Nevertheless, biomarker potential has been ascribed to various RNA molecules, including microRNA (miRNA), messenger RNA (mRNA), long-non-coding RNA and circular RNA (circRNA) in several diseases such as cancer, autoimmune diseases, diabetes, and cardiovascular diseases^3–7^. Given the labile nature of RNA and the release of exRNA and cellular RNA by cells under stress^8, 9^, the growing interest in exRNA as a biomarker resource requires the strict implementation of standardized methods for sample collection, processing and molecular profiling. Blood serum and plasma are amongst the most studied liquid biopsies and several pre-analytical variables, including blood collection tube type, needle type and blood centrifugation speed and duration, are known to influence exRNA abundance patterns (Supplementary Table 1)^10–12^. Over time, several consortia were founded with the aim to standardize some of these pre-analytical variables, including the NIH’s Extracellular RNA Communication Consortium (ERCC)^13, 14^, Blood Profiling Atlas in Cancer (BloodPAC) Consortium (www.bloodpac.org)^15,16, SPIDIA/SPIDIA4P (www.spidia.eu^) and CANCER-ID (www.cancer-id.eu). Nevertheless, pre-analytical variables are typically not reported in studies, as demonstrated by a recent literature review conducted by our group (Van Der Schueren et al., manuscript in preparation). Out of 22 studied pre-analytical variables (including the blood collection tube anticoagulant, plasma storage temperature, RNA purification method and plasma input volume for RNA purification), only 6 were sufficiently detailed in more than half of the 100 exRNA publications that were reviewed (Van Der Schueren et al., manuscript in preparation). This makes it challenging to replicate findings or directly compare biomarker studies.

While it is well recognized that pre-analytical variables need to be considered when studying exRNA biomarkers, studies investigating their impact are either focused on microRNAs or are restricted to a limited number of mRNA genes (Supplementary Table 1), and generally do not investigate interactions between pre-analytics. In the Extracellular RNA Quality Control (exRNAQC) study, we performed an extensive massively parallel sequencing-based analysis of the impact of pre-analytical variables on both extracellular small RNA and mRNA profiles. We systematically evaluated ten blood collection tubes, three time intervals between blood draw and downstream processing, and eight RNA purification methods using the supplier specified minimum and maximum plasma input volumes. The impact of these pre-analytical factors was firmly established using deep transcriptome profiling of all small and messenger RNAs from healthy donors’ plasma or serum, and was assessed by evaluating each of the pre-analytics separately (exRNAQC phase 1), as well as by analyzing interactions between pre-analytics (exRNAQC phase 2). Synthetic spike-in RNAs were added during and after RNA purification and a wide variety of (novel) performance metrics were introduced and evaluated (Fig. 1). Such a comprehensive analysis of pre-analytical variables in the context of exRNA profiling has not yet been performed (Supplementary Fig. 1).

**Fig. 1:**
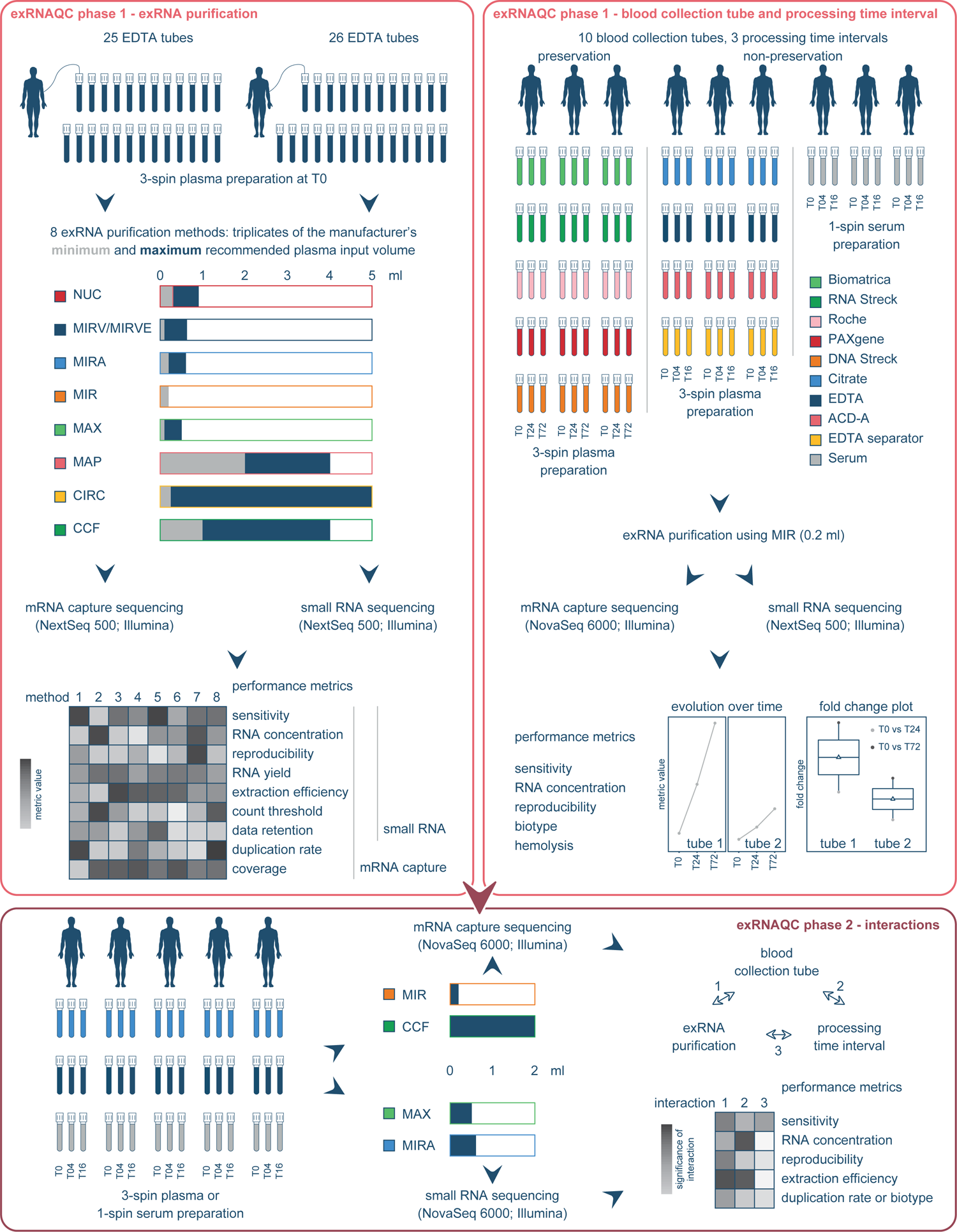
In the extracellular RNA Quality Control (exRNAQC) study, the impact of eight exRNA purification methods, ten blood collection tubes and three processing time intervals on mRNA capture and small RNA sequencing is assessed by evaluating each of the pre-analytics separately (exRNAQC phase 1), as well as by analyzing interactions between pre-analytics (exRNAQC phase 2). To evaluate the impact of the eight exRNA purification methods (upper left panel), two blood draws from a single individual were performed to separately apply mRNA capture and small RNA sequencing. To compare RNA purification performance, nine performance metrics were calculated (see Methods). To evaluate the impact of the ten blood collection tubes (upper right panel), nine individuals were sampled, enabling to test three time intervals between blood draw and downstream processing for each of the tubes. Preservation tubes were processed immediately upon blood collection (T0), after 24 hours (T24) or after 72 hours (T72). Non-preservation plasma and serum tubes were processed immediately upon blood collection (T0), after 4 hours (T04) or after 16 hours (T16). Both mRNA capture and small RNA sequencing were performed, and the data was analyzed using five performance metrics (see Methods). Based on the sensitivity and reproducibility metrics, a dedicated selection of precise and sensitive exRNA purification methods and blood collection tubes was put forward for further evaluation in exRNAQC phase 2. For both mRNA capture and small RNA sequencing, five individuals were sampled to test three blood collection tubes and four exRNA purification methods. Interactions between exRNA purification methods, blood collection tubes and processing time intervals were assessed by the evaluation of six performance metrics (see Methods). ACD-A: BD Vacutainer Glass ACD Solution A tube; Biomatrica: LBgard Blood Tube; CCF: QIAamp ccfDNA/RNA Kit; CIRC: Plasma/Serum Circulating and Exosomal RNA Purification Kit/Slurry Format; Citrate: Vacuette Tube 9 ml 9NC Coagulation sodium citrate 3.2%; DNA Streck: Cell-Free DNA BCT; EDTA: BD Vacutainer Plastic K2EDTA tube; EDTA separator: Vacuette Tube 8 ml K2E K2EDTA Separator; MAP: MagNA Pure 24 Total NA Isolation Kit in combination with the MagNA Pure instrument; MAX: the Maxwell RSC miRNA Plasma and Exosome Kit in combination with the Maxwell RSC Instrument; MIR: the miRNeasy Serum/Plasma Kit; MIRA: the miRNeasy Serum/Plasma Advanced Kit; MIRV: the mirVana PARIS Kit with purification protocol for total RNA; MIRVE: mirVana PARIS Kit with purification protocol for RNA enriched for small RNAs; NUC: the NucleoSpin miRNA Plasma Kit; PAXgene: PAXgene Blood ccfDNA Tube; RNA Streck: Cell-Free RNA BCT; Roche: Cell-Free DNA Collection Tube; Serum: BD Vacutainer SST II Advance Tube.

## Results

### RNA purification methods influence miRNA and mRNA abundance profiles

In the first phase of the exRNAQC study, eight total RNA purification methods were selected for evaluation (Fig. 1): the miRNeasy Serum/Plasma Kit (abbreviated to MIR), miRNeasy Serum/Plasma Advanced Kit (MIRA), mirVana PARIS Kit (MIRV), NucleoSpin miRNA Plasma Kit (NUC), QIAamp ccfDNA/RNA Kit (CCF), Plasma/Serum Circulating and Exosomal RNA Purification Kit/Slurry Format (CIRC), Maxwell RSC miRNA Plasma and Exosome Kit in combination with the Maxwell RSC Instrument (MAX), and MagNA Pure 24 Total NA Isolation Kit in combination with the MagNA Pure instrument (MAP). Since most methods allow a range of blood plasma input volumes, we tested both the minimum and maximum input volume recommended by each supplier (indicated in ml following the RNA purification method abbreviation). For the mirVana PARIS Kit, also an alternative protocol for specific enrichment of small RNAs (MIRVE) was tested for small RNA purification. Blood was collected (in EDTA tubes) from a healthy donor and three technical replicates were used for every kit-plasma input volume combination, resulting in 45 and 51 samples that were processed for RNA extraction and mRNA capture or small RNA sequencing library preparation, respectively.

We first investigated potential DNA contamination in the RNA eluates using the strandedness of the mRNA capture sequencing data as a proxy. As we applied a stranded sequencing library preparation protocol, strandedness should be close to 100% in the absence of DNA contamination. Strandedness for libraries from MAP purified RNA, however, was considerably lower: only 70-75% and 80-85% of reads mapped to the correct strand for MAP2 and MAP4 purification, respectively, while this percentage was above 95% for all other purification methods (Supplementary Fig. 2c). Moreover, the small RNA sequencing data from MAP contained a much higher fraction of mapped reads that did not overlap annotated small RNA sequences (35 to 52% of mapped reads for MAP compared to only 1 to 6% for other purification kits) and more than 80% of these unannotated reads did not overlap with known exons. Despite DNase treatment, these findings strongly suggest residual DNA contamination in MAP RNA eluates and we therefore excluded this purification method from all further analyses.

To evaluate performance differences among RNA purification methods, we calculated nine purposely developed metrics (see Methods and Table 1): (1) sensitivity, (2) RNA concentration, (3) RNA yield, (4) extraction efficiency, (5) count threshold, (6) data retention, (7) reproducibility, (8) duplication rate, and (9) coverage (see Methods for a detailed description of each individual metric; the last two metrics were only evaluated for the mRNA data). In terms of **sensitivity**, the absolute number of mRNAs and miRNAs detected ranged from 989 to 11,322 and from 69 to 171, respectively. While a higher plasma input volume consistently resulted in a higher number of detected mRNAs for a given method (Fig. 2a & b), this was not always true when comparing different methods e.g., MIRA0.6 (7424 mRNAs on average, 0.6 ml) versus NUC0.9 (4766 mRNAs on average, 0.9 ml); Fig. 2a). This also holds true for miRNAs, except for CCF, CIRC and NUC (Fig. 2b). To compare the eluate **RNA concentration** of the different RNA purification methods, the ratio of endogenous counts versus ERCC spikes counts (for mRNA capture sequencing) and endogenous counts versus LP spikes counts (for small RNA sequencing) were determined. This sequencing-based RNA concentration metric correlates significantly with RNA concentrations determined by Femto Pulse electropherogram analysis (p-value < 0.001, Spearman correlation coefficient of 0.67 for mRNA capture sequencing and 0.86 for small RNA sequencing, Supplementary Fig. 3a & b, respectively). Femto Pulse analyses further demonstrate that blood-derived exRNA is highly fragmented (Supplementary Fig. 4). The purification method resulting in the highest mRNA concentration (CCF4) had on average a 76 times higher eluate concentration than the kit with the lowest concentration (MIRV0.1) (Fig. 2c). For miRNAs, the difference was even larger, with a 238 times higher concentration in CCF4 compared to MIRVE0.1 (Fig. 2d). When excluding MIRVE, a kit not tested at mRNA level, the difference between the kit with the highest and lowest miRNA concentration was 29-fold. The **RNA yield** metric represents the relative amount of RNA in the total eluate volume after purification. For mRNA capture sequencing, there was on average a 30-fold difference between the kit with the highest RNA yield (CIRC5) compared to the kit with the lowest RNA yield (NUC0.3) (Supplementary Fig. 5e). For small RNA sequencing, there was on average an 85-fold difference between the kit with the highest RNA yield (MAX0.5) compared to the kit with the lowest RNA yield (MIRVE0.1) (Supplementary Fig. 5f). **Extraction efficiency** is a performance metric that, besides RNA yield, also takes into account differences in plasma input volume for RNA purification. It is a relative measure of how well a certain kit purifies RNA from the applied plasma input volume. When looking at the extracellular mRNA transcriptome, the highest average extraction efficiency (MAX0.1) was ten times higher than the lowest (MIRV0.1) (Supplementary Fig. 5g). For small RNAs, the highest average extraction efficiency (MAX0.1) was 25 times higher than the lowest (MIRV0.625) (Supplementary Fig. 5h). Of note we did not observe differences in extraction efficiency between the maximum and minimum input volume for a given kit. For each purification kit, we further determined a **count threshold** to filter noisy data based on eliminating 95% of single positive observations between technical replicates. Higher count thresholds indicate higher variability. This threshold varied from 5 to 14 counts at mRNA level for CCF4 and MIRV0.1, respectively, and from 2 to 16 counts at miRNA level for MIRA0.6 and MIRVE0.1, respectively (Supplementary Fig. 5a & b; Supplementary Table 2). A related performance metric, **data retention**, represents the fraction of total counts that are retained after applying the count threshold. For mRNA capture sequencing, data retention ranged from an average of 93.5% in MIRV0.1 to an average of 99.7% in CCF4 (Supplementary Fig. 5c). For small RNA sequencing, data retention ranged from an average of 98.8% in MIRVE0.1 to an average of 99.8% in MAX0.5 (Supplementary Fig. 5d). To assess **reproducibility**, we determined the area left of the curve (ALC), a robust metric based on differences in mRNA or miRNA counts between technical replicates (as defined in the miRQC study^17^, see Methods). The higher the reproducibility, the lower the ALC value. Most kits performed equally well with respect to miRNA count reproducibility (Fig. 2f) except for MIRVE0.1. For mRNA, CIRC0.25 and MIRV0.1 displayed a lower reproducibility than the other kits, while CCF4 had the best reproducibility, closely followed by CIRC5 and MIR0.2 (Fig. 2e). Within a kit, the maximum plasma input volume consistently resulted in a better reproducibility compared to the minimum plasma input volume. A low amount of input RNA, as is the case for biofluids, typically results in mRNA capture sequencing libraries with a high fraction of PCR duplicates. The average **duplication rate** ranged from 82.2% (CCF4) to 97.3% (NUC0.3) of mRNA capture sequencing reads (lower is better, Supplementary Fig. 2a). Note that even a small difference in duplication rate can have a high impact on the total number of non-duplicated reads: with CCF4, on average six times more non-duplicated reads were generated compared to NUC0.3 (Supplementary Table 3). Finally, transcriptome **coverage** was determined to assess the diversity of mRNA capture sequencing reads. The MIRV0.1 kit had the lowest average coverage: only 1.8% of the human Ensemble v91 transcriptome was covered by at least one sequencing read. Purification with CCF4 resulted in the highest average coverage (17.7%, Supplementary Fig. 2b).

**Fig. 2:**
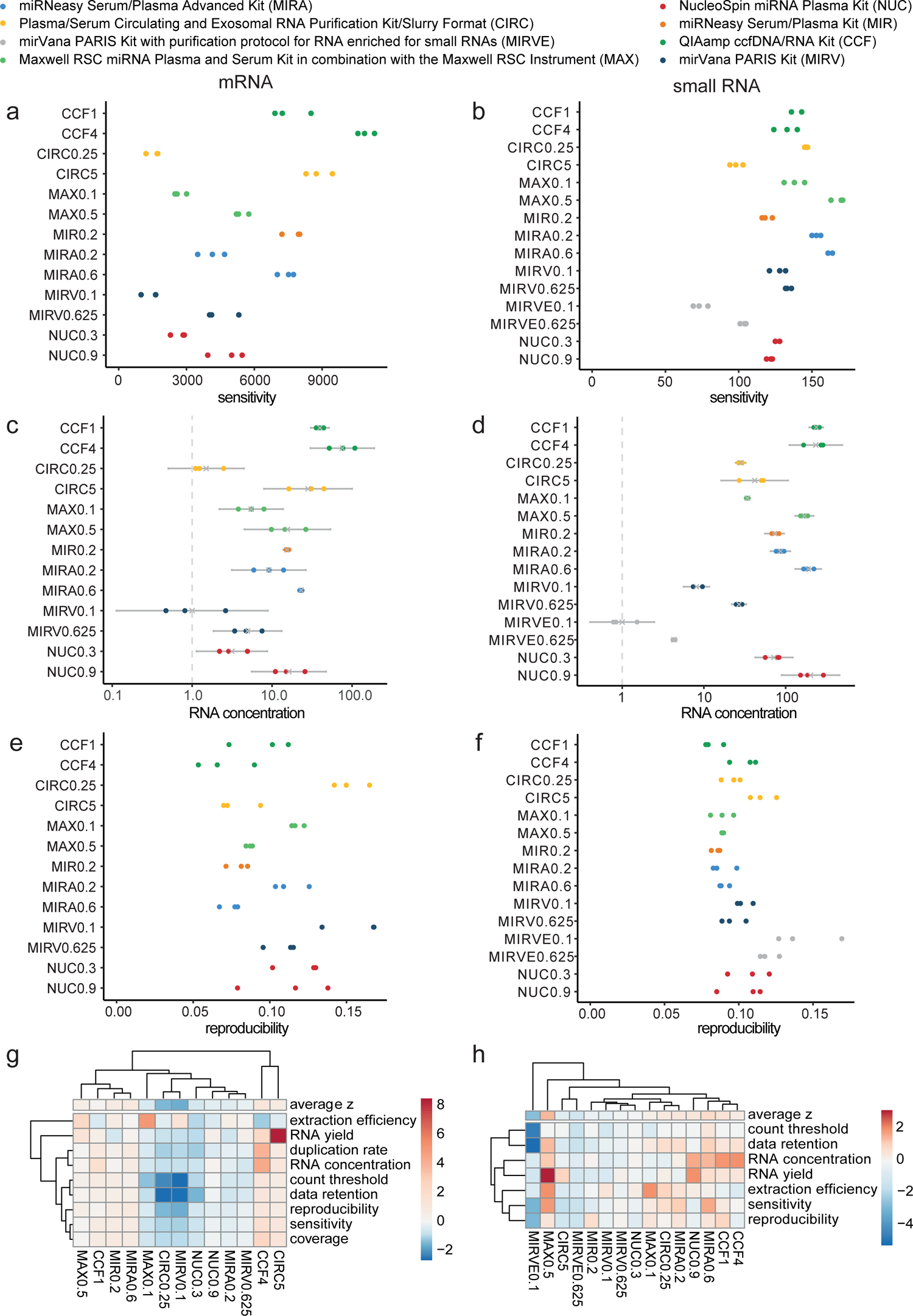
RNA purification methods strongly influence mRNA and small RNA sequencing. For both mRNA capture (left panels) and small RNA (right panels) sequencing, performance metrics are shown. For each of the unique RNA purification-plasma input volume combinations, 3 technical replicates are analyzed (n = 39 for mRNA capture sequencing, n = 45 for small RNA sequencing). (**a&b**) Absolute number of mRNAs and miRNAs, respectively, that reach the count threshold (i.e., sensitivity). (**c&d**) Endogenous RNA concentration. Values are log rescaled to the lowest mean of all kits and transformed back to linear scale. The mean and 95% confidence interval are shown. (**e&f**) Reproducibility between technical replicates based on ALC (smaller ALC indicates better reproducibility) at mRNA and miRNA level, respectively. (**g&h**) Overview of all performance metrics at mRNA capture and small RNA sequencing level, respectively, after transforming the values to robust z-scores. A higher z-score indicates a better performance. Rows and columns of the heatmaps are clustered according to complete hierarchical clustering based on Euclidean distance. Average z refers to the mean of robust z-scores for a specific RNA purification method. The number that follows the abbreviation of the purification kit is the plasma input volume (in ml).

**Table 1.**
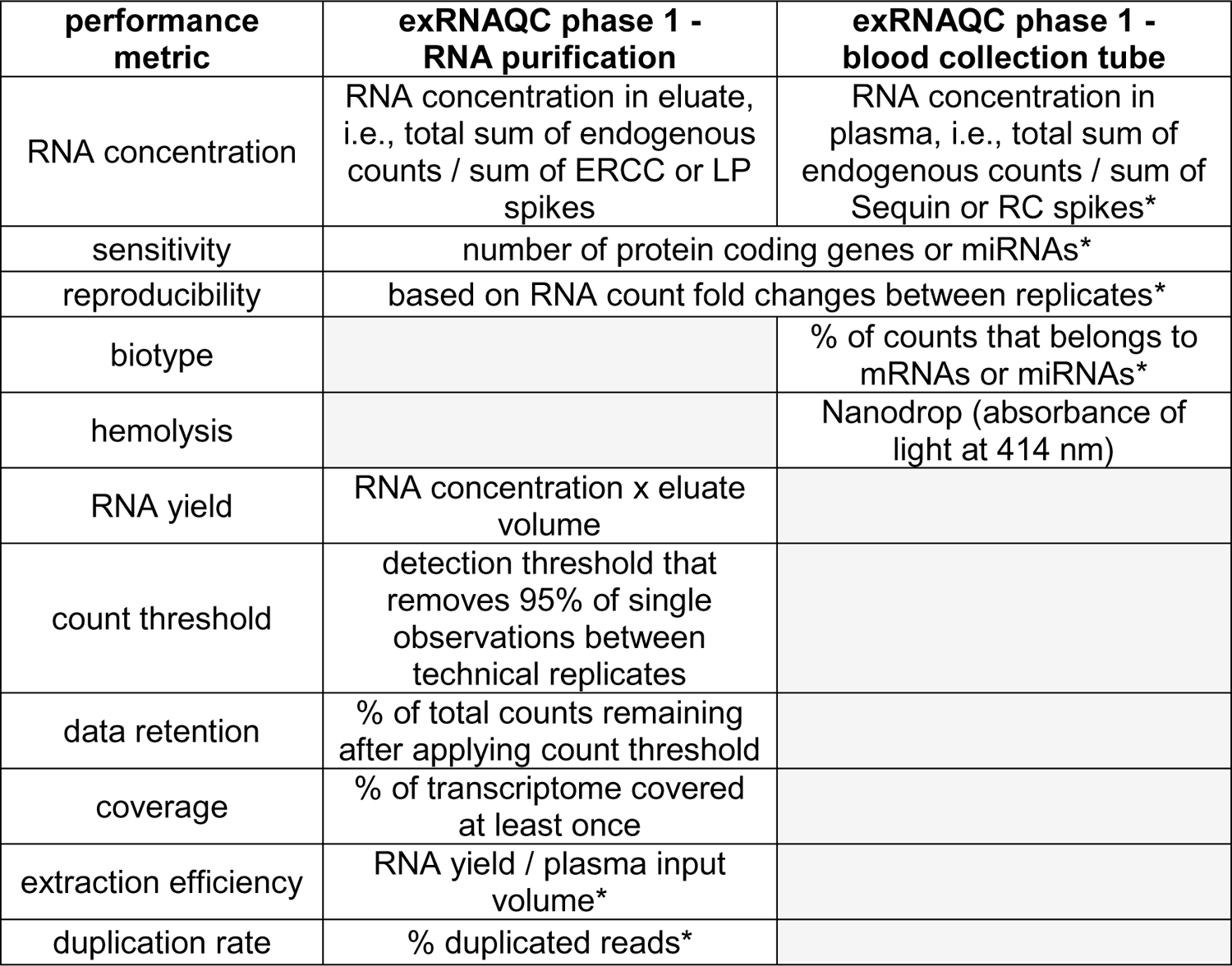
In total, 11 performance metrics are determined to evaluate the impact of RNA purification methods and blood collection tubes on mRNA capture and small RNA sequencing results. An overview of the different performance metrics is given. Gray indicates that the performance metric is not calculated. Metrics marked with an * were also calculated in exRNAQC phase 2. ERCC: Extracellular RNA Communication Consortium; LP: Library Prep Control; RC: RNA extraction Control.

A summary plot of all performance metrics after robust z-score transformation is shown in Fig. 2g & h, for mRNA and small RNA, respectively. For each metric, a higher z-score indicates a better performance. In general, kit differences are smaller for miRNA than for mRNA (less variability in z-score). For mRNA capture sequencing, kits with a higher plasma input volume, such as CIRC5 and CCF4, scored better on most performance metrics. Kits with plasma input volumes below 0.5 ml were in general less performant than other kits, except for MIR0.2. Despite lower performance scores, MAX0.1 and MIRV0.1 were efficient in purifying RNA from the given 0.1 ml of plasma. Of note, plasma input volume alone does not completely determine performance as some kits with a lower plasma input volume (for example MIRA0.6 and CCF1) still perform better than kits with a higher input. Similarly, for small RNAs, we mainly observe low performance in the lower input volume kits, but there were also exceptions. MAX0.5 and MIRA0.6, for example, scored surprisingly well or even better compared to kits with a much larger plasma input volume such as CIRC5 and CCF4. In contrast to mRNA capture sequencing, more plasma input for a given kit did not always result in better small RNA sequencing performance (see CIRC5 vs CIRC0.25).

### Blood preservation tubes are not suitable for exRNA analysis

In addition to RNA purification methods, different blood collection tubes and processing time intervals were evaluated in exRNAQC phase 1. Ten blood collection tubes were selected, belonging to two categories: tubes not designed to stabilize nucleic acids (which we termed ‘non-preservation tubes’; n = 5), and so-called ‘preservation tubes’ (n = 5) that are purposely developed to conveniently allow more time between the blood draw and further processing steps. The selected non-preservation tubes were the BD Vacutainer Plastic K2EDTA tube (abbreviated to EDTA), Vacuette Tube 8 ml K2E K2EDTA Separator (EDTA separator), BD Vacutainer Glass ACD Solution A tube (ACD-A), Vacuette Tube 9 ml 9NC Coagulation sodium citrate 3.2% (Citrate), and BD Vacutainer SST II Advance Tube (Serum). The preservation tubes were the Cell-Free RNA BCT (RNA Streck), Cell-Free DNA BCT (DNA Streck), PAXgene Blood ccfDNA Tube (PAXgene), Cell-Free DNA Collection Tube (Roche) and LBgard Blood Tube (Biomatrica). For each of the blood collection tubes, we recruited three healthy volunteers and selected three time intervals between blood draw and processing: immediately (T0), time interval 1 (4 hours (T04) for non-preservation tubes, 24 hours (T24) for preservation tubes), and time interval 2 (16 hours (T16) for non-preservation tubes and 72 hours (T72) for preservation tubes). This resulted in a total of 180 samples that were subsequently processed for RNA purification (using MIR0.2) and mRNA capture or small RNA sequencing.

To evaluate exRNA profilesbetween tubes and over time, we calculated five different performance metrics: (1) hemolysis, (2) RNA concentration, (3) sensitivity, (4) biotype, and (5) reproducibility (see Methods and Table 1). The stability of each performance metric over time was evaluated as a fold change between the first (T0) and the second (T04 or T16) or third time interval (T16 or T72), subsequently, as exemplified in Supplementary Fig. 6. If the processing time interval has no impact on any of the above-described metrics, respective fold changes should be close to one. For each blood collection tube, the average fold change of each performance metric over time is shown in Fig. 3. **Hemolysis** was quantified based on absorbance units at 414 nm and evaluated by visual inspection during liquid biopsy preparation. For the non-preservation tubes, hemolysis measurements were below the generally accepted absorbance threshold of 0.2^18, 19^ across all donors and time intervals (Supplementary Fig. 7a, Supplementary Fig. 8a and Supplementary Fig. 9). In contrast, for all preservation tubes except the Biomatrica tube, plasma was hemolytic for at least one donor at T0. At T72, the Biomatrica hemolysis measurements also exceeded the 0.2 threshold. Despite the low absorbance values, we did observe up to two-fold differences in function of time: mean fold changes in non-preservation tubes ranged from 1.05 to 2.04, and in preservation tubes from 1.19 to 2.08 (Supplementary Fig. 10a & Supplementary Fig. 11a). To assess **RNA concentration** differences in plasma prepared from the different blood collection tubes, ratios of endogenous counts versus Sequin spikes counts (for mRNA capture sequencing) and endogenous counts versus RC spikes counts (for small RNA capture sequencing) were calculated. RNA concentration in non-preservation tubes remained quite stable over time, with a 1.23 to 1.48 fold increase in mRNA mass and a 1.57 to 2.97 fold increase in miRNA mass (Supplementary Fig. 10b & Supplementary Fig. 11b). Unexpectedly, RNA concentration was much less stable in preservation tubes with fold changes of 1.84 to 4.03 and 1.75 to 10.50 for mRNA and small RNA, respectively. While RNA concentration did not change substantially between time intervals for the RNA Streck tubes, the RNA concentration at the individual time intervals for these tubes was substantially lower compared to the other tubes (on average 4.97-fold lower for mRNA and 10.36-fold lower for small RNA (Supplementary Fig. 7b & Supplementary Fig. 8b)). The **sensitivity** (i.e., the absolute number of mRNAs and miRNAs) in non-preservation tubes remained relatively constant over time: mean fold changes ranged from 1.29 to 1.59 and from 1.10 to 1.36 at mRNA and small RNA sequencing level, respectively. In preservation tubes, the mean fold change ranged from 1.86 to 4.01 and from 1.08 to 1.67 for mRNA and miRNA, respectively (Supplementary Fig. 10c & Supplementary Fig. 11c). Furthermore, and similar to the RNA concentration, the sensitivity was substantially lower in DNA Streck and RNA Streck tubes compared to the others (mean number of mRNAs: 385 for RNA Streck and 840 for DNA Streck; mean number of miRNAs: 60 for RNA Streck and 103 for DNA Streck) (Supplementary Fig. 7c & Supplementary Fig. 8c). The fraction of total counts mapping to mRNAs and miRNAs (Supplementary Fig. 7d & Supplementary Fig. 8d), referred to as the **biotype** performance metric, in non-preservation tubes remained fairly constant over time: mean fold changes ranged from 1.08 to 1.14 and from 1.13 to 1.47, for mRNA and miRNA, respectively. For the preservation tubes, the mean fold changes were higher: from 1.69 to 2.28 and from 1.38 to 4.52, for mRNA and miRNA, respectively (Supplementary Fig. 10e & Supplementary Fig. 11e). **Reproducibility** remained stable over time for both preservation and non-preservation tubes: mean fold changes ranged from 1.06 to 1.18 (Supplementary Fig. 10d & Supplementary Fig. 11d). In general, the stability of the performance metrics over time was substantially better for the non-preservation tubes compared to the preservation tubes (Fig. 3).

**Fig. 3.**
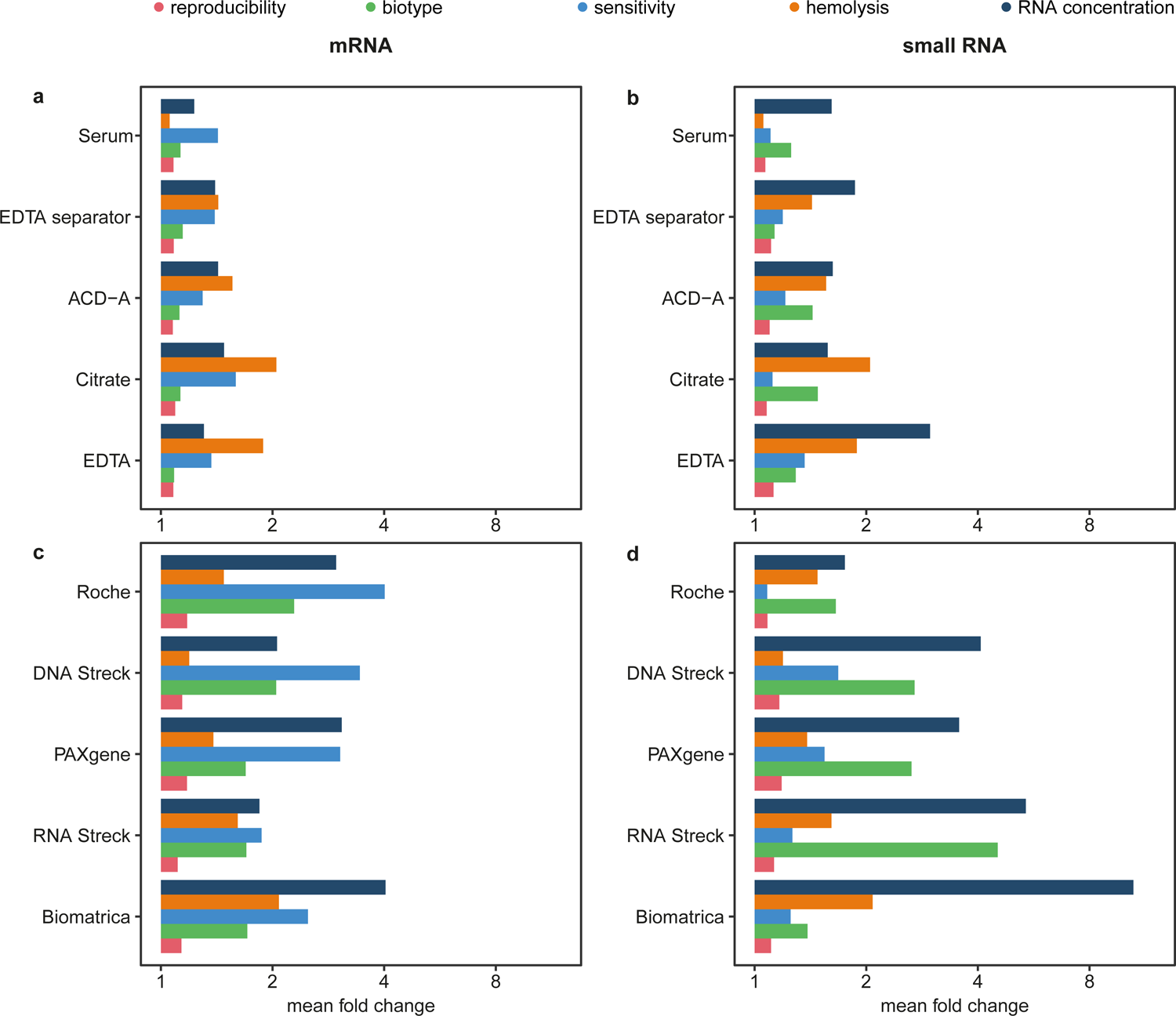
The tested blood preservation tubes are not suitable for exRNA analysis. Per blood collection tube and per performance metric, a summary of mean fold changes (FC) between time interval 1 (i.e., T04 (blood processing starts 4 hours after blood collection) for non-preservation tubes and T24 for preservation tubes) and time interval 0 (T0) versus time interval 2 (i.e., T16 for non-preservation tubes and T72 for preservation tubes) and time interval 0 (T0) is given, for both mRNA capture sequencing (**a&c**) and small RNA sequencing (**b&d**). Ideally, the mean FC of the stability metrics approaches 1, indicating that there is little change from baseline and the blood collection tube performs well over time. Non-preservation blood tubes are the BD Vacutainer Plastic K2EDTA tube (EDTA), Vacuette Tube 8 ml K2E K2EDTA Separator (EDTA separator), BD Vacutainer Glass ACD Solution A tube (ACD-A), Vacuette Tube 9 ml 9NC Coagulation sodium citrate 3.2% (Citrate), and BD Vacutainer SST II Advance Tube (Serum). The preservation tubes are the Cell-Free RNA BCT (RNA Streck), Cell-Free DNA BCT (DNA Streck), PAXgene Blood ccfDNA Tube (PAXgene), Cell-Free DNA Collection Tube (Roche) and LBgard Blood Tube (Biomatrica). Note that different donors were sampled and that tubes were processed at different time intervals for preservation and non-preservation blood tubes.

Tubes were further evaluated by determining the circRNA and linear RNA fractions at the different time intervals for each of the tubes separately, as well as by comparing RNA abundance levels across tubes (at T0) and time intervals, and by evaluating differences in immune cell RNA contributions over time. Fractions of circRNA and linear RNA do not significantly differ across time intervals (pairwise comparisons using Wilcoxon Rank Sum test with Holm-Bonferroni adjustment, all adjusted p-values > 0.05; Supplementary Fig. 12). After normalization and scaling of the count data, transcript abundance levels of all genes for the different tubes at time interval T0 were compared and visualized in a heatmap (Supplementary Fig. 13). The Roche, Biomatrica and RNA Streck tubes cluster separately from the other tubes. To assess tube stability across time intervals, distributions of log2 fold change differences between time interval 1 (T04 or T24) and 0 versus time interval 2 (T16 or T72) and 0 were compared and GSEA was performed (Supplementary Table 4). Clearly, preservation tubes show higher log2 fold change differences compared to non-preservation tubes, but also non-preservation tubes have, although less outspoken, higher log2 fold changes at the second time interval compared to the first one (Supplementary Fig. 14a and Supplementary Table 4). Non-preservation tubes that do not show significantly enriched gene sets when processed within four hours upon blood draw are EDTA and Citrate (Supplementary Table 4). Computational deconvolution of exRNA profiles (see Methods) revealed that the estimated proportions of several cell types change over time, in a tube-dependent manner (**Fig. 4**). Remarkably, Serum, Citrate and ACD-A seem to maintain a stable immune cell composition over time. Note that, in contrast to Serum, Citrate and ACD-A show no significant changes if blood processing is performed within four hours upon blood draw.

**Fig. 4:**
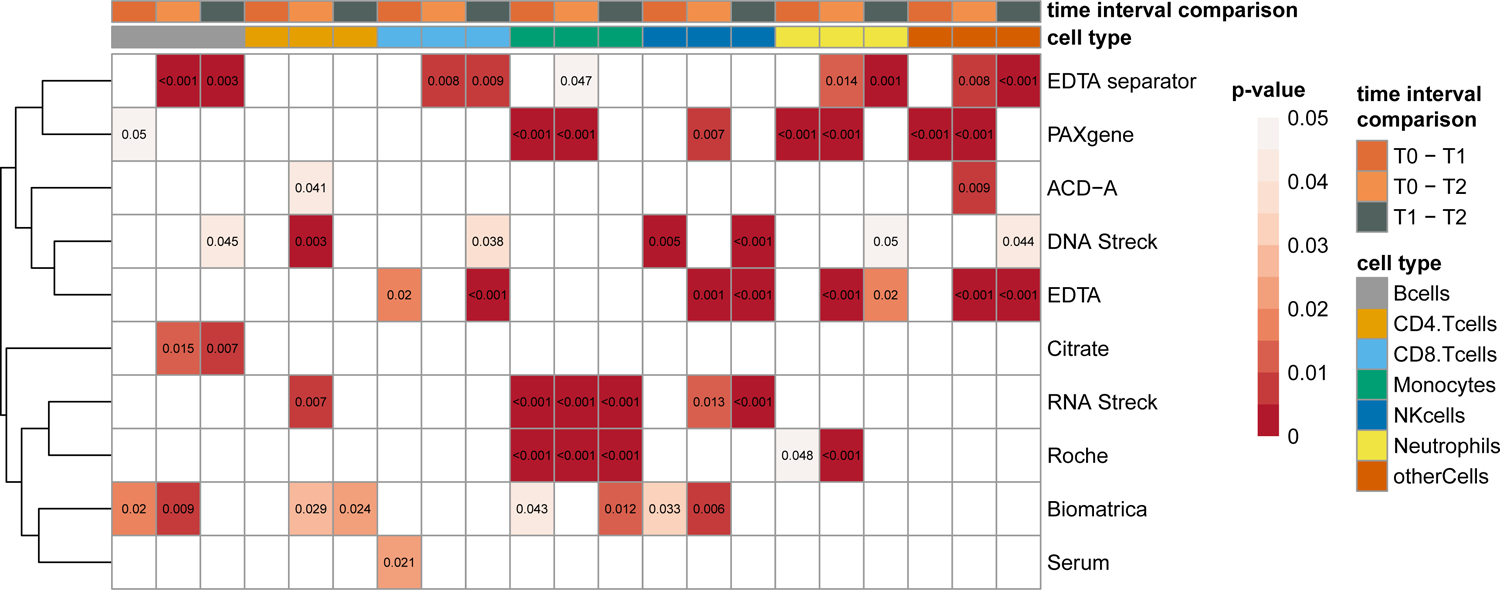
Changes in immune cell composition over time. Colored cells represent adjusted p-values < 0.05 (Tukey’s method) from beta regression models with random effects for all cell types. Time interval 0 corresponds to plasma prepared immediately after blood draw. For the non-preservation tubes, time interval 1 corresponds to T04 and time interval 2 to T16. For the preservation tubes, time interval 1 corresponds to T24 and time interval 2 to T72. T04, T16, T24, T72: plasma prepared 4, 16, 24 and 72 hours after blood draw, respectively. Note that different donors were sampled and that tubes were processed at different time intervals for preservation and non-preservation tubes. ACD-A: BD Vacutainer Glass ACD Solution A tube; Biomatrica: LBgard Blood Tube; Citrate: Vacuette Tube 9 ml 9NC Coagulation sodium citrate 3.2%; DNA Streck: Cell-Free DNA BCT; EDTA: BD Vacutainer Plastic K2EDTA tube; EDTA separator: Vacuette Tube 8 ml K2E K2EDTA Separator; PAXgene: PAXgene Blood ccfDNA Tube; RNA Streck: Cell-Free RNA BCT; Roche: Cell-Free DNA Collection Tube; Serum: BD Vacutainer SST II Advance Tube.

### Interactions between pre-analytics should be considered when comparing kit or tube performance

In the second phase of the exRNAQC study, we evaluated if performance interactions between pre-analytic variables exist. More specifically, we evaluated whether the impact of a certain pre-analytic variable on exRNA sequencing outcome depends on the choice of other pre-analytical variables. To this purpose, three non-preservation blood collection tubes and two RNA purification kits were specifically selected for further evaluation (Fig. 1). The tube selection (Serum, EDTA and Citrate) was based on their superior performance in phase 1 and on their widespread use in the clinic. The kit selection was based on their sensitivity (Fig. 2a & b) and reproducibility (Fig. 2e & f) from phase 1 (Supplementary Fig. 15). Plasma input volume was used as an additional criterion, as we aimed to include at least one kit that requires less than one milliliter biofluid. Because of the differences in kit performance on mRNA and miRNA level, MIR0.2 and CCF2 were selected for probing the mRNA transcriptome and MAX0.5 and MIRA0.6 for the small RNA transcriptome. For evaluation of the different blood collection tube- kit combinations in exRNAQC phase 2, blood was drawn from five healthy individuals and processed at three time intervals (immediately (T0), 4 hours (T04) or 16 hours (T16)), resulting in 180 samples processed for RNA purification and mRNA capture or small RNA sequencing. Interactions were analyzed using six relevant performance metrics: (1) duplication rate, (2) RNA concentration, (3) extraction efficiency, (4) sensitivity, (5) reproducibility and (6) biotype. Significant interactions between pre-analytic variables on mRNA and miRNA level are summarized in Fig. 5a & b, respectively. For mRNA, the number of detected genes (**sensitivity**) was significantly lower (adjusted p-value = 0.004) in the Serum tube compared to the other tubes, but only if extracted with CCF2 (Fig. 5c). The Serum tube also resulted in higher **duplication rates** at time interval T16 when purified with CCF2 (adjusted p-value of 0.048), albeit with small effect sizes (95% for serum and 92/91% for EDTA/citrate). We observed a significantly higher **RNA concentration** and **extraction efficiency** for the EDTA tube at time interval T16 compared to other time intervals. In terms of reproducibility, no significant interactions were observed. Also, between RNA purification kits a clear difference in plasma RNA concentration was observed, which is unexpected considering that the same biofluid was used for both kits. This suggests a difference in purification efficiency between MIR0.2 and CCF2. Apart from the interaction analyses, also RNA abundance analyses and GSEA across time intervals were performed, and confirmed results from phase 1; for these selected non-preservation tubes higher log2 fold changes (log 2 FC) were observed at the second time interval compared to the first one (Supplementary Fig. 14b and Supplementary Table 4), except for the Serum-CCF2 combination. Differential genes (|log 2 FC| > 1 and Benjamini-Hochberg adjusted p-value < 0.05) were only detected for the T0-T16 comparison and ranged from 0/6386 genes (Serum-CCF2) to 280/7876 genes (Citrate and EDTA in combination with CCF2). On the other hand, differential gene sets were observed for both time interval comparisons, although lower numbers of significant gene set changes (Benjamini-Hochberg adjusted p-value < 0.05) were detected for the T0-T04 comparison (0-17 gene sets) compared to the T0-T16 comparison (0-92 gene sets). For small RNAs, more significant interactions were found compared to mRNA. EDTA tubes were found to have significantly lower **sensitivity** compared to the other tubes (adjusted p-value < 0.001), but only if extracted with MAX0.5. Conversely, when extracted with MIRA0.6, Citrate was the tube with lower sensitivity (adjusted p-value < 0.001). For both purification kits, Serum tubes showed the highest sensitivity. Sensitivity was also significantly lower for EDTA and Citrate tubes at time intervals T0 (adjusted p-value < 0.003) and T16, respectively (adjusted p-value < 0.001). EDTA at time interval T16 showed significantly higher **RNA concentration** and **extraction efficiency** (adjusted p-value < 0.001). Moreover, EDTA also showed higher extraction efficiency compared to other tubes when MAX0.5 purification was used. Significant interactions of tube type with purification kit and time interval were also observed for the **biotype** metric. The miRNA fraction in Serum was significantly lower when the purification was done using MAX0.5 (adjusted p-value < 0.001). On the other hand, RNA extraction of Citrate plasma with the MIRA0.6 kit resulted in the lowest miRNA fraction (adjusted p-value < 0.001). Finally, the **reproducibility** for Citrate was significantly higher only when extracted with the MIRA0.6 kit (adjusted p-value < 0.001).

**Fig. 5:**
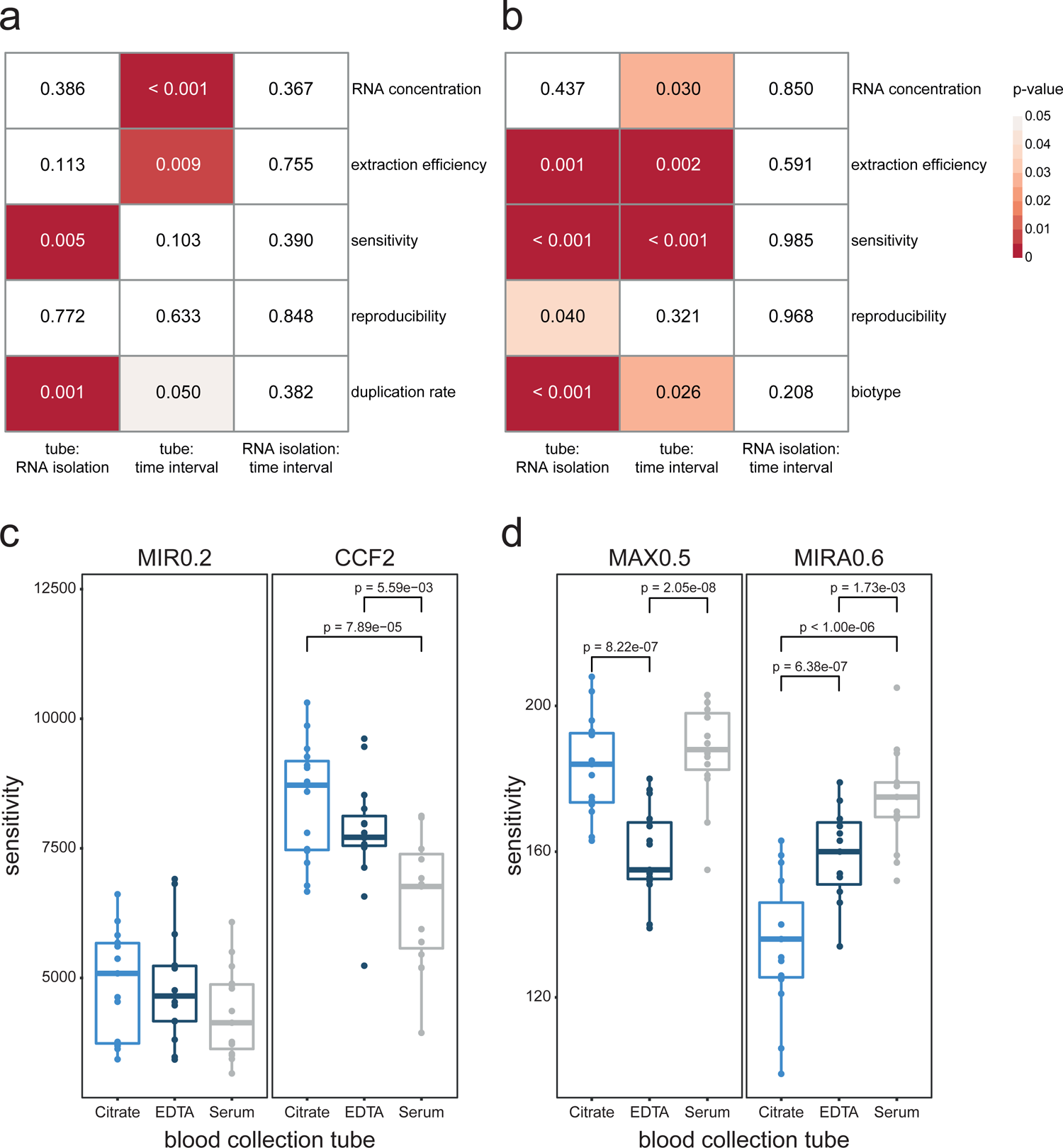
Interactions between pre-analytics should be considered when comparing RNA purification kit or blood collection tube performance. For both mRNA capture and small RNA sequencing, 5 biological replicates were used for each of the 18 unique tube (n = 3), purification method (n = 2) and time interval (n = 3) combinations (total n = 90). Shown are the interactions between pre-analytics for mRNA capture (a) and small RNA (b) sequencing. P-values correspond to the Wald test for the terms in the linear mixed-effects model. Example of a significant blood collection tube – kit interaction on mRNA (c) and small RNA (d) level. Adjusted p-values were calculated using Tukey’s method for comparing a family of 3 estimates. In the boxplots, the lower and upper hinge of the boxes represents the 25th and 75th percentile, respectively. The whiskers extend to the lowest and highest value that is within 1.5 times the interquartile range. Data beyond the end of the whiskers are outliers. CCF: QIAamp ccfDNA/RNA Kit; MAX: Maxwell RSC miRNA Plasma and Exosome Kit in combination with the Maxwell RSC Instrument; MIR: miRNeasy Serum/Plasma Kit; MIRA: miRNeasy Serum/Plasma Advanced Kit.

## Discussion

In the extracellular RNA Quality Control (exRNAQC) study, we examined eight RNA purification methods, ten blood collection tubes, and three time intervals as pre-analytical variables affecting exRNA quantification, using full transcriptome mRNA and small RNA sequencing. Eight kits marketed for RNA purification from serum or plasma and ten blood collection tubes commonly used in the clinic and available at study initiation were selected for investigation. More than 1.6 liter of blood was collected from 20 different healthy donors to conduct experiments in triplicate (exRNAQC phase 1) or quintuplicate (exRNAQC phase 2), resulting in 456 extracellular transcriptomes. To control the RNA purification and library preparation workflows, 189 synthetic spike-in RNA molecules (Sequin and ERCC spike-ins for mRNA capture sequencing, and RC and LP spike-ins for small RNA sequencing) were used. We previously demonstrated the importance of using these spike-in RNAs for sequencing-based quantification of exRNA^20, 21^, and further confirmed their critical importance in the current exRNAQC study. Here, spike-in RNAs were used to assess the RNA concentration and yield, and to determine the extraction efficiency of the different RNA purification methods. Importantly, we provide full access to the data and analysis pipelines through the European Genome-phenome Archive (EGA), ArrayExpress, R2 Genomics Analysis and Visualization Platform (http://r2platform.com/exRNAQC/) and https://github.com/OncoRNALab/exRNAQC (Data availability and Code availability), supplying the research community with different output formats (from raw sequencing data for bioinformaticians to browsable data access for any researcher) to mine and analyze the exRNAQC data. Along with the data, we also provide consistent and standardized pre-analytics information to better interpret, compare, and reproduce our results. To this purpose, the transcriptomes are well annotated according to the Biospecimen Reporting for Improved Study Quality (BRISQ) guidelines^22, 23^ (Supplementary Table 5). Overall, these aspects make the exRNAQC study not only the largest, but also the most comprehensive sequencing-based evaluation of pre-analytical factors affecting extracellular transcriptomes so far.

Although all eight tested RNA purification kits are marketed for purification of exRNA from serum or plasma, unexpectedly large performance differences were observed for both small RNA and, to a greater extent, mRNA. With most exRNA kits specifically developed for microRNA quantification, it is not very surprising that the kit performance at miRNA level is more homogenous than at mRNA level. We clearly noted that the mRNA purification performance was linked to the biofluid input and eluate volume. More specifically, a higher biofluid input volume resulted in higher mRNA concentrations. This association did not hold true for miRNA, as exemplified by CCF1 and CCF4. Also, RNA purification kits with a large eluate volume typically showed a high RNA yield but low RNA concentration. For these kits, condensing the eluate volume prior to library preparation could potentially increase their overall performance. Kits with a high extraction efficiency did not always result in better RNA quantification results because of limited biofluid input volumes. If these kits would accommodate a larger biofluid input volume (while maintaining their extraction efficiency), their overall performance could improve dramatically. Note, however, that the extraction efficiency of some kits decreased when using the maximum input volume compared to the minimum (e.g., CCF). Finally, we want to emphasize the importance of removing co-purified genomic DNA (gDNA) from the extracted RNA samples prior to exRNA quantification^24^. We observed high-level gDNA contamination in RNA-eluates produced with the MAP kit despite applying a commonly used gDNA removal strategy that worked well for all other RNA purification kits. This gDNA contamination is most likely due to an incompatibility between the RNA elution buffer and gDNA removal reagents. Alternative gDNA removal strategies should be evaluated and implemented before applying the MAP RNA extraction kit for exRNA analysis.

To evaluate the impact of the blood collection tube on downstream exRNA sequencing, biofluids (serum and plasma) were prepared at three different time intervals upon blood collection to assess potential changes in exRNA content due to blood storage at room temperature. To set a reference, each tube type was processed immediately after blood collection. For non-preservation tubes, we set the processing time intervals at 4 and 16 hours to mimic same-day processing and next-day processing, real-life situations often happening in the clinic. For preservation tubes that are specifically marketed to stabilize extracellular nucleic acids for 7 up to 14 days, more extreme time intervals for plasma preparation were selected, i.e., 24 and 72 hours upon blood collection. Surprisingly, in terms of stability over time, preservation tubes performed far worse than non-preservation tubes (including Serum), as reflected in increasing RNA concentrations and number of detected genes over time, and by compromised reproducibility. While preservation tubes were stored at room temperature for longer duration compared to non-preservation tubes, storage time was still substantially shorter than advertised for these tubes. In addition, RNA concentrations were much lower and hemolysis levels markedly higher in some of these tubes compared to non-preservation tubes, even at baseline (i.e., immediate processing upon blood draw). Although hemolysis may induce changes in exRNA content, the observed instability of the performance metrics over time for these tubes cannot solely be explained by differences in hemolysis over time. In this context, it is worth mentioning that, between individuals and across time intervals, we observed substantial differences in the amount of plasma that could be prepared from the preservation tubes, an issue that was reported prevously^25^. This also points towards performance instability (over time). Moreover, for DNA Streck and RNA Streck, library preparation resulted in insufficient yield for equimolar pooling and the fraction of reads mapping to the correct strand was lower compared to other tubes (see strandedness in https://github.com/OncoRNALab/exRNAQC). Apart from the performance metrics used to evaluate the different blood collection tubes, tube stability over time was further evaluated by analyzing circular versus linear RNA fractions, as well as by differential RNA abundance and originating cell type composition (via deconvolution) analyses. We hypothesized that the difference in stability between linear and circular RNA would translate to a distinct change in abundance over time, but this could not be confirmed. However, both the differential abundance analyses (across tubes and across time intervals) and the deconvolution analyses underscore the previous observation that preservation tubes perform far worse than non-preservation tubes. Based on these findings, we conclude that the tested preservation tubes are not suitable for exRNA analysis at the examined time intervals. It should be noted that during the exRNAQC study course, additional blood preservation tubes were developed and marketed, including the cfDNA/cfRNA Preservation Blood Tube (Zymo Research, R1075), the RNA Complete BCT (Streck, 230579), and cf-DNA/cf-RNA Preservative Tube (Norgen Biotek Corp., 63980). Posthoc evaluation using small RNA (for RNA Complete BCT) or mRNA capture sequencing (for cf-DNA/cf-RNA Preservative Tube) demonstrated poor performance (considering all evaluated performance metrics). The Zymo Research tube was not evaluated, as it could not be delivered by the company due to long delays in tube manufacturing (email communication). Importantly, although non-preservation tubes show better performance, it should be highlighted that the differential abundance analyses over time demonstrate that also for these tubes, the abundance of a considerable number of mRNAs changes over time. Based on the GSEA and deconvolution results, we recommend Citrate tubes for extracellular analysis and to process tubes within four hours upon blood draw for the analysis of exRNA. We invite blood collection tube manufacturers to increase their efforts to develop a plasma or serum tube that preserves the extracellular transcriptome for at least three days.

In the second phase of the exRNAQC study, we assessed whether interactions occur between pre-analytical variables such as RNA purification method, blood collection tube, and time interval. For both mRNA capture and small RNA sequencing, several two-way interactions between the blood collection tube and the RNA purification method, and between the blood collection tube and the time interval, were observed. In line with our expectations, no significant RNA purification method-time interval interactions were identified. Remarkably, for small RNA sequencing, more significant interactions affecting the performance metrics were detected compared to mRNA capture sequencing. We observed significant tube-kit interactions for the duplication rate and sensitivity performance metrics for the mRNA profiles, and for the extraction efficiency, sensitivity, reproducibility, and biotype metrics for the small RNA profiles. These interactions are not unexpected, given that blood collection tube coatings/preservatives/anticoagulants may induce changes in RNA purification conditions that impact performance, such as specific monovalent or divalent ions, general salt concentrations or pH values^26^. Importantly, the presence of these interactions demonstrates that one should not simply combine the best-performing blood collection tubes and RNA purification kits (from single factor studies), but that compatibility between these pre-analytical variables should be tested when optimizing a specific sample processing workflow. For both small RNA and mRNA sequencing, significant tube-time interval interactions were observed, confirming that even for the best-performing tubes, standardization of blood processing time intervals remain crucially important. Note that for mRNA profiles, results of Sequin-based performance metrics (i.e., RNA concentration and extraction efficiency) should be interpreted with caution (in exRNAQC phase 2), as we cannot exclude differential extraction efficiencies of Sequin spike-in RNA between MIR0.2 and CCF2. Importantly, this finding does not impact our conclusions of exRNAQC phase 1, given that Sequins were only used for performance evaluation of the different blood collection tubes for which exRNA was extracted using the same RNA purification method (i.e., MIR0.2). Also, the performance metrics for evaluation of the different RNA purification methods in exRNAQC phase 1 do not rely on Sequin spike-ins.

Although the exRNAQC study represents the most comprehensive performance assessment of RNA purification methods and blood collection tubes in the context of exRNA profiling to date, the study also comes with a few limitations. All experiments were performed in a single laboratory. Ideally, a multicenter study should confirm the present findings. Although liquid biopsy collection and processing procedures in the exRNAQC study were tightly controlled, we cannot exclude a potential impact of other pre-analytics, e.g. fasting status of the donor or biofluid storage conditions. In addition, different donors were sampled for evaluation of non-preservation and preservation blood collection tubes. As such, absolute values of the performance metrics (Supplementary Fig. 7 and Supplementary Fig. 8) cannot be directly compared across these two groups of tubes. However, note that this does not impact our conclusions for preservation tubes, as these are based on the stability of the performance metrics over time, i.e. comparing performance of the same tube type at different time intervals. Also, in the cellular deconvolution analysis, we did not include red blood cells and platelets, since the EPIC deconvolution tool has no signatures for these blood fractions. Therefore, blood collection tube performance in terms of platelet activation mediated exRNA release was not studied in the exRNAQC study. Nevertheless, the GSEA results of some tubes, including Serum and EDTA (Supplementary Table 4), point towards the presence of platelet activation. Note that the performance metrics solely assess technical performance, and that the impact of the pre-analytics on biomarker detection was not addressed in this study. Finally, we only focused on the analysis of microRNAs for small RNA sequencing. Although important, the study of other types of small RNAs was beyond the scope of this study.

In the exRNAQC study, we demonstrate that the choice of RNA purification method and blood collection tube substantially impacts mRNA and miRNA quantification by evaluation of 11 performance metrics. Here, eight commercially available RNA purification methods and ten blood collection tubes were studied, but the proposed framework and metrics can also be used to evaluate the performance of more recently developed RNA purification methods and blood collection tubes. Based on the findings presented here, we highly recommend a) standardizing sample collection and processing, b) carefully annotating and reporting pre-analytics, and c) making use of synthetic spike-in RNA molecules for sequencing-based quality control and optional normalization of exRNA. This is crucially important for interpretation and comparison of all exRNA study results and will enhance the reproducibility of exRNA research as a starting point for biofluid based biomarker studies.

## Data availability

Full access to the data of the exRNAQC study is available through the European Genome-phenome Archive (EGA; study ID EGAS00001005263 (exRNAQC phase1) and EGAS00001006499 (exRNAQC phase 2) and ArrayExpress (accession ID E-MTAB-10504 - E-MTAB-10507). mRNA capture and small RNA sequencing of the RNA purification kit experiment (exRNAQC phase 1) were identified with study codes exRNAQC004 and exRNAQC011, respectively. mRNA capture and small RNA sequencing of the blood collection tube experiment (exRNAQC phase 1) were identified with study codes exRNAQC005 and exRNAQC013, respectively. mRNA capture and small RNA sequencing of exRNAQC phase 2 were identified with the study code exRNAQC017. Browsable access is provided through the R2 Genomics Analysis and Visualization Platform (http://r2platform.com/exRNAQC/), as exemplified for the analysis of differential gene abundance across time intervals in Supplementary Fig. 16. Note that in R2, data on MAP RNA eluates were excluded (because of residual DNA contamination), and that data can be analyzed using two different normalization strategies, i.e., counts normalized by variance stabilizing transformation (DESeq2) or spike-normalized counts. Detailed sample annotation (pre-analytics information) can be found in Supplementary Table 5.

## Code availability

Analysis pipelines are available through GitHub (https://github.com/OncoRNALab/exRNAQC).

## Abbreviations

ACD-A: BD Vacutainer Glass ACD Solution A tube

ALC: area left of the curve

Biomatrica: LBgard Blood Tube

BRISQ: Biospecimen Reporting for Improved Study Quality

bp: base pair

CCF: QIAamp ccfDNA/RNA Kit

cfRNA: cell-free RNA

CIRC: Plasma/Serum Circulating and Exosomal RNA Purification Kit/Slurry Format

circRNA: circular RNA

Citrate: Vacuette Tube 9 ml 9NC Coagulation sodium citrate 3.2%

DNA Streck: Cell-Free DNA BCT

EDTA: BD Vacutainer Plastic K2EDTA tube

EDTA separator: Vacuette Tube 8 ml K2E K2EDTA Separator

EGA: European Phenome-Genome Archive

ERCC: Extracellular RNA Communication Consortium

exRNA: extracellular RNA

FC: fold change

gDNA: genomic DNA

LP: Library Prep Control

MAP: MagNA Pure 24 Total NA Isolation Kit in combination with the MagNA Pure instrument

MAX: Maxwell RSC miRNA Plasma and Exosome Kit in combination with the Maxwell RSC Instrument

MIR: miRNeasy Serum/Plasma Kit

MIRA: miRNeasy Serum/Plasma Advanced Kit

miRNA: microRNA

MIRV: mirVana PARIS Kit with purification protocol for total RNA

MIRVE: mirVana PARIS Kit with purification protocol for RNA enriched for small RNAs

mRNA: messenger RNA

NUC: NucleoSpin miRNA Plasma Kit

PAXgene: PAXgene Blood ccfDNA Tube

RA3: RNA 3’ adapter

RA5: RNA 5’ adapter

RC: RNA extraction Control

RNA Streck: Cell-Free RNA BCT

Roche: Cell-Free DNA Collection Tube

Serum: BD Vacutainer SST II Advance Tube

SOP: standard operating procedure.

## Methods

### Donor material and liquid biopsy preparation

Sample collection was approved by the ethics committee of Ghent University Hospital (Belgian Registration number B670201733701) and written informed consent was obtained from 20 healthy donors, including 5 males and 15 females (age ranges from 27 to 54 years old). Incapacitated or pregnant individuals, as well as individuals younger than 20 years old were excluded from the study. Venous blood was collected from an elbow vein after disinfection with 2% chlorhexidine in 70% alcohol. In total, ten different blood collection tubes were used: the BD Vacutainer SST II Advance Tube (referred to as Serum in this study; Becton Dickinson and Company, 366444), BD Vacutainer Plastic K2EDTA tube (EDTA; Becton Dickinson and Company, 367525), Vacuette Tube 8 ml K2E K2EDTA Separator (EDTA separator; Greiner Bio-One, 455040), BD Vacutainer Glass ACD Solution A tube (ACD-A; Becton Dickinson and Company, 366645), Vacuette Tube 9 ml 9NC Coagulation sodium citrate 3.2% (citrate; Greiner Bio-One, 455322), Cell-Free RNA BCT (RNA Streck; Streck, 230248), Cell-Free DNA BCT (DNA Streck; Streck, 218996), PAXgene Blood ccfDNA Tube (PAXgene; Qiagen, 768115), Cell-Free DNA Collection Tube (Roche; Roche, 07785666001), and LBgard Blood Tube (Biomatrica; Biomatrica, M68021-001). Immediately after blood draw, blood collection tubes were inverted five times and all tubes were transported to the laboratory for plasma or serum preparation. Tubes were immediately processed or at 4h, 16h, 24h or 72h upon blood collection. Details on the different blood draws and plasma/serum preparations are available in the Supplementary Information.

### RNA isolation and gDNA removal

In total, eight different exRNA purification methods, including six spin column-based kits and two automated extraction procedures, were used according to the manufacturer’s manual: the miRNeasy Serum/Plasma Kit (abbreviated to MIR in this study; Qiagen, 217184), miRNeasy Serum/Plasma Advanced Kit (MIRA; Qiagen, 217204), mirVana PARIS Kit (MIRV; Life Technologies, AM1556), NucleoSpin miRNA Plasma Kit (NUC; Macherey-Nagel, 740981.50), QIAamp ccfDNA/RNA Kit (CCF; Qiagen, 55184), Plasma/Serum Circulating and Exosomal RNA Purification Kit/Slurry Format (CIRC; Norgen Biotek Corp., 42800), Maxwell RSC miRNA Plasma and Serum Kit (Promega, AX5740 and AS1680) in combination with the Maxwell RSC Instrument (MAX; Promega, AS4500), and MagNA Pure 24 Total NA Isolation Kit (Roche, 07658036001) in combination with the MagNA Pure 24 instrument (MAP; Roche, 07290519001). Per 100 µl liquid biopsy input volume, 1 µl Sequin spike-in controls (Garvan Institute of Medical Research^27^) and/or 1 µl RNA extraction Control (RC) spike-ins^28^ (IDT) were added to the lysate for TruSeq RNA Exome Library Prep sequencing and/or TruSeq Small RNA Library Prep sequencing, respectively (see Supplementary Information for concentrations). To maximally concentrate the RNA eluate, minimum eluate volumes were used, unless otherwise recommended by the manufacturer. For evaluation of the different purification methods in exRNAQC phase 1, both the minimum and maximum recommended plasma input volumes were tested in triplicate. Details on the exRNA purification methods, and Sequin and RC spike-in controls are available in the Supplementary Information.

gDNA removal of RNA samples for TruSeq RNA Exome Library Prep sequencing was performed using HL-dsDNase (ArcticZymes, 70800-202) and Heat & Run 10X Reaction Buffer (ArcticZymes, 66001). Briefly, 2 µl External RNA Control Consortium (ERCC) spike-in controls (ThermoFisher Scientific, 4456740), 1 µl HL-dsDNase and 1.4 µl reaction buffer were added to 12 µl RNA eluate, and incubated for 10 min at 37 °C, followed by 5 min at 55 °C. To RNA samples used for both TruSeq RNA Exome Library Prep sequencing and TruSeq Small RNA Library Prep sequencing, also 2 µl Library Prep Control (LP) spike-ins^29^ (IDT) were added to the RNA eluate before starting gDNA removal and 1.6 µl reaction buffer was used. RNA samples solely used for TruSeq Small RNA Library Prep sequencing were not DNase treated. Here, 2 µl LP spike-ins were added to 12 µl RNA eluate before starting library preparation. Details on ERCC and LP spike-in control concentrations are available in the Supplementary Information.

### mRNA capture sequencing

mRNA libraries were prepared starting from 8.5 µl RNA eluate using the TruSeq RNA Exome Kit (Illumina, 20020189, 20020490, 20020492, 20020493, 20020183), according to the manufacturer’s protocol with following adaptations: fragmentation of RNA for 2 min at 94 °C, second strand cDNA synthesis for 30 minutes at 16 °C (with the thermal cycler lid pre-heated at 40 °C), and second PCR amplification using 14 PCR cycles. Upon the first and second PCR amplification, libraries were validated on a Fragment Analyzer (Advanced Analytical Technologies), using 1 µl of library. Library concentrations were determined using Fragment Analyzer software for smear analysis in the 160 to 700 base pair (bp) range. Library quantification was qPCR-based, using the KAPA Library Quantification Kit (Kapa Biosystems), and/or based on NanoDrop 1000 measurements. Further details on the library preparation and quantification protocol are described in Hulstaert *et al.*^21^ For evaluation of the different RNA purification methods, 45 libraries were pooled on replicate level at 4 nM, yielding three pools of 15 samples, quality controlled using the KAPA Library Quantification Kit, and sequenced on a NextSeq 500 instrument (NextSeq 500/550 High Output Kit v2.5 (Illumina, 20024907, PE 2 x 75 cycles)). Loading concentrations of the three pools ranged from 2.1 pM to 2.3 pM. Percentage PhiX was 3%. For evaluation of the different blood collection tubes, all 90 libraries were pooled at 1.5 nM or the highest possible concentration, quality controlled using the KAPA Library Quantification Kit, and sequenced on a NovaSeq 6000 instrument (NovaSeq 6000 S2 Reagent Kit (Illumina, 20012861, PE 2 x 75 cycles)). Loading concentration of the pool was 324 pM. Percentage PhiX was 1 %. For exRNAQC phase 2, 90 libraries were pooled at 5.50 nM, quality controlled using the KAPA Library Quantification Kit, and sequenced on a SP100 flow cell (Illumina, NovaSeq 6000). Loading concentration of the pool is 340 pM. Differences in read distribution across samples were subsequently used to re-pool individual libraries in order to obtain an equimolar pool. Subsequently, samples were sequenced on a S2 flow cell, at a loading concentration of 360 pM.

### Small RNA sequencing

Small RNA libraries were prepared starting from 5 µL RNA eluate using the TruSeq Small RNA Library Prep Kit (Illumina, RS-200-0012, RS-200-0024, RS-200-0036, RS-200-0048), according to the manufacturer’s protocol with following adaptations: the RNA 3’ adapter (RA3) and the RNA 5’ adapter (RA5) were 4-fold diluted with RNase-free water, and the number of PCR cycles was increased to 16^20, 30^. For phase 1, samples were divided across library prep batches according to index availability. For each batch, 3 µl of small RNA library from each sample was pooled prior to automated size selection using the Pippin prep (Sage Sciences, CDH3050). Size selected libraries were quantified using qPCR, and sequenced on a MO flow cell (Illumina, NextSeq 500) using loading concentrations ranging from 1.2 to 2.4 pM. Differences in read distribution across samples were subsequently used to re-pool individual libraries in order to obtain an equimolar pool. After size selection on a Pippin prep and qPCR quantification, these pools were sequenced on a HO flow cell (Illumina, NextSeq 500, NextSeq 500/550 High Output Kit v2.5, 20024907) using loading concentrations ranging from 1.2 to 3 pM. For phase 2, individual libraries were quantified using qPCR and pooled equimolarly across 2 pools. After size selection on a Pippin prep and qPCR quantification, library pools were sequenced on a NovaSeq 6000 XP flow cell (Illumina) using a loading concentration of 270 nM.

### Data analysis

In total, 456 transcriptomes were profiled and analyzed. The raw, processed and metadata were submitted to the European Genome-phenome Archive (EGA), ArrayExpress and R2 Genomics Analysis and Visualization Platform (see Data availability). A high-level summary of the sequencing statistics can be found in Supplementary Table 3 and Supplementary Table 6-10. Detailed pre-analytics information (for the BRISQ elements^22, 23^) can be found in Supplementary Table 5.

### Quality control and quantification of mRNA capture sequencing data

In case of adapter contamination indicated by FASTQC (v0.11.8; www.bioinformatics.babraham.ac.uk/projects/fastqc), adapters were trimmed with Cutadapt^31^ (v1.18; 3’ adapter R1: ‘AGATCGGAAGAGCACACGTCTGAACTCCAGTCA’; 3’ adapter R2 ‘AGATCGGAAGAGCGTCGTGTAGGGAAAGAGTGT’). Only reads with ≥ 99% accuracy in at least 80% of bases of both mates were kept. Subsequently, FASTQ files were subsampled with Seqtk (v1.3; https://github.com/lh3/seqtk) to the lowest number of reads pairs obtained in the experiment. Since the low amount of input RNA resulted in a high number of duplicates (Supplementary Table 3 and Supplementary Table 7), we removed these duplicates using Clumpify dedupe (v38.26; www.sourceforge.net/projects/bbmap) with the following specifications: paired-end mode, 2 substitutions allowed, kmersize of 31, and 20 passes. For duplicate removal, only the first 60 bases of both reads were considered to account for the sequencing quality drop at the end of the reads. Strand-specific transcript-level quantification of the deduplicated FASTQ files was performed with Kallisto^32^ (v0.44.0). For coverage and strandedness analysis, mapped reads were obtained by STAR^33^ (v2.6.0c) using the default parameters (except for --twopassMode Basic, --outFilterMatchNmin 20 and -- outSAMprimaryFlag AllBestScore). For all exons coverage information was retrieved by the genomeCoverageBed and intersectBed functions of BEDTools^34^ (v2.27.1). Strandedness information was obtained with RSeQC^35^ (v2.6.4). The reference files for all analyses were based on genome build hg38 (www.ncbi.nlm.nih.gov/assembly/GCF_000001405.26) and transcriptome build Ensembl v91^36^. Spike annotations were added to both genome and transcriptome files.

### Quality control and quantification of small RNA sequencing data

First, adaptor trimming (3’ adapter: TGGAATTCTCGGGTGCCAAGG) was performed using Cutadapt (v1.16) with a maximum error rate of 0.15 and discarding reads shorter than 15 bp and those in which no adaptor was found. Subsequently, low quality reads were filtered out (reads with ≥ Q20 in at least 80% of bases were kept) by FASTX-Toolkit (v0.0.14; http://hannonlab.cshl.edu/fastx_toolkit/index.html). Filtered FASTQ files were subsampled to the minimum number of reads in the experiment (Supplementary Table 6 and Supplementary Table 8) using Seqtk (v1.3). Reads were collapsed with FASTX-Toolkit and LP and RC spike reads (including possible fragments) were annotated. The non-spike reads were mapped with Bowtie^37^ (v1.2.2, with additional parameters -k 10 -n 1 -l 25) considering only perfect matches. Mapped reads were annotated by matching the genomic coordinates of each read with genomic locations of miRNAs (obtained from miRBase^38–43^, v22) and other small RNAs (tRNAs obtained from UCSC GRCh38/hg38; snoRNA, snRNA, MT_tRNA, MT_rRNA, rRNA, and miscRNA from Ensembl, v91).

### Defining performance metrics

The statistical programming language R (v4.0.3; www.r-project.org) was used throughout this section and all scripts can be found at GitHub (https://github.com/OncoRNALab/exRNAQC). In total, 11 performance metrics were developed, of which nine were used for evaluation of the different RNA purification methods (exRNAQC phase 1), five for blood collection tube evaluation and six for phase 2 interaction analyses (Table 1). Each performance metric is briefly explained below. For each part of the study, more in-depth descriptions of the metrics and results are available through GitHub.

### Count threshold

To distinguish signal from noise, we made use of pairwise count comparisons across three technical replicates for evaluation of the different RNA purification methods. We defined a count threshold for each RNA purification method and biotype in a similar manner as defined in the miRQC study^17^. Specifically, a threshold that reduces the fraction of single positives in technical replicates by at least 95 % (single positives are cases where a given gene has zero counts in one replicate and a non-zero value in the other one). This threshold can be used as a reproducibility metric between technical replicates. For each kit-volume combination, the median threshold of the three pairwise replicate comparisons was used (Supplementary Table 2). As the blood collection tube experiment in exRNAQC phase 1 did not have technical replicates and RNA purification for all tubes was performed using MIR0.2, the median thresholds of MIR0.2 (3 counts for small RNAs; 6 counts for mRNAs) were applied here as well.

### Data retention

Data retention is defined as the percentage of gene counts remaining after applying the count threshold as filter, therefore giving information about the fraction of counts lost by applying the cut-off.

### Sensitivity

We defined sensitivity as the number of different protein coding genes or miRNAs picked up above the count threshold.

### RNA concentration

For the RNA purification method experiment in exRNAQC phase 1, the different methods were tested on the same plasma sample. By adding equal amounts of ERCC and LP spikes (for mRNA and small RNA, respectively) after RNA purification, we were able to calculate endogenous RNA concentrations in the eluate. For instance, in cases of low endogenous RNA content after RNA purification, relatively more ERCC and LP spikes will be sequenced. By dividing the total sum of endogenous counts by the sum of ERCC or LP spikes, we could therefore compare the RNA concentrations in the eluate of the different purification methods. For the blood collection tube experiment of phase 1 and for the blood collection tube-kit combinations of phase 2, we were interested in the impact of the different blood collection tubes on the RNA concentration in plasma. By adding equal amounts of Sequin and RC spikes (for mRNA and small RNA, respectively) during RNA purification (upon sample lysis), we were able to calculate relative endogenous RNA concentrations in the plasma. For instance, in cases of low endogenous RNA content before extraction, relatively more Sequin and RC spikes will be sequenced. By dividing the total sum of endogenous counts by the sum of Sequin or RC spikes, we could therefore compare the RNA concentrations in plasma of the different tubes.

### RNA yield

Multiplying the RNA concentration by the eluate volume gives the RNA yield in the total eluate.

### Extraction efficiency

Correcting the relative RNA yield for the plasma input volume (dividing yield by input volume) gives an idea of the theoretical RNA extraction efficiency of the method.

### Reproducibility

As described in the miRQC study^17^, the area left of the cumulative distribution curve (ALC) was calculated by comparing the actual cumulative distribution curve of log2 fold changes in gene or miRNA abundance between pairs of replicates to the theoretical cumulative distribution (optimal curve). Less reproducibility between samples results in more deviations from this optimal curve and therefore larger ALC-values.

### Duplication rate

Duplication rate was obtained by dividing the number of reads removed by Clumpify (see Methods) by the number of subsampled reads.

### Coverage

Coverage is the percentage of bases from the total transcriptome covered by at least one sequencing read.

### Hemolysis

Hemolysis was measured with Nanodrop (absorbance of light at 414 nm) in plasma.

### Biotype

Fraction of total counts that go to mRNA (mRNA capture sequencing) or miRNAs (small RNA sequencing).

### Accounting for size selection bias

For the small RNA library preparation of the RNA purified using the different methods in exRNAQC phase 1, the three technical replicates of each extraction method were divided over three different pools. Next, pippin prep size selection for miRNAs occurred on each pool individually. To account for size selection bias (which resulted in consistently lower sequencing counts in the second pool), we each time downsampled the miRNA counts of the other two replicates to the sum of miRNA counts of the replicate in the second pool. Down-sampling was based on reservoir sampling - random sampling without replacement (subsample_miRs.py script on https://github.com/OncoRNALab/exRNAQC).

### Transforming performance metrics into robust z-scores

For evaluation of the different RNA purification methods in exRNAQC phase 1, individual scores for performance metrics were transformed to z-scores. As the standard z-score is sensitive to outliers, we used a robust z-score transformation, based on the median (μ_1/2_) and median absolute deviation (*MAD = median_i_(|X_i_ − median X_1…n_|)*), instead. The general formula for robust z-score calculation is shown below:

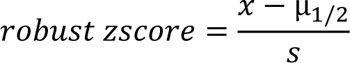

Where s is a scaling factor that depends on the MAD. In case MAD is not zero: s = MAD ∗ 1.4826. If MAD equals zero, s approximately equals the standard deviation: s = meanAD ∗ 1.2533, with meanAD = mean_i_(|X_i_ − mean X_1…n_|) (https://asq.org/quality-press/display-item?item=E0801, https://www.ibm.com/docs/en/cognos-analytics/11.1.0?topic=terms-modified-z-score).

### Fold change analyses for stability over time assessment

To evaluate tube stability across time intervals in exRNAQC phase 1 and 2, we determined several performance metrics per blood collection tube at different time intervals. We then calculated, for every tube and donor, the fold chance across different time intervals (relative to the base interval at T0, so excluding T24-72 and T04-16). A theoretical example is shown in Supplementary Fig. 6.

### circRNA and linear RNA fraction determination

For the assessment of blood collection tube stability over time in exRNAQC phase 1, an in-house pipeline was used to investigate the differences in fractions of circRNAs between tubes and time intervals. Starting from the raw FASTQ files from the mRNA capture sequencing, Cutadapt^31^ (v1.18) was used to remove the adapter sequences and reads that end up shorter than 20 bp. Next, the reads of which less than 80% of the bases had a Q-score higher than 19 were removed. Subsequently, clumpify.sh from BBMap (v38.26; (https://sourceforge.net/projects/bbmap) with default parameters was used to remove PCR duplicate reads. The deduplicated reads were mapped using TopHat^44^ (v2.1.0) with Bowtie (v1.1.2) and fusion mapping turned on. Next, the CIRCexplorer2^45^ (v2.3.3) functions parse, annotate, assemble and denovo were used to identify and annotate known circRNAs and to identify novel circRNAs or alternative back-splicing events. Last, the circRNA ratios on back-splice junction and gene level were calculated using CiLiQuant^46^ (v1.0).

### Differential abundance analyses

Differential abundance analyses were performed on the data of the blood collection tube experiment of exRNAQC phase 1. In a matrix selected for T0 samples, genes were filtered out when not present with a minimum of 10 counts in all three replicates of one tube type. For the comparison of the time intervals, only the samples for the respective tube were selected and genes were filtered out when not present with a minimum of 10 counts in all three replicates of one time interval. The filtered data was normalized with Limma voom (v3.52.4) and contrasts, comparing subsequent time intervals to T0, were fit. Genes with a |log 2 FC| > 1 and an Benjamini-Hochberg adjusted p-value < 0.05 were retained as significant. On the log2 fold change ranked gene list, gene set enrichment analysis (GSEA) with fgsea (v1.22.0) on the MSigDB C2 pathways was performed. Pathways with an Benjamini-Hochberg adjusted p-value of less than 0.05 were retained as significantly up or down regulated.

### Differences in immune cell composition over time

To further evaluate blood collection tube stability over time in exRNAQC phase 1, we first we used computational deconvolution (on subsampled data) to infer the cell type composition (proportions) of different immune cell types present in blood^47^. Since the origin (niche) of the expression profiles has a tremendous impact on the deconvolution results^48^ and it is possible that RNA coming from other cell type(s) is also present in circulation, we used EPIC^49^, a method that has a built-in reference matrix from circulating immune cells (known as ‘BRef’ signature) and includes the presence of an unknown component (‘otherCells’). Specifically, we used TPM normalized count matrices as input, as recommended by the authors^49^ and shown as the optimal choice for this method in a recent benchmarking study^50^.

Next, to evaluate differences in cell type composition of several blood immune cell types, we performed a repeated-measures analysis by means of beta regression models with random effects. For each cell type a separate model was fitted with ‘tube’ and ‘time interval’ as factor variables (main and interaction effects included), with donor as random effect and with tube-specific variance components (allowing for variance heterogeneity that was observed in the data exploration phase). All models were fit with the glmmTMB R package (v1.1.2.3)^51^. Based on the model fits, all pairwise comparisons between the three time intervals were tested for each type of tube: T0 vs T1, T0 vs T2 and T1 vs T2. For each combination of cell type and tube, the p-values were adjusted for multiple testing with Tukey’s method as implemented in the emmeans.glmmTMB R function (R packages glmmTMB and emmeans (v1.7.0; https://CRAN.R-project.org/package=emmeans)). All analyses were done with the statistical software R (v4.1.0; www.r-project.org). See https://github.com/OncoRNALab/exRNAQC (exRNAQC005, deconvolution) for a detailed report with the corresponding R code.

### Repeated measures analyses

For data analysis of exRNAQC phase 2, linear mixed-effects models were built with the nlme package (v3.1-157) in R. Blood collection tube, RNA purification method and time interval were included as fixed effects and donor ID as random effect. The heteroscedasticity introduced by different RNA purification methods was considered. Next, an ANOVA test was performed on the model to estimate the significance of the interactions. The normality of the residuals was checked with the qqnorm function (see https://github.com/OncoRNALab/exRNAQC).

## Acknowledgements

This study was in part funded by Ghent University (BOF-GOA), Stand up to Cancer (Kom op tegen Kanker, the Flemish cancer society), the Foundation against Cancer, Research Foundation Flanders (FWO), the Liquidhope Transcan-2 project and the European Union’s Horizon 2020 research and innovation program (grant agreement 826121). H.H.H., C.E., J.D.W., E.H., P.D., and R.V.P. were funded by a predoctoral fellowship grant from the FWO (12Q6217N, 1S07416N, 1S90621N, 1133120N,11H7523N, and 11B3718N). F.A.C was supported by a Special Research Fund postdoctoral scholarship from Ghent University (BOF21/PDO/007). A.M. was supported by a Special Research Fund scholarship from Ghent University, Stand up to Cancer and a predoctoral fellowship grant from the FWO (11C1621N). A.D. was supported by a postdoctoral fellowship grant from the Special Research Fund of Ghent University and the FWO (1224021N). K.S. was supported by a grant from Stand up to Cancer. J.D. was supported by the Special Research Fund of Ghent University. We thank Illumina for sponsoring library preparation and sequencing reagents, and Qiagen, Promega and Roche for sponsoring blood collection tubes and/or RNA purification kits. The sponsors had no role in the design, interpretation, or writing of the study.

## Author contributions

The exRNAQC Consortium

Author contributions are reported according to the CRediT taxonomy^52^. Within the CRediT groups, authors are in alphabetical order and * indicates lead contributions. All authors approved the manuscript.

## Conceptualization

Anneleen Decock^1,2^, Olivier De Wever^2,3^, Celine Everaert^1,2,4^, Hetty Hilde Helsmoortel^1,2^, An Hendrix^2,3,^ Pieter Mestdagh^1,2,5^, Annelien Morlion^1,2^, Jo Vandesompele^1,2,5^ & Ruben Van _Paemel_^1^_,2,4,6_

## Data curation

Francisco Avila Cobos^1,2,4^, Anneleen Decock^1,2^, Celine Everaert^1,2,4^, Hetty Hilde Helsmoortel^1,2^, Jan Koster^7^, Annelien Morlion^1,2^, Franco Poma-Soto^1,2^, Ruben Van Paemel^1,2,4,6^ & Jasper Verwilt^1,2^

## Formal analysis

Jasper Anckaert^1,2^, Francisco Avila Cobos^1,2,4^, Jilke De Wilde^1,2,4*^, Celine Everaert^1,2,4^, Carolina Fierro^5^, Pieter Mestdagh^1,2,5^, Annelien Morlion^1,2,*^, Franco Poma-Soto^1,2^, Jo Vandesompele^1,2,5^, Ruben Van Paemel^1,2,4,6,*^ & Jasper Verwilt^1,2^

## Funding acquisition

Anneleen Decock^1,2^, Pieter Mestdagh^1,2,5^ & Jo Vandesompele^1,2,5^

## Investigation

Anneleen Decock^1,2^, Philippe Decruyenaere^1,2,8^, Jill Deleu^1,2^, Jilke De Wilde^1,2,4^, Bert Dhondt^2,3,9,^ Hetty Hilde Helsmoortel^1,2^, Eva Hulstaert^1,2,10^, Nele Nijs^5^, Justine Nuytens^1,2^, Annouck Philippron^2,11,12^, Kathleen Schoofs^1,2,4^, Eveline Vanden Eynde^1,2^, Ruben Van Paemel^1,2,4,6,^ Kimberly Verniers^1,2^, Jasper Verwilt^1,2^ & Nurten Yigit^1,2^

## Methodology

Francisco Avila Cobos^1,2,4^, Anneleen Decock^1,2^, Bert Dhondt^2,3,9,^ Thibaut D’huyvetter^1,2^, Celine Everaert^1,2,4,^ Carolina Fierro^5^, Hetty Hilde Helsmoortel^1,2^, Pieter Mestdagh^1,2,5^, Annelien Morlion^1,2^, Nele Nijs^5^, Annouck Philippron^2,11,12,^ Thomas Piofczyk^5^, Olivier Thas^13,14,15^, Jo Vandesompele^1,2,5^ & Ruben Van Paemel^1,2,4,6^

## Project administration

Anneleen Decock^1,2^

## Resources

Philippe Decruyenaere^1,2,8^, Jilke De Wilde^1,2,4^, Bert Dhondt^2,3,9^, Eva Hulstaert^1,2,10^, Jan Koster^7^, Scott Kuersten^16^, Tim R. Mercer^17,18^, Annouck Philippron^2,11,12^, Gary P. Schroth^16^ & Ruben Van Paemel^1,2,4,6^

## Software

Jasper Anckaert^1,2^, Francisco Avila Cobos^1,2,4,^ Celine Everaert^1,2,4^, Annelien Morlion^1,2^, Olivier Thas^13,14,15,^ Ruben Van Paemel^1,2,4,6^ & Jasper Verwilt^1,2^

## Supervision

Katleen De Preter^2,4^, Pieter Mestdagh^1,2,5^, Jo Vandesompele^1,2,5^ & Tom Van Maerken^1,2,19^

## Visualization

Francisco Avila Cobos^1,2,4^, Anneleen Decock^1,2^, Jilke De Wilde^1,2,4^, Celine Everaert^1,2,4^, Jan Koster^7^, Annelien Morlion^1,2,^ Franco Poma-Soto^1,2^, Kathleen Schoofs^1,2,4^, Ruben Van Paemel^1,2,4,6^ & Jasper Verwilt^1,2^

## Writing - original draft

Francisco Avila Cobos^1,2,4^, Anneleen Decock^1,2^, Philippe Decruyenaere^1,2,8^, Jill Deleu^1,2^, Jilke De Wilde^1,2,4^, Celine Everaert^1,2,4^, Annelien Morlion^1,2^, Kathleen Schoofs^1,2,4^, Ruben Van Paemel^1,2,4,6^ & Jasper Verwilt^1,2^

## Writing - review & editing

Anneleen Decock^1,2^, Celine Everaert^1,2,4,^ Pieter Mestdagh^1,2,5,^ Annelien Morlion^1,2^ & Jo Vandesompele^1,2,5^

## Affiliations

1. OncoRNALab, Center for Medical Genetics, Department of Biomolecular Medicine, Ghent University, Ghent, Belgium.
2. Cancer Research Institute Ghent (CRIG), Ghent, Belgium.
3. Laboratory of Experimental Cancer Research, Ghent University, Ghent, Belgium.
4. TOBI lab, Center for Medical Genetics, Department of Biomolecular Medicine, Ghent University, Ghent, Belgium.
5. Biogazelle, Zwijnaarde, Belgium.
6. Department of Pediatrics, Ghent University Hospital, Ghent, Belgium.
7. Center for Experimental and Molecular Medicine (CEMM), Amsterdam University Medical Centers, University of Amsterdam, Amsterdam, The Netherlands.
8. Department of Hematology, Ghent University Hospital, Ghent, Belgium. 9 Department of Urology, Ghent University Hospital, Ghent, Belgium.
9. Department of Dermatology, Ghent University Hospital, Ghent, Belgium. 11 Lab for Experimental Surgery, Ghent University, Ghent, Belgium.
10. Department of Gastrointestinal Surgery, Ghent University Hospital, Ghent, Belgium.
11. Data Science Institute (DSI), Interuniversity Institute for Biostatistics and Statistical Bioinformatics (I-BioStat), Hasselt University, Hasselt, Belgium.
12. Department of Applied Mathematics, Computer Science and Statistics, Ghent University, Gent, Belgium.
13. National Institute of Applied Statistics Research Australia (NIASRA), University of Wollongong, Wollongong, Australia.
14. Illumina, San Diego, California, USA.
15. Australian Institute of Bioengineering and Nanotechnology, University of Queensland, Brisbane, QLD, Australia.
16. Kinghorn Centre for Clinical Genomics, Garvan Institute of Medical Research, Sydney, NSW, Australia.
17. Department of Laboratory Medicine, AZ Groeninge, Kortrijk, Belgium.

## Conflict of interest

Carolina Fierro and Nele Nijs are employees, Thomas Piofczyk is a former employee, Pieter Mestdagh is a consultant, and Jo Vandesompele a co-founder of Biogazelle, a clinical CRO providing human biofluid extracellular RNA sequencing, now a CellCarta company. Gary P. Schroth and Scott Kuersten are employees of Illumina, providing library preparation and sequencing reagents. Promega, Qiagen and Roche sponsored blood collection tubes and/or RNA purification kits. Funders did not influence data analysis, interpretation, and manuscript writing.

### Additional information

#### Supplementary Table legends

**Supplementary Table 1. Available literature on the influence of pre-analytics on RNA sequencing data, including studies on plasma and/or serum.** The pre-analytics analyzed in the selected studies are listed: number of blood collection tube types; hemolysis measured (yes/no); the fluid (serum/plasma or both); number of centrifugation protocols; number of RNA isolation kits; the RNA type; the gene expression analysis method; other pre-analytics.

**Supplementary Table 2. Filter threshold of the different RNA purification methods.** Kit: RNA purification kit abbreviation; mRNA threshold: median threshold that removes 95% of single positive genes between technical replicates; miRNA threshold: median threshold that removes 95% of single positive miRNAs between technical replicates. More explanation on these thresholds in Methods. NA: Not applicable.

**Supplementary Table 3. mRNA capture sequencing data statistics of RNA purification kit experiment (exRNAQC004).** UniqueID: RNA identifier; SampleID: combination of kit abbreviation and technical replicate number; raw_reads: number of sequenced reads pairs; qcfiltered_reads: number of read pairs after quality filtering; post_subsampling: number of read pairs after subsampling; post_deduplication: number of read pairs after Clumpify duplicate removal; duplicate_prct: % of duplicates in subsampled reads; kallisto_prct_alignment: % of duplicate removed reads that were pseudoaligned; strandedness_prct: % of reads on correct strand (stranded protocol).

**Supplementary Table 4. Gene set enrichment analysis results on differential abundant mRNAs across time intervals for each blood collection tube type of exRNAQC phase 1 and 2.** Results of the different comparisons are summarized in different tabs. For each comparison, gene sets significantly enriched for differential abundant genes are indicated, as well as their p-value (pval), BH-adjusted p-value (padj), expected error for the standard deviation of the p-value logarithm (log2err), normalized enrichment score (NES) and leading-edge genes that drive the enrichment.

**Supplementary Table 5. Pre-analytical variable annotation for all samples included in the exRNAQC study.** In the first tab, the different pre-analytical variables are listed, and for each of them a description is provided. Note that the pre-analytics are categorized into three groups, i.e., variables linked to the blood draw (with prefix B_), biofluid preparation (with prefix L_) or RNA purification (with prefix R_). This tab also includes a description of the BRISQ elements^22,23^. In the following tabs, annotated samples are listed per experiment (the mRNA capture sequencing of the RNA purification kit study (exRNAQC004), the mRNA capture sequencing of the blood collection tube study (exRNAQC005), the small RNA sequencing of the RNA purification kit study (exRNAQC011), the small RNA sequencing of the blood collection tube study (exRNAQC013) or the mRNA capture/small RNA sequencing of phase 2 (exRNAQC017_mRNA and exRNAQC017_small RNA)).

**Supplementary Table 6. Small RNA sequencing data statistics of RNA purification kit experiment (exRNAQC011). UniqueID:** RNA identifier; SampleID: combination of kit abbreviation and technical replicate number; raw_reads: number of sequenced (single-end) reads; qcfiltered_reads: number of reads after quality filtering; post_subsampling: number of reads after subsampling; aligned_reads: number of subsampled reads aligned to reference genome; spike_reads: number of reads aligned to spikes; prct_aligned: % of subsampled reads aligned to reference genome; prct_aligned_plus_spikes: % of subsampled reads aligned to reference genome or to spikes.

**Supplementary Table 7. mRNA capture sequencing data statistics of blood collection tube experiment (exRNAQC005).** UniqueID: RNA identifier; SampleID: combination of tube abbreviation, donor number (biological replicate), and time interval; raw_reads: number of sequenced reads pairs; qcfiltered_reads: number of read pairs after quality filtering; post_subsampling: number of read pairs after subsampling; post_deduplication: number of read pairs after Clumpify duplicate removal; duplicate_prct: % of duplicates in subsampled reads; kallisto_prct_alignment: % of duplicate removed reads that were pseudoaligned; strandedness_prct: % of reads on correct strand (stranded protocol).

**Supplementary Table 8. Small RNA sequencing data statistics of blood collection tube experiment (exRNAQC013).** UniqueID: RNA identifier; SampleID: combination of tube abbreviation, donor number (biological replicate), and time interval; raw_reads: number of sequenced (single-end) reads; qcfiltered_reads: number of reads after quality filtering; post_subsampling: number of reads after subsampling; aligned_reads: number of subsampled reads aligned to reference genome; spike_reads: number of reads aligned to spikes; prct_aligned: % of subsampled reads aligned to reference genome; prct_aligned_plus_spikes: % of subsampled reads aligned to reference genome or to spikes.

**Supplementary Table 9. mRNA capture sequencing data statistics of phase 2 (exRNAQC017).** UniqueID: RNA identifier; SampleID: combination of RNA isolation kit abbreviation, tube abbreviation, time interval and donor number (biological replicate); raw_reads: number of sequenced reads pairs; qcfiltered_reads: number of read pairs after quality filtering; post_subsampling: number of read pairs after subsampling; post_deduplication: number of read pairs after Clumpify duplicate removal; duplicate_prct: % of duplicates in subsampled reads; kallisto_prct_alignment: % of duplicate removed reads that were pseudoaligned; strandedness_prct: % of reads on correct strand (stranded protocol).

**Supplementary Table 10. Small RNA sequencing data statistics of phase 2 (exRNAQC017).** UniqueID: RNA identifier; SampleID: combination of RNA isolation kit abbreviation, tube abbreviation, time interval and donor number (biological replicate); raw_reads: number of sequenced (single-end) reads; qcfiltered_reads: number of reads after quality filtering; post_subsampling: number of reads after subsampling; aligned_reads: number of subsampled reads aligned to reference genome; spike_reads: number of reads aligned to spikes; prct_aligned: % of subsampled reads aligned to reference genome; prct_aligned_plus_spikes: % of subsampled reads aligned to reference genome or to spikes.

**Supplementary Table 11. Capture probes for Sequin and External RNA Control Consortium (ERCC) spike-in controls.** Oligos to capture the Sequin and ERCC spike-in controls are listed. For each oligo, the probe_ID, sequence, GC content (%), melting temperature (Tm in °C), ΔG and binding position in the Sequin or ERCC spike-in sequence are given.

### Supplementary Figure legends

**Supplementary Fig. 1: The exRNAQC study represents the most comprehensive analysis of pre-analytics in the context of exRNA profiling.** The exRNAQC study outperforms previous studies analyzing pre-analytics impacting exRNA analyses in terms of the number of evaluated blood collection tubes and RNA purification methods (Supplementary Table 1). Studies are annotated with NA if the number of blood collection tubes or RNA purification methods cannot be clearly determined from the corresponding publication or if no RNA purification was performed.

**Supplementary Fig. 2: Performance of RNA purification kits on duplication rate, coverage and strandedness at mRNA level.** For each of the unique RNA purification-plasma input volume combinations, 3 technical replicates are analyzed. (**a**) Percentage of read duplicates found by Clumpify after subsampling (n = 39). (**b**) Percentage of bases in the total transcriptome that are covered at least once (n = 39). (**c**) Percentage of reads on correct strand according to strand-specific protocol (n = 45). The number that follows the abbreviation of the purification kit is the plasma input volume (in ml).

**Supplementary Fig. 3: Correlation between Femto Pulse-based (in ng/µl) and sequencing-based eluate RNA concentrations.** (**a**) Endogenous mRNA vs ERCC ratio of RNA purification kits. (**b**) Endogenous small RNA vs LP ratio of RNA purification kits. Only samples with a Femto Pulse concentration above the limit of quantification (15 pg/µl) were kept (n = 23 for mRNA capture sequencing, n = 45 for small RNA sequencing). Axes showed in logarithmic scale. Spearman correlation coefficients and p-values are indicated (calculated using the spearmanr function (Scipy library) in Python). CCF: QIAamp ccfDNA/RNA Kit; CIRC: Plasma/Serum Circulating and Exosomal RNA Purification Kit/Slurry Format; ERCC: Extracellular RNA Communication Consortium; LP: Library Prep Control; MAX: Maxwell RSC miRNA Plasma and Exosome Kit in combination with the Maxwell RSC Instrument; MIR: miRNeasy Serum/Plasma Kit; MIRA: miRNeasy Serum/Plasma Advanced Kit; MIRV: mirVana PARIS Kit with purification protocol for total RNA; MIRVE: mirVana PARIS Kit with purification protocol for RNA enriched for small RNAs; NUC: NucleoSpin miRNA Plasma Kit.

**Supplementary Fig. 4: Extracellular RNA is highly fragmented.** For each RNA purification method, FemtoPulse results are shown for one of the triplicate RNA purifications using the maximum plasma input volume. Measurements are on DNase-treated samples (using 2 µl sample), except for MIRVE0.625 (RNA004084; 2 µl RNA eluate). Number that follows the abbreviation of the purification kit is the plasma input volume (in ml). CCF: QIAamp ccfDNA/RNA Kit; CIRC: Plasma/Serum Circulating and Exosomal RNA Purification Kit/Slurry Format; MAX: the Maxwell RSC miRNA Plasma and Exosome Kit in combination with the Maxwell RSC Instrument; MIR: the miRNeasy Serum/Plasma Kit; MIRA: the miRNeasy Serum/Plasma Advanced Kit; MIRV: the mirVana PARIS Kit with purification protocol for total RNA; MIRVE: mirVana PARIS Kit with purification protocol for RNA enriched for small RNAs; NUC: the NucleoSpin miRNA Plasma Kit.

**Supplementary Fig. 5: Performance of RNA purification methods on count threshold, data retention, RNA yield, and extraction efficiency at mRNA and small RNA level.** For each of the unique RNA purification-plasma input volume combinations, 3 technical replicates are analyzed (n = 39 for mRNA capture sequencing, n = 45 for small RNA sequencing). (**a&b**) Count threshold required to eliminate at least 95% of single positive genes or miRNAs, respectively, between technical replicates. (**c&d**) Data retention: % of total counts that are kept after applying count threshold. (**e&f**) RNA yield, obtained by correcting the RNA concentration for eluate volume, values are log rescaled to the lowest mean of all kits and transformed back to linear space, mean and 95% confidence interval are shown. (**g&h**) Extraction efficiency, obtained by correcting the RNA yield for input volume, values are log rescaled to the lowest mean of all kits and transformed back to linear space, mean and 95% confidence interval are shown. Number that follows the abbreviation of the purification kit is the plasma input volume (in ml).

**Supplementary Fig. 6: Illustrative example of performance metric evolution over time** for one donor, two blood collection tubes and three time intervals (**a**) and corresponding boxplot of the fold changes per blood collection tube (**b**). T0: plasma prepared immediately after blood draw, T24, T72: plasma prepared 24 hours and 72 hours after blood draw, respectively. The white triangle on the boxplot corresponds to the mean. Reproduced from Van Paemel *et al.*^53^

**Supplementary Fig. 7: Performance metrics of blood collection tubes over time at mRNA level.** (**a**) Evolution of hemolysis in plasma, measured by absorbance at 414 nm with Nanodrop. (**b**) Evolution of RNA concentration calculated based on the number of endogenous counts vs Sequin spike-in RNA. (**c**) Evolution of sensitivity, i.e., the number of protein coding genes. (**d**) Evolution of the fraction of counts mapping to mRNAs versus all counts (biotype performance metric). (**e**) Evolution of the pairwise area left of the curve (reproducibility performance metric). T0: plasma prepared immediately after blood draw. T04, T16, T24, T72: plasma prepared 4, 16, 24 and 72 hours after blood draw, respectively. Note that different donors were sampled and that tubes were processed at different time intervals for preservation and non-preservation tubes. ACD-A: BD Vacutainer Glass ACD Solution A tube; Biomatrica: LBgard Blood Tube; Citrate: Vacuette Tube 9 ml 9NC Coagulation sodium citrate 3.2%; DNA Streck: Cell-Free DNA BCT; EDTA: BD Vacutainer Plastic K2EDTA tube; EDTA separator: Vacuette Tube 8 ml K2E K2EDTA Separator; PAXgene: PAXgene Blood ccfDNA Tube; RNA Streck: Cell-Free RNA BCT; Roche: Cell-Free DNA Collection Tube; Serum: BD Vacutainer SST II Advance Tube.

**Supplementary Fig. 8: Performance metrics of blood collection tubes over time at small RNA level.** (**a**) Evolution of hemolysis in plasma, measured by absorbance at 414 nm with Nanodrop. (**b**) Evolution of RNA concentration calculated based on number of endogenous counts vs RC spike-in RNA. (**c**) Evolution of sensitivity, i.e., the number miRNAs. (**d**) Evolution of the fraction of counts mapping to miRNAs versus all counts (biotype performance metric). (**e**) Evolution of the pairwise area left of the curve (reproducibility performance metrics). T0: plasma prepared immediately after blood draw. T04, T16, T24, T72: plasma prepared 4, 16, 24 and 72 hours after blood draw, respectively. Note that different donors were sampled and that tubes were processed at different time intervals for preservation and non-preservation tubes. ACD-A: BD Vacutainer Glass ACD Solution A tube; Biomatrica: LBgard Blood Tube; Citrate: Vacuette Tube 9 ml 9NC Coagulation sodium citrate 3.2%; DNA Streck: Cell-Free DNA BCT; EDTA: BD Vacutainer Plastic K2EDTA tube; EDTA separator: Vacuette Tube 8 ml K2E K2EDTA Separator; PAXgene: PAXgene Blood ccfDNA Tube; RNA Streck: Cell-Free RNA BCT; Roche: Cell-Free DNA Collection Tube; Serum: BD Vacutainer SST II Advance Tube.

**Supplementary Fig. 9: Example of hemolysis in preservation tubes.** (a) Visual inspection of non-preservation plasma tubes of donor 7 (Supplementary Fig. 7a and Supplementary Fig. 8a) and (b) of preservation plasma tubes of donor 5 (Supplementary Fig. 7a and Supplementary Fig. 8a) at time interval T0. For donor 5, plasma from the PAXgene, RNA Streck and Roche tube showed to be hemolytic, which is in line with the NanoDrop measurements (Supplementary Fig. 7a and Supplementary Fig. 8a). ACD-A: BD Vacutainer Glass ACD Solution A tube; Biomatrica: LBgard Blood Tube; **C**itrate: Vacuette Tube 9 ml 9NC Coagulation sodium citrate 3.2%; DNA Streck: Cell-Free DNA BCT; EDTA: BD Vacutainer Plastic K2EDTA tube; EDTA separator: Vacuette Tube 8 ml K2E K2EDTA Separator; PAXgene: PAXgene Blood ccfDNA Tube; RNA Streck: Cell-Free RNA BCT; Roche: Cell-Free DNA Collection Tube.

**Supplementary Fig. 10: Fold changes over time at mRNA level for each blood collection tube performance metric.** (**a**) Boxplot of the fold change within each donor across time intervals, per tube, for hemolysis as measured by absorbance at 414 nm with Nanodrop. (**b**) Boxplot of fold change of plasma RNA concentration, based on the ratio of endogenous vs Sequin spike-in RNA reads. (**c**) Boxplot of the fold change of the sensitivity, i.e., the number of genes after filtering out genes with counts fewer than 6 reads. (**d**) Reproducibility, i.e., area left of the curve, transformed from log2 to linear scale. (**e**) Boxplot of the fold change of the fraction of the counts mapping to protein coding genes versus all counts (biotype performance metric). In the boxplots, the lower and upper hinge of the boxes represents the 25th and 75th percentile, respectively. The whiskers extend to the lowest and highest value that is within 1.5 times the interquartile range. Data beyond the end of the whiskers are outliers. The white triangle on the boxplot corresponds to the mean of the fold change. Individual data points are shown as colored dots (for non-preservation tubes) or triangles (for preservation tubes). The first time interval corresponds to the comparison of T04 versus T0 (non-preservation tubes) or T24 versus T0 (preservation tubes). The second time interval corresponds to the comparison of T16 versus T0 (non-preservation tubes) or T72 vs T0 (preservation tubes). T0: plasma prepared immediately after blood draw. T04, T16, T24, T72: plasma prepared 4, 16, 24 and 72 hours after blood draw, respectively. Note that different donors were sampled and that tubes were processed at different time intervals for preservation and non-preservation tubes. ACD-A: BD Vacutainer Glass ACD Solution A tube; Biomatrica: LBgard Blood Tube; Citrate: Vacuette Tube 9 ml 9NC Coagulation sodium citrate 3.2%; DNA Streck: Cell-Free DNA BCT; EDTA: BD Vacutainer Plastic K2EDTA tube; EDTA separator: Vacuette Tube 8 ml K2E K2EDTA Separator; PAXgene: PAXgene Blood ccfDNA Tube; RNA Streck: Cell-Free RNA BCT; Roche: Cell-Free DNA Collection Tube; Serum: BD Vacutainer SST II Advance Tube.

**Supplementary Fig. 11: Fold changes over time at small RNA level for each blood collection tube performance metric.** (**a**) Boxplot of the fold change within each donor across time intervals, per tube, for hemolysis, as measured by absorbance at 414 nm with Nanodrop. (**b**) Boxplot of fold change of plasma RNA concentration, based on the ratio of endogenous vs RC spike-in RNA reads. (**c**) Boxplot of the fold change of the sensitivity, i.e., the number of miRNAs after filtering out miRNAs with counts fewer than 3 reads. (**d**) Reproducibility, i.e., area left of the curve, transformed from log2 to linear scale. (**e**) Boxplot of the fold change of the fraction of the counts mapping to miRNAs versus all counts (biotype performance metric). In the boxplots, the lower and upper hinge of the boxes represents the 25th and 75th percentile, respectively. The whiskers extend to the lowest and highest value that is within 1.5 times the interquartile range. Data beyond the end of the whiskers are outliers. The white triangle on the boxplot corresponds to the mean of the fold change. Individual data points are shown as colored dots (for non-preservation tubes) or triangles (for preservation tubes). The first time interval corresponds to the comparison of T04 versus T0 (non-preservation tubes) or T24 versus T0 (preservation tubes). The second time interval corresponds to the comparison of T16 versus T0 (non-preservation tubes) or T72 vs T0 (preservation tubes). T0: plasma prepared immediately after blood draw. T04, T16, T24, T72: plasma prepared 4, 16, 24 and 72 hours after blood draw, respectively. Note that different donors were sampled and that tubes were processed at different time intervals for preservation and non-preservation tubes. ACD-A: BD Vacutainer Glass ACD Solution A tube; Biomatrica: LBgard Blood Tube; Citrate: Vacuette Tube 9 ml 9NC Coagulation sodium citrate 3.2%; DNA Streck: Cell-Free DNA BCT; EDTA: BD Vacutainer Plastic K2EDTA tube; EDTA separator: Vacuette Tube 8 ml K2E K2EDTA Separator; PAXgene: PAXgene Blood ccfDNA Tube; RNA Streck: Cell-Free RNA BCT; Roche: Cell-Free DNA Collection Tube; Serum: BD Vacutainer SST II Advance Tube.

**Supplementary Fig. 12: Total number of circular and linear reads relative to T0.** Each white dot with colored outline represents the number of reads relative to the same donor’s T0 tube. The fully colored dots represent the average relative number of reads and 95% confidence intervals are shown. Each plot represents a different tube type. The relative number of linear and circular reads are calculated by multiplying the fraction of the endogenous RNA reads and the Sequin spikes by the fraction of linear or circRNA reads, respectively. T0: plasma prepared immediately after blood draw. T04, T16, T24, T72: plasma prepared 4, 16, 24 and 72 hours after blood draw, respectively. Note that different donors were sampled and that tubes were processed at different time intervals for preservation and non-preservation tubes. ACD-A: BD Vacutainer Glass ACD Solution A tube; Biomatrica: LBgard Blood Tube; Citrate: Vacuette Tube 9 ml 9NC Coagulation sodium citrate 3.2%; DNA Streck: Cell-Free DNA BCT; EDTA: BD Vacutainer Plastic K2EDTA tube; EDTA separator: Vacuette Tube 8 ml K2E K2EDTA Separator; PAXgene: PAXgene Blood ccfDNA Tube; RNA Streck: Cell-Free RNA BCT; Roche: Cell-Free DNA Collection Tube; Serum: BD Vacutainer SST II Advance Tube.

**Supplementary Fig. 13: RNA abundance levels differ across blood collection tubes.** Using normalized and scaled count data, gene abundance levels (i.e., mean abundance of the tube replicates) are shown for each tube type at time interval T0. The normalised counts are scaled between −1 (low abundance) and 1 (high abundance) per gene to make abundance differences across genes comparable. Only genes with ≥ 10 counts in all three replicates of one tube type were included. T0: plasma prepared immediately after blood draw. Note that different donors were sampled for preservation and non-preservation tubes. ACD-A: BD Vacutainer Glass ACD Solution A tube; Biomatrica: LBgard Blood Tube; Citrate: Vacuette Tube 9 ml 9NC Coagulation sodium citrate 3.2%; DNA Streck: Cell-Free DNA BCT; EDTA: BD Vacutainer Plastic K2EDTA tube; EDTA separator: Vacuette Tube 8 ml K2E K2EDTA Separator; PAXgene: PAXgene Blood ccfDNA Tube; RNA Streck: Cell-Free RNA BCT; Roche: Cell-Free DNA Collection Tube; Serum: BD Vacutainer SST II Advance Tube.

**Supplementary Fig. 14: RNA abundance levels differ across time intervals.** For each blood collection tube in exRNAQC phase 1 (a) and 2 (b), distributions of log2 fold changes between time interval 1 and 0 (T0), and time interval 2 and 0 (T0) are shown. For the non-preservation tubes, time interval 1 corresponds to T04 and time interval 2 to T16. For the preservation tubes, time interval 1 corresponds to T24 and time interval 2 to T72. T0: plasma prepared immediately after blood draw. T04, T16, T24, T72: plasma prepared 4, 16, 24 and 72 hours after blood draw, respectively. Note that different donors were sampled and that tubes were processed at different time intervals for preservation and non-preservation tubes. ACD-A: BD Vacutainer Glass ACD Solution A tube; Biomatrica: LBgard Blood Tube; Citrate: Vacuette Tube 9 ml 9NC Coagulation sodium citrate 3.2%; DNA Streck: Cell-Free DNA BCT; EDTA: BD Vacutainer Plastic K2EDTA tube; EDTA separator: Vacuette Tube 8 ml K2E K2EDTA Separator; PAXgene: PAXgene Blood ccfDNA Tube; RNA Streck: Cell-Free RNA BCT; Roche: Cell-Free DNA Collection Tube; Serum: BD Vacutainer SST II Advance Tube.

**Supplementary Fig. 15: Kit selection for exRNAQC phase 2 for mRNA capture (a) and small RNA (b) sequencing.** Median robust z-score (see Methods) per kit-input volume combination (13 in a, 15 in b) shown for sensitivity and reproducibility metrics; Number that follows the abbreviation of the purification kit is the plasma input volume (in ml).

**Supplementary Fig. 16: Data access through the R2 Genomics Analysis and Visualization Platform enables browsable result access for any researcher to mine and analyze the exRNAQC data**. Shown are abundance levels of a gene (C5AR1), identified by the gene set enrichment analyses on the data of exRNAQC phase 2 as differentially abundant between time interval T0 and T16 in EDTA. Following the settings in the upper panel to ‘View a gene in groups’, this can be nicely visualized in R2 (lower panel). Red panel is Citrate, green is EDTA, purple is Serum. In the annotation track ‘hours_to_start_processing’, red represents T0, green T04, and blue T16. Mouse hover actions enable to visualize these annotations on the online platform.

## Supplementary Information

