## Supplementary Fig 1 for "Performance evaluation of RNA purification kits and blood collection tubes in the Extracellular RNA Quality Control (exRNAQC) study"

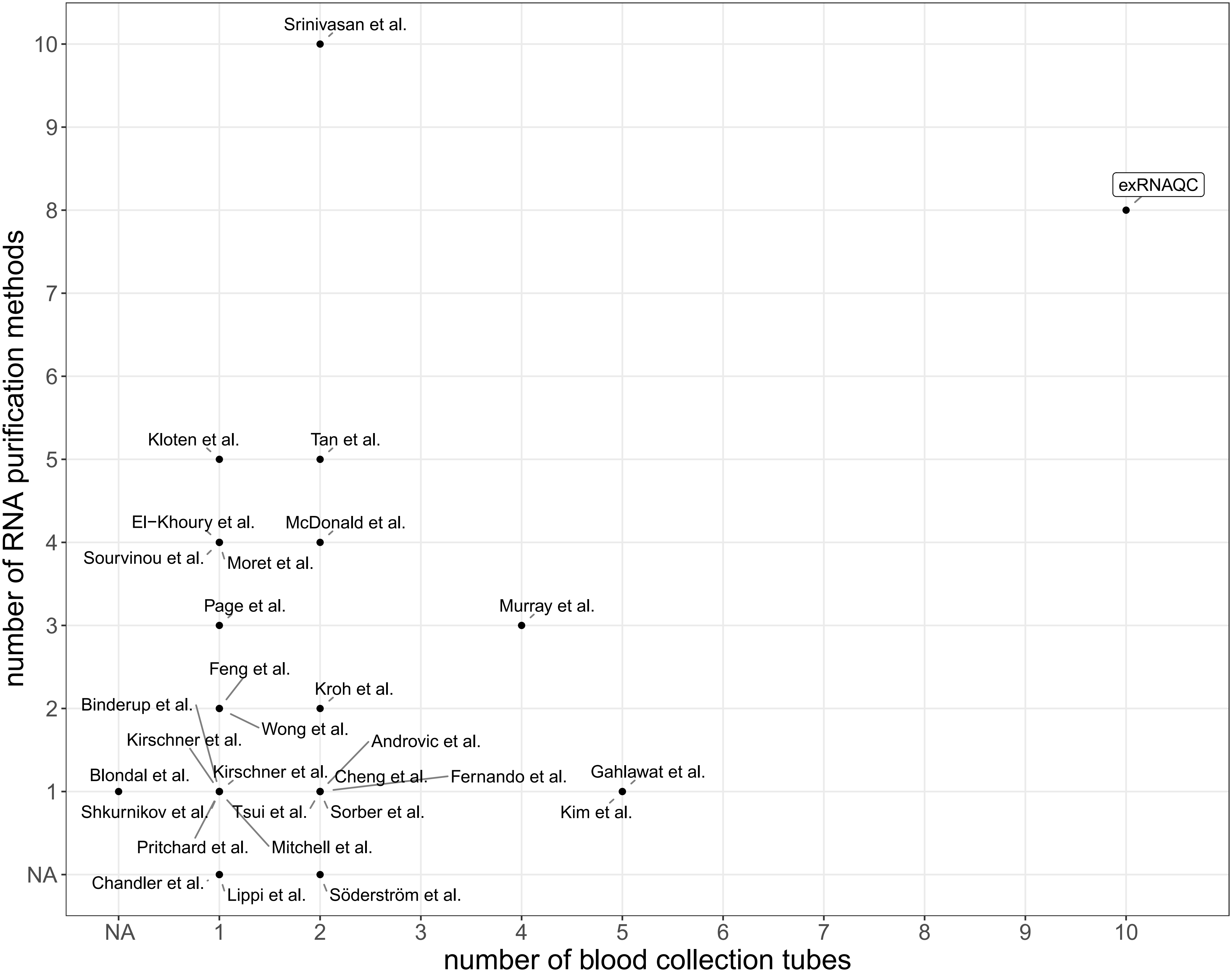


**Supplementary Fig. 1: The exRNAQC study represents the most comprehensive analysis of pre-analytics in the context of exRNA profiling.** The exRNAQC study outperforms previous studies analyzing pre-analytics impacting exRNA analyses in terms of the number of evaluated blood collection tubes and RNA purification methods (Supplementary Table 1). Studies are annotated with NA if the number of blood collection tubes or RNA purification methods cannot be clearly determined from the corresponding publication or if no RNA purification was performed.
