## Supplementary Fig 2 for "Performance evaluation of RNA purification kits and blood collection tubes in the Extracellular RNA Quality Control (exRNAQC) study"

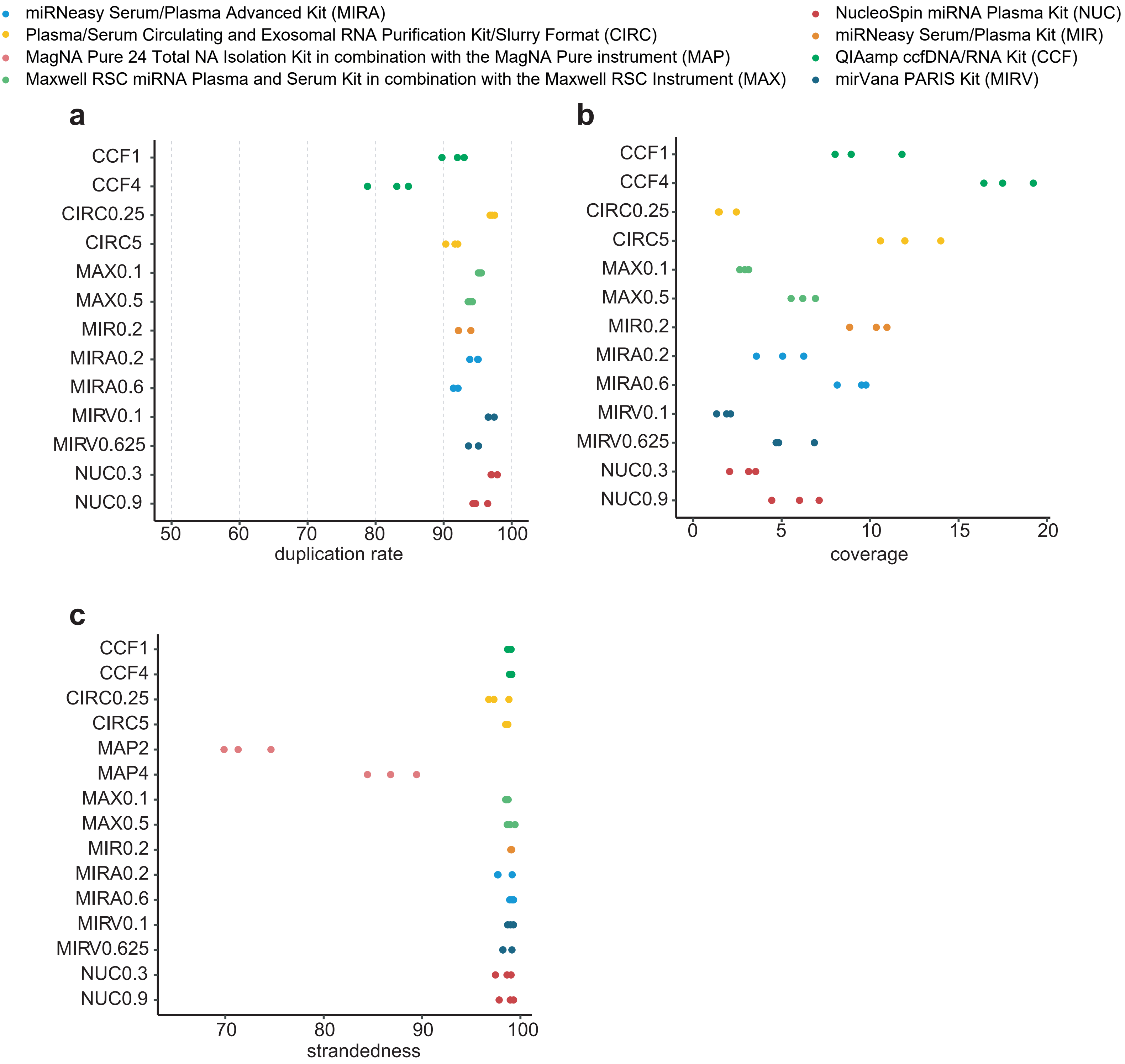


**Supplementary Fig. 2: Performance of RNA purification kits on duplication rate, coverage and strandedness at mRNA level.** For each of the unique RNA purification-plasma input volume combinations, 3 technical replicates are analyzed. **(a)** Percentage of read duplicates found by Clumpify after subsampling (n = 39). **(b)** Percentage of bases in the total transcriptome that are covered at least once (n = 39). **(c)** Percentage of reads on correct strand according to strand-specific protocol (n = 45). The number that follows the abbreviation of the purification kit is the plasma input volume (in ml).
