## Supplementary Fig 3 for "Performance evaluation of RNA purification kits and blood collection tubes in the Extracellular RNA Quality Control (exRNAQC) study"

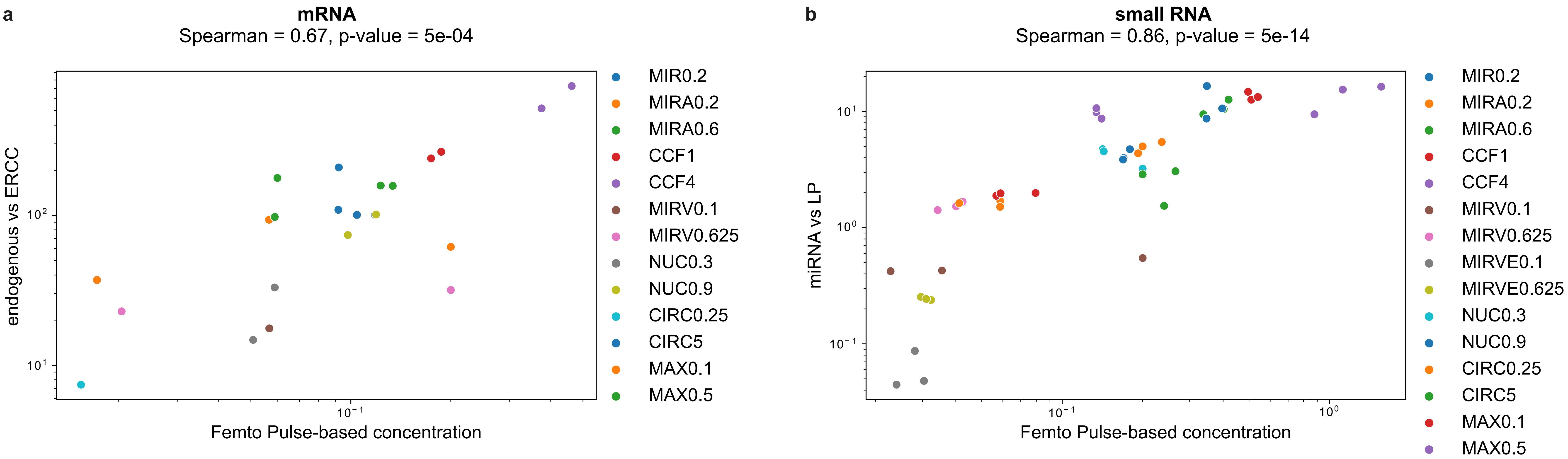


**Supplementary Fig. 3: Correlation between Femto Pulse-based (in ng/µl) and sequencing-based eluate RNA concentrations. (a)** Endogenous mRNA vs ERCC ratio of RNA purification kits. **(b)** Endogenous small RNA vs LP ratio of RNA purification kits. Only samples with a Femto Pulse concentration above the limit of quantification (15 pg/µl) were kept (n = 23 for mRNA capture sequencing, n = 45 for small RNA sequencing). Axes showed in logarithmic scale. Spearman correlation coefficients and p-values are indicated (calculated using the spearmanr function (Scipy library) in Python). CCF: QIAamp ccfDNA/RNA Kit; CIRC: Plasma/Serum Circulating and Exosomal RNA Purification Kit/Slurry Format; ERCC: Extracellular RNA Communication Consortium; LP: Library Prep Control; MAX: Maxwell RSC miRNA Plasma and Exosome Kit in combination with the Maxwell RSC Instrument; MIR: miRNeasy Serum/Plasma Kit; MIRA: miRNeasy Serum/Plasma Advanced Kit; MIRV: mirVana PARIS Kit with purification protocol for total RNA; MIRVE: mirVana PARIS Kit with purification protocol for RNA enriched for small RNAs; NUC: NucleoSpin miRNA Plasma Kit.
