## Supplementary Fig 4 for "Performance evaluation of RNA purification kits and blood collection tubes in the Extracellular RNA Quality Control (exRNAQC) study"

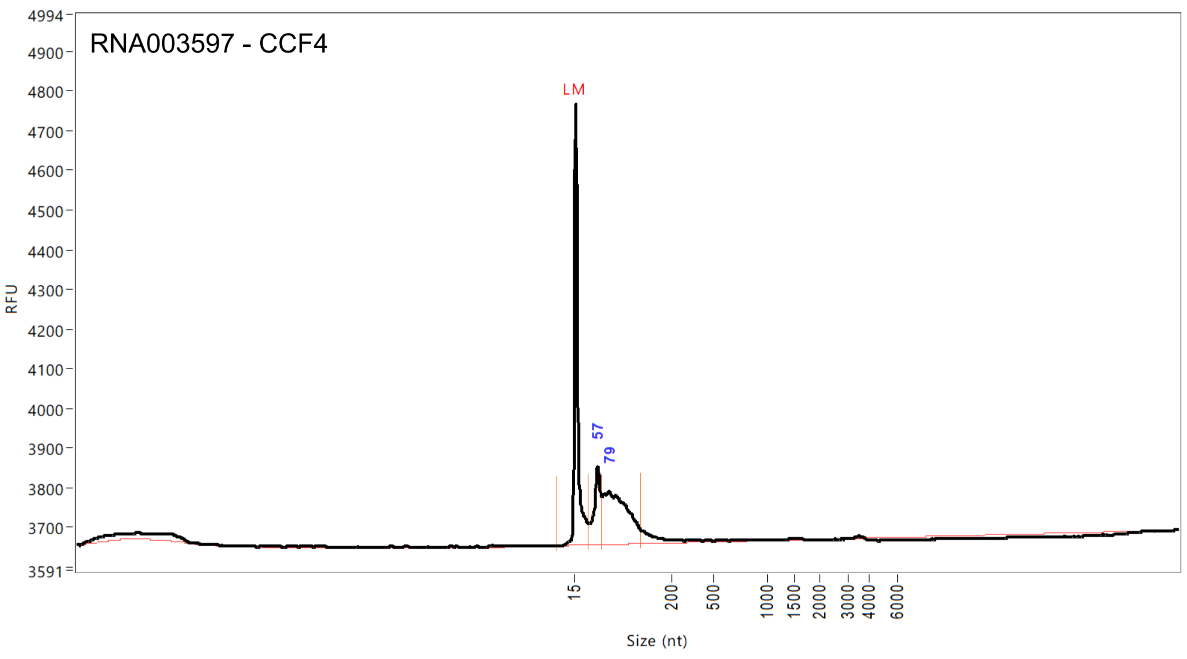


| Peak | Size (nt) | Conc. (pg/uL) | From (nt) | To (nt) | RFU |
|---|---|---|---|---|---|
| 1 | 15 (LM) | 6.2267 | 0 | 40 | 1113 |
| 2 | 57 | 20.7608 | 40 | 66 | 195 |
| 3 | 79 | 50.1091 | 66 | 142 | 132 |

| | | |
|---|---|---|
| TIC: | 70.8698 | pg/uL |
| TIM: | 2787.4565 | pmole/L |
| Total Conc.: | 77.1040 | pg/uL |
| 28S/18S: | 0.0 | |
| RQN | 1.2 | |

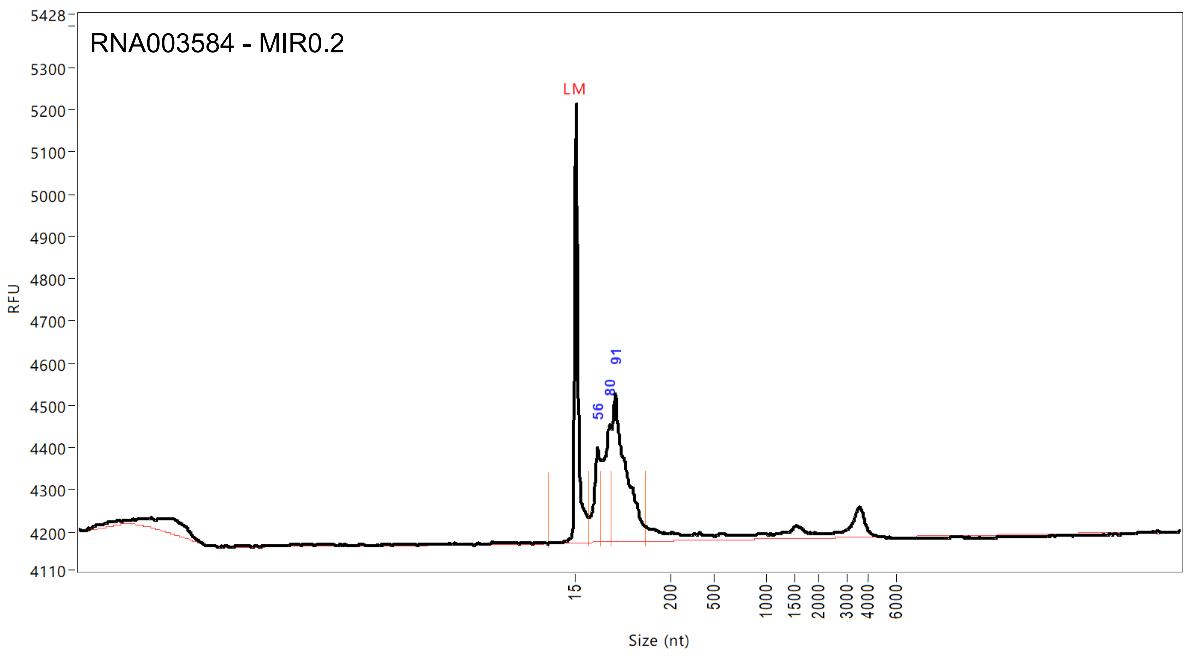


| Peak | Size (nt) | Conc. (pg/uL) | From (nt) | To (nt) | RFU |
|---|---|---|---|---|---|
| 1 | 15 (LM) | 6.2267 | 0 | 40 | 1040 |
| 2 | 56 | 23.9809 | 40 | 63 | 222 |
| 3 | 80 | 34.2518 | 63 | 84 | 277 |
| 4 | 91 | 73.7274 | 84 | 150 | 351 |

| | | |
|---|---|---|
| TIC: | 131.9601 | pg/uL |
| TIM: | 4967.6406 | pmole/L |
| Total Conc.: | 164.9395 | pg/uL |
| 28S/18S: | 0.0 | |
| RQN | 3.2 | |

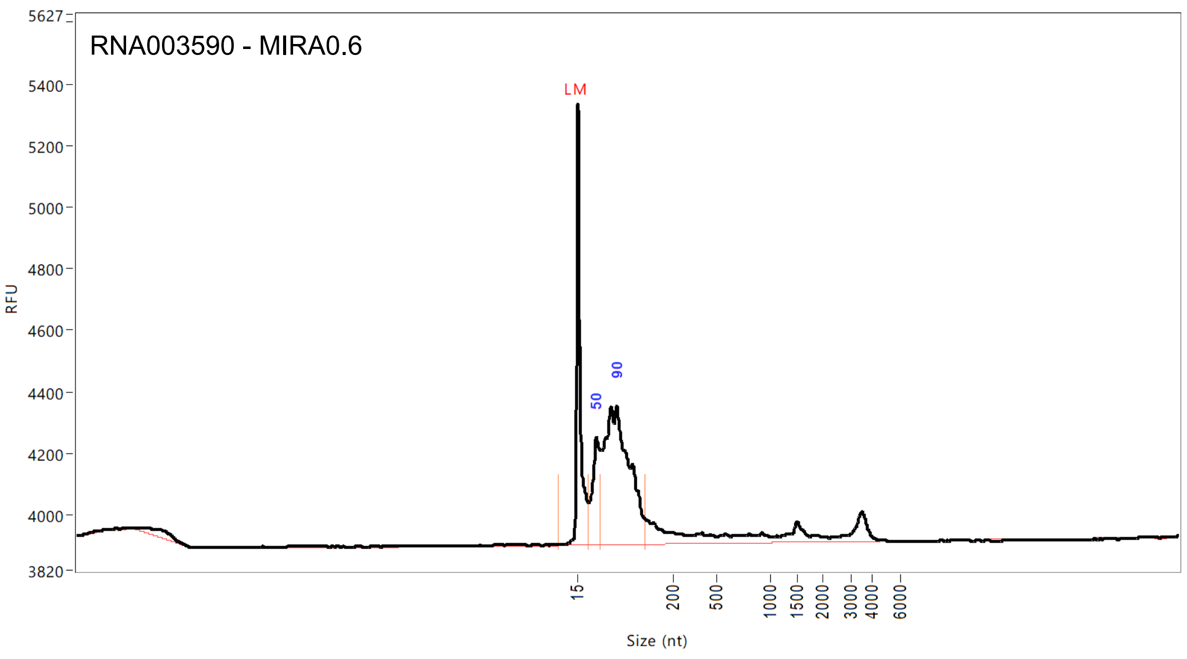


| Peak | Size (nt) | Conc. (pg/uL) | From (nt) | To (nt) | RFU |
|---|---|---|---|---|---|
| 1 | 15 (LM) | 6.2267 | 0 | 35 | 1434 |
| 2 | 50 | 32.7797 | 35 | 60 | 348 |
| 3 | 90 | 123.2868 | 60 | 145 | 448 |

| | | |
|---|---|---|
| TIC: | 156.0664 | pg/uL |
| TIM: | 6137.1094 | pmole/L |
| Total Conc.: | 205.0533 | pg/uL |
| 28S/18S: | 0.0 | |
| RQN | 2.4 | |

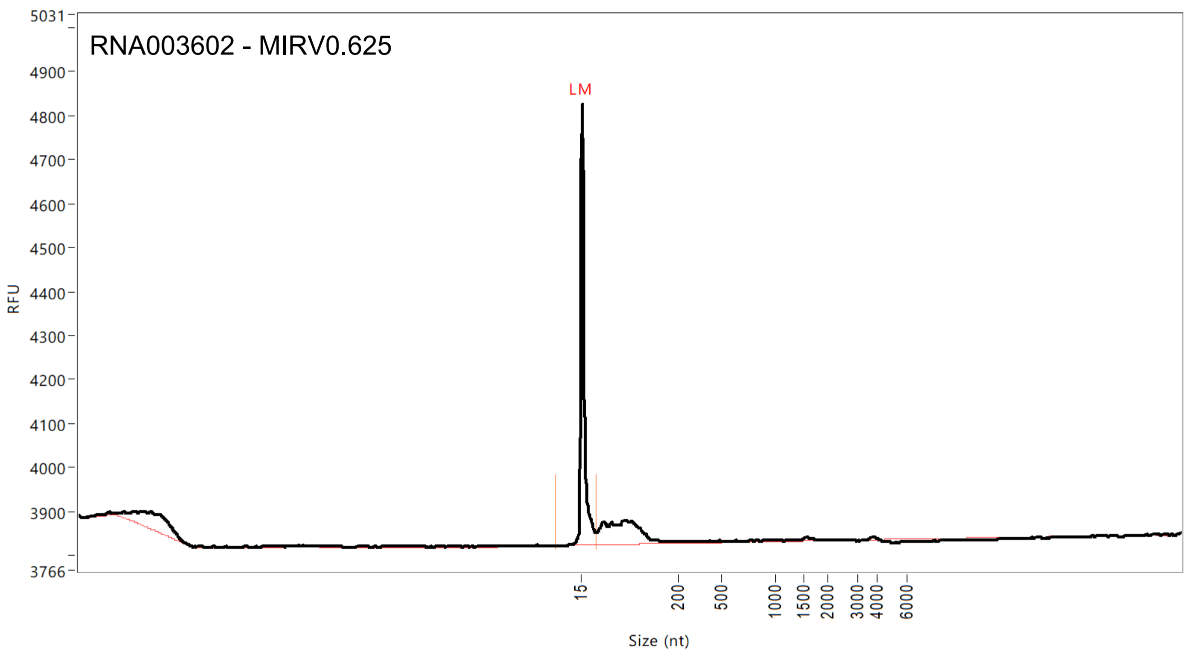


| Peak | Size (nt) | Conc. (pg/uL) | From (nt) | To (nt) | RFU |
|---|---|---|---|---|---|
| 1 | 15 (LM) | 6.2267 | 0 | 42 | 1003 |

| | | |
|---|---|---|
| TIC: | 0.0000 | pg/uL |
| TIM: | 0.0000 | pmole/L |
| Total Conc.: | 32.1047 | pg/uL |
| 28S/18S: | 0.0 | |
| RQN | 10.0 | |

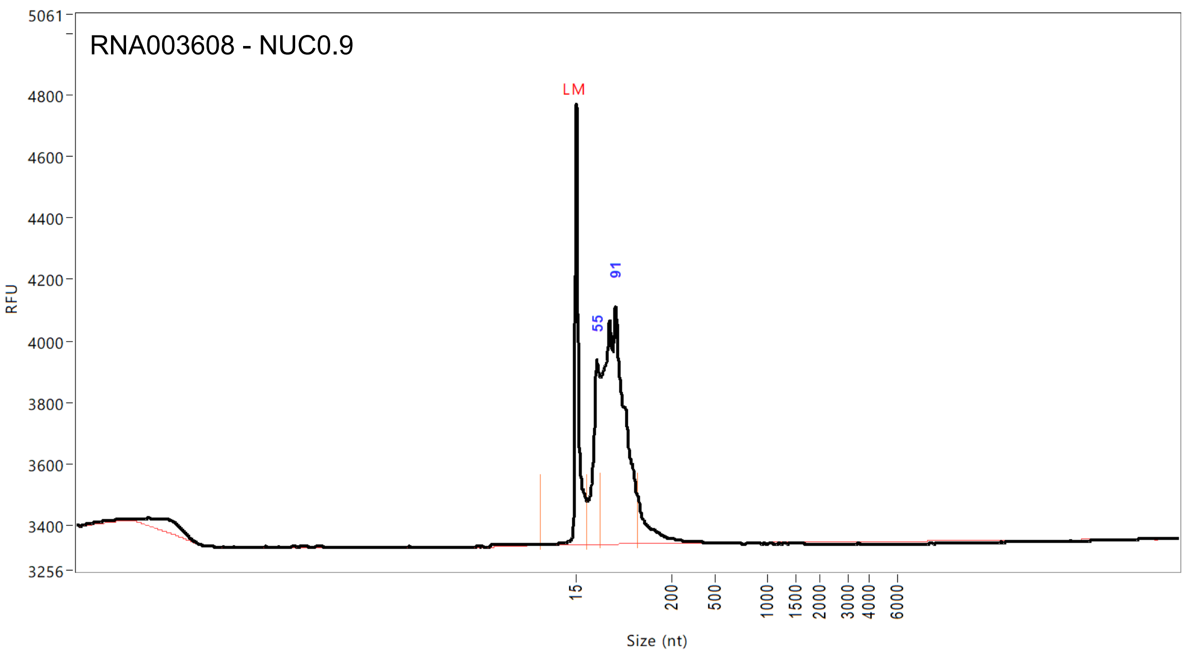


| Peak | Size (nt) | Conc. (pg/uL) | From (nt) | To (nt) | RFU |
|---|---|---|---|---|---|
| 1 | 15 (LM) | 6.2267 | 0 | 37 | 1431 |
| 2 | 55 | 45.9980 | 37 | 62 | 596 |
| 3 | 91 | 173.9776 | 62 | 134 | 769 |

| | | |
|---|---|---|
| TIC: | 219.9756 | pg/uL |
| TIM: | 8666.7471 | pmole/L |
| Total Conc.: | 233.0036 | pg/uL |
| 28S/18S: | 0.0 | |
| RQN | 1.0 | |

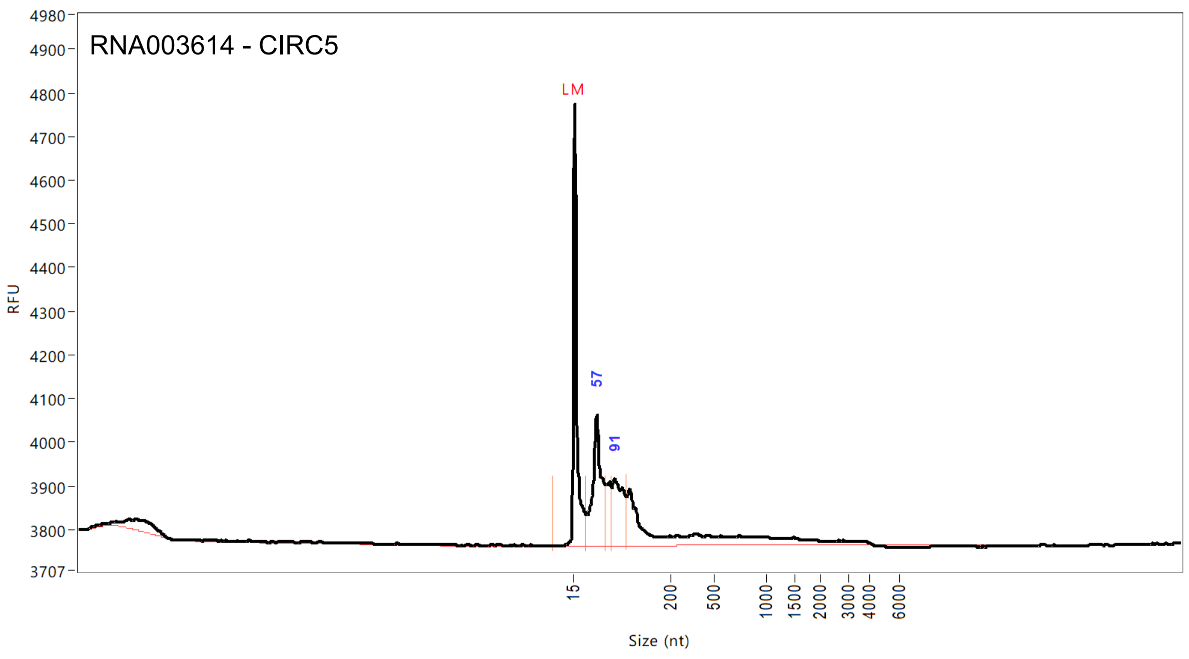


| | Size (nt) | Conc. (pg/uL) | From (nt) | To (nt) | RFU |
|---|---|---|---|---|---|
| Peak | 15 (LM) | 8.3737 | 0 | 38 | 1011 |
| | 57 | 62.2613 | 38 | 75 | 297 |
| | 91 | 37.7524 | 87 | 114 | 151 |
| 1 | | | | | |
| 2 | | | | | |
| 3 | | | | | |

| | | |
|---|---|---|
| TIC: | 100.0137 | pg/uL |
| TIM: | 4493.0649 | pmole/L |
| Total Conc.: | 193.0621 | pg/uL |
| 28S/18S: | 0.0 | |
| RQN | 1.5 | |

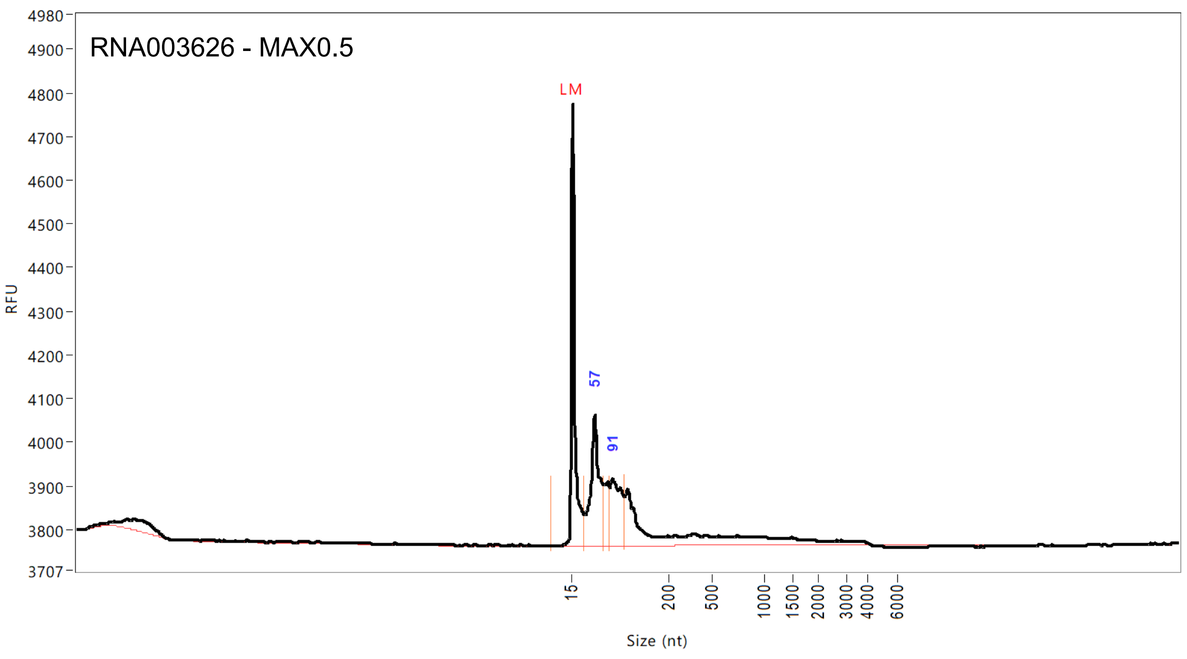


| Peak | Size (nt) | Conc. (pg/uL) | From (nt) | To (nt) | RFU |
|---|---|---|---|---|---|
| 1 | 15 (LM) | 8.3737 | 0 | 38 | 1011 |
| 2 | 57 | 62.2613 | 38 | 75 | 297 |
| 3 | 91 | 37.7524 | 87 | 114 | 151 |

| | | |
|---|---|---|
| TIC: | 100.0137 | pg/uL |
| TIM: | 4493.0649 | pmole/L |
| Total Conc.: | 193.0621 | pg/uL |
| 28S/18S: | 0.0 | |
| RQN | 1.5 | |

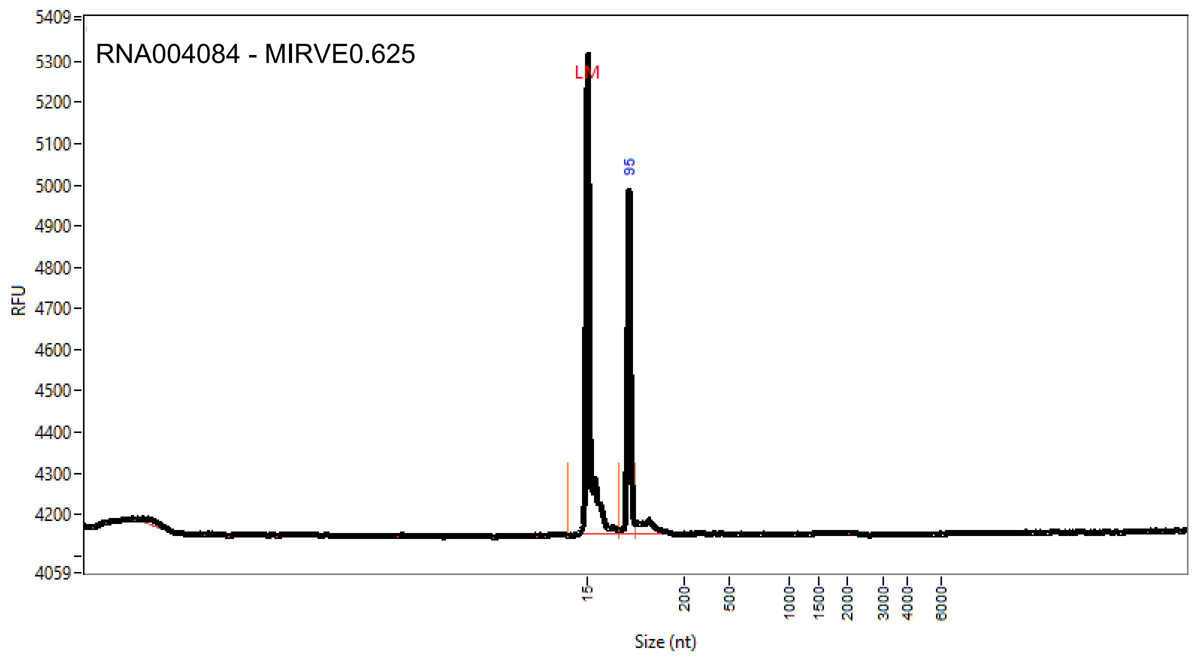


| Peak | Size (nt) | Conc. (pg/uL) | From (nt) | To (nt) | RFU |
|---|---|---|---|---|---|
| 1 | 15 (LM) | 4.7 | 0 | 73 | 1163 |
| 2 | 95 | 25.7 | 73 | 107 | 832 |

| | | |
|---|---|---|
| TIC: | 25.7 | pg/uL |
| TIM: | 847.6 | pmole/L |
| Total Conc.: | 29.6 | pg/uL |
| 28S/18S: | 0.0 | |
| RQN | 1.0 | |

**Supplementary Fig. 4: Extracellular RNA is highly fragmented.** For each RNA purification method, FemtoPulse results are shown for one of the triplicate RNA purifications using the maximum plasma input volume. Measurements are on DNase-treated samples (using 2 µl sample), except for MIRVE0.625 (RNA004084; 2 µl RNA eluate). Number that follows the abbreviation of the purification kit is the plasma input volume (in ml). CCF: QIAamp ccfDNA/RNA Kit; CIRC: Plasma/Serum Circulating and Exosomal RNA Purification Kit/Slurry Format; MAX: the Maxwell RSC miRNA Plasma and Exosome Kit in combination with the Maxwell RSC Instrument; MIR: the miRNeasy Serum/Plasma Kit; MIRA: the miRNeasy Serum/Plasma Advanced Kit; MIRV: the mirVana PARIS Kit with purification protocol for total RNA; MIRVE: mirVana PARIS Kit with purification protocol for RNA enriched for small RNAs; NUC: the NucleoSpin miRNA Plasma Kit.
