## Supplementary Fig 5 for "Performance evaluation of RNA purification kits and blood collection tubes in the Extracellular RNA Quality Control (exRNAQC) study"

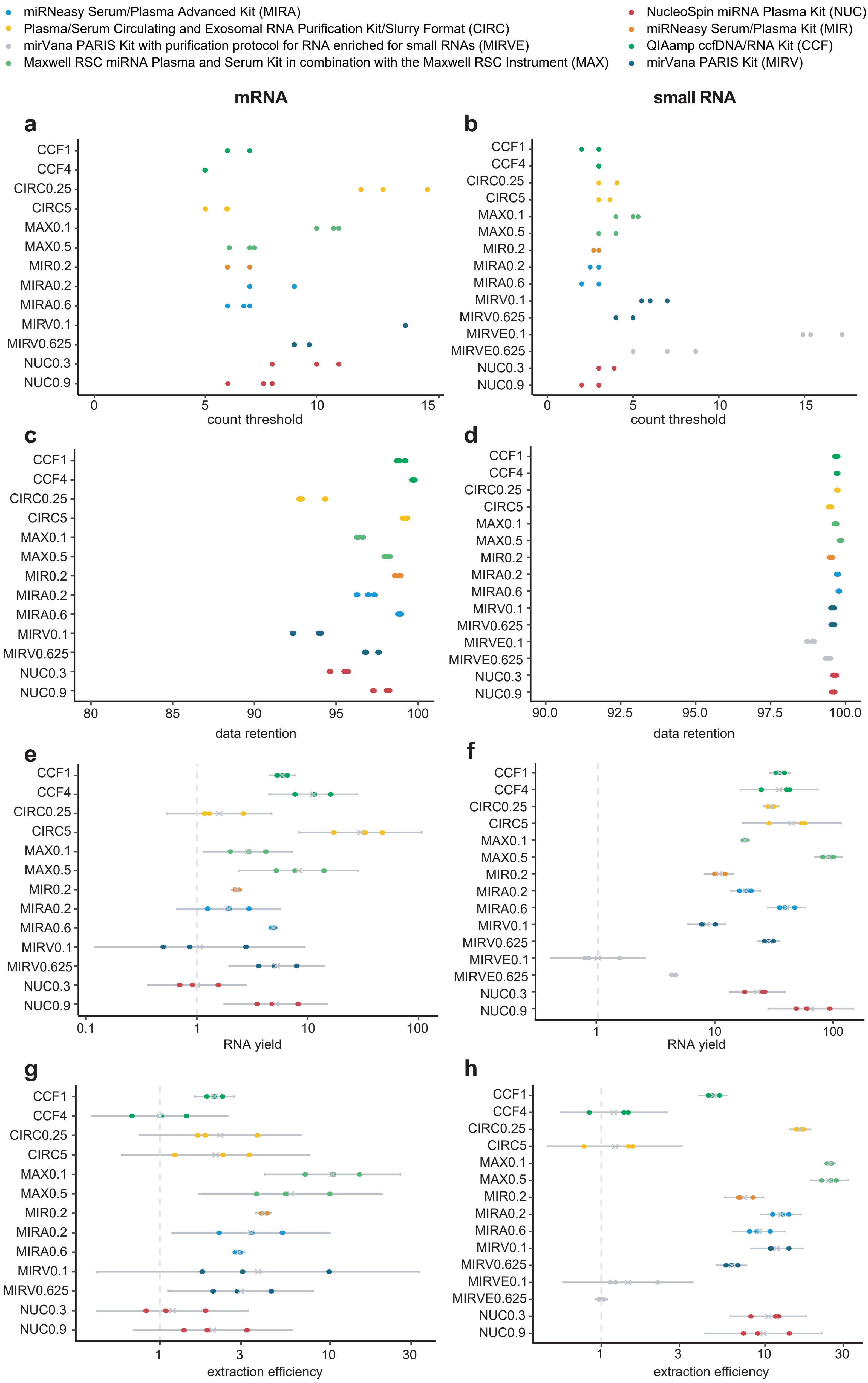


**Supplementary Fig. 5: Performance of RNA purification methods on count threshold, data retention, RNA yield, and extraction efficiency at mRNA and small RNA level.** For each of the unique RNA purification-plasma input volume combinations, 3 technical replicates are analyzed (n = 39 for mRNA capture sequencing, n = 45 for small RNA sequencing). **(a&b)** Count threshold required to eliminate at least 95% of single positive genes or miRNAs, respectively, between technical replicates. **(c&d)** Data retention: % of total counts that are kept after applying count threshold. **(e&f)** RNA yield, obtained by correcting the RNA concentration for eluate volume, values are log rescaled to the lowest mean of all kits and transformed back to linear space, mean and 95% confidence interval are shown. **(g&h)** Extraction efficiency, obtained by correcting the RNA yield for input volume, values are log rescaled to the lowest mean of all kits and transformed back to linear space, mean and 95% confidence interval are shown. Number that follows the abbreviation of the purification kit is the plasma input volume (in ml).
