## Supplementary Fig 6 for "Performance evaluation of RNA purification kits and blood collection tubes in the Extracellular RNA Quality Control (exRNAQC) study"

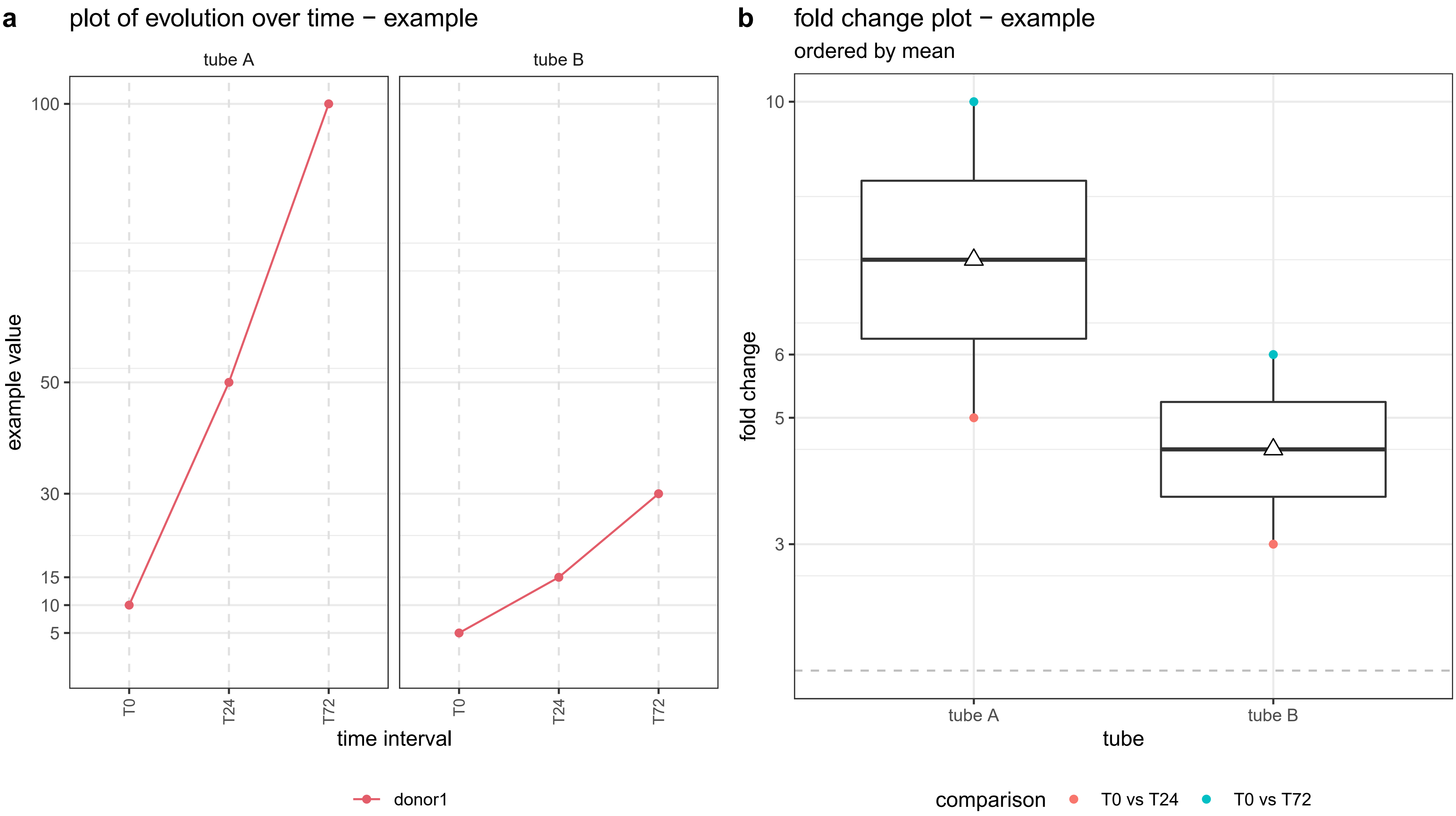


**Supplementary Fig. 6: Illustrative example of performance metric evolution over time** for one donor, two blood collection tubes and three time intervals (a) and corresponding boxplot of the fold changes per blood collection tube (b). T0: plasma prepared immediately after blood draw, T24, T72: plasma prepared 24 hours and 72 hours after blood draw, respectively. The white triangle on the boxplot corresponds to the mean. Reproduced from Van Paemel et al.[53]
