## Supplementary Fig 8 for "Performance evaluation of RNA purification kits and blood collection tubes in the Extracellular RNA Quality Control (exRNAQC) study"

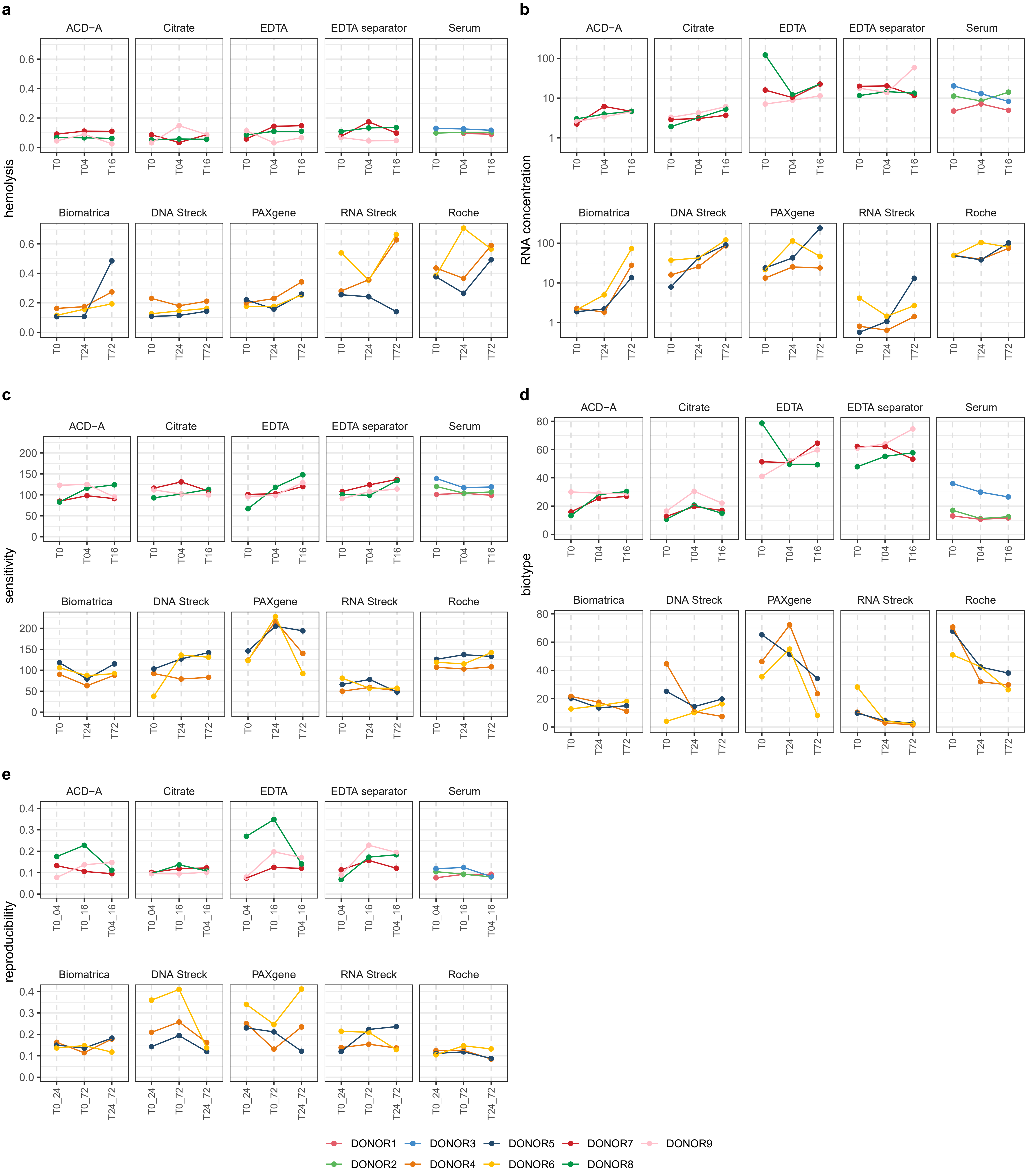


**Supplementary Fig. 8: Performance metrics of blood collection tubes over time at small RNA level. (a)** Evolution of hemolysis in plasma, measured by absorbance at 414 nm with Nanodrop. **(b)** Evolution of RNA concentration calculated based on number of endogenous counts vs RC spike-in RNA. **(c)** Evolution of sensitivity, i.e., the number miRNAs. **(d)** Evolution of the fraction of counts mapping to miRNAs versus all counts (biotype performance metric). **(e)** Evolution of the pairwise area left of the curve (reproducibility performance metrics). T0: plasma prepared immediately after blood draw. T04, T16, T24, T72: plasma prepared 4, 16, 24 and 72 hours after blood draw, respectively. Note that different donors were sampled and that tubes were processed at different time intervals for preservation and non-preservation tubes. ACD-A: BD Vacutainer Glass ACD Solution A tube; Biomatrica: LBgard Blood Tube; Citrate: Vacuette Tube 9 ml 9NC Coagulation sodium citrate 3.2%; DNA Streck: Cell-Free DNA BCT; EDTA: BD Vacutainer Plastic K2EDTA tube; EDTA separator: Vacuette Tube 8 ml K2E K2EDTA Separator; PAXgene: PAXgene Blood ccfDNA Tube; RNA Streck: Cell-Free RNA BCT; Roche: Cell-Free DNA Collection Tube; Serum: BD Vacutainer SST II Advance Tube.
