## Supplementary Fig 9 for "Performance evaluation of RNA purification kits and blood collection tubes in the Extracellular RNA Quality Control (exRNAQC) study"

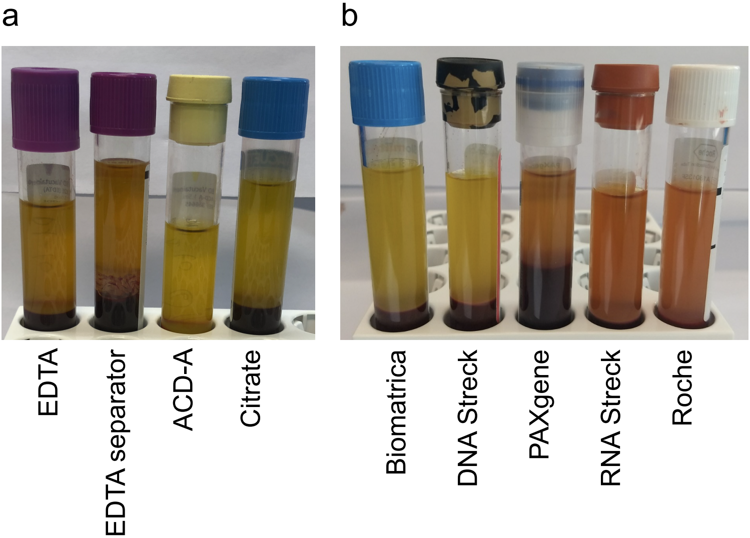


**Supplementary Fig. 9: Example of hemolysis in preservation tubes. (a)** Visual inspection of non-preservation plasma tubes of donor 7 (Supplementary Fig. 7a and Supplementary Fig. 8a) and **(b)** of preservation plasma tubes of donor 5 (Supplementary Fig. 7a and Supplementary Fig. 8a) at time interval T0. For donor 5, plasma from the PAXgene, RNA Streck and Roche tube showed to be hemolytic, which is in line with the NanoDrop measurements (Supplementary Fig. 7a and Supplementary Fig. 8a). ACD-A: BD Vacutainer Glass ACD Solution A tube; Biomatrica: LBgard Blood Tube; Citrate: Vacuette Tube 9 ml 9NC Coagulation sodium citrate 3.2%; DNA Streck: Cell-Free DNA BCT; EDTA: BD Vacutainer Plastic K2EDTA tube; EDTA separator: Vacuette Tube 8 ml K2E K2EDTA Separator; PAXgene: PAXgene Blood ccfDNA Tube; RNA Streck: Cell-Free RNA BCT; Roche: Cell-Free DNA Collection Tube.
