## Supplementary Fig 11 for "Performance evaluation of RNA purification kits and blood collection tubes in the Extracellular RNA Quality Control (exRNAQC) study"

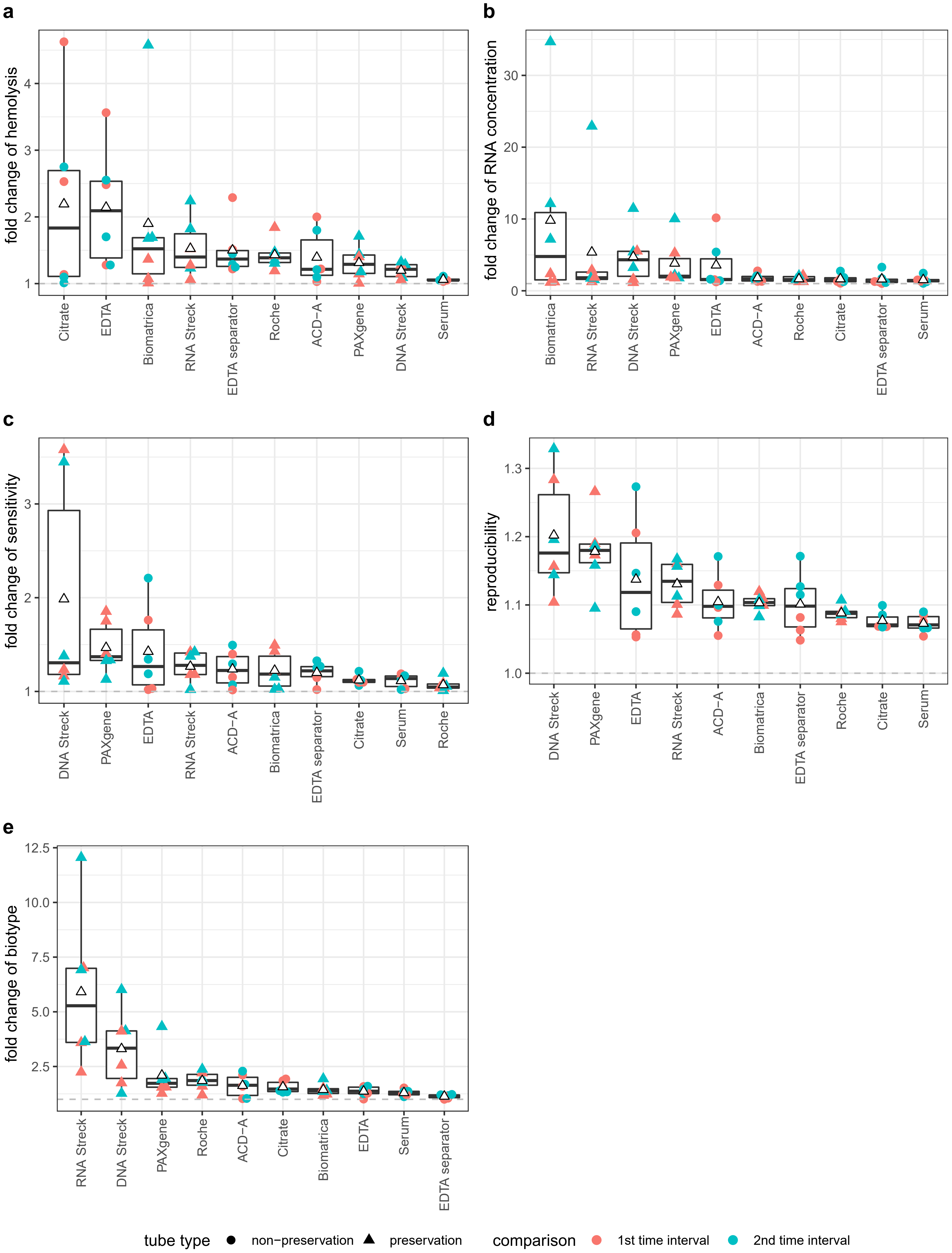


**Supplementary Fig. 11: Fold changes over time at small RNA level for each blood collection tube performance metric.** **(a)** Boxplot of the fold change within each donor across time intervals, per tube, for hemolysis, as measured by absorbance at 414 nm with Nanodrop. **(b)** Boxplot of fold change of plasma RNA concentration, based on the ratio of endogenous vs RC spike-in RNA reads. **(c)** Boxplot of the fold change of the sensitivity, i.e., the number of miRNAs after filtering out miRNAs with counts fewer than 3 reads. **(d)** Reproducibility, i.e., area left of the curve, transformed from log2 to linear scale. **(e)** Boxplot of the fold change of the fraction of the counts mapping to miRNAs versus all counts (biotype performance metric). In the boxplots, the lower and upper hinge of the boxes represents the 25th and 75th percentile, respectively. The whiskers extend to the lowest and highest value that is within 1.5 times the interquartile range. Data beyond the end of the whiskers are outliers. The white triangle on the boxplot corresponds to the mean of the fold change. Individual data points are shown as colored dots (for non-preservation tubes) or triangles (for preservation tubes). The first time interval corresponds to the comparison of T04 versus T0 (non-preservation tubes) or T24 versus T0 (preservation tubes). The second time interval corresponds to the comparison of T16 versus T0 (non-preservation tubes) or T72 vs T0 (preservation tubes). T0: plasma prepared immediately after blood draw. T04, T16, T24, T72: plasma prepared 4, 16, 24 and 72 hours after blood draw, respectively. Note that different donors were sampled and that tubes were processed at different time intervals for preservation and non-preservation tubes. ACD-A: BD Vacutainer Glass ACD Solution A tube; Biomatrica: LBgard Blood Tube; Citrate: Vacuette Tube 9 ml 9NC Coagulation sodium citrate 3.2%; DNA Streck: Cell-Free DNA BCT; EDTA: BD Vacutainer Plastic K2EDTA tube; EDTA separator: Vacuette Tube 8 ml K2E K2EDTA Separator; PAXgene: PAXgene Blood ccfDNA Tube; RNA Streck: Cell-Free RNA BCT; Roche: Cell-Free DNA Collection Tube; Serum: BD Vacutainer SST II Advance Tube.
