## Supplementary Fig 12 for "Performance evaluation of RNA purification kits and blood collection tubes in the Extracellular RNA Quality Control (exRNAQC) study"

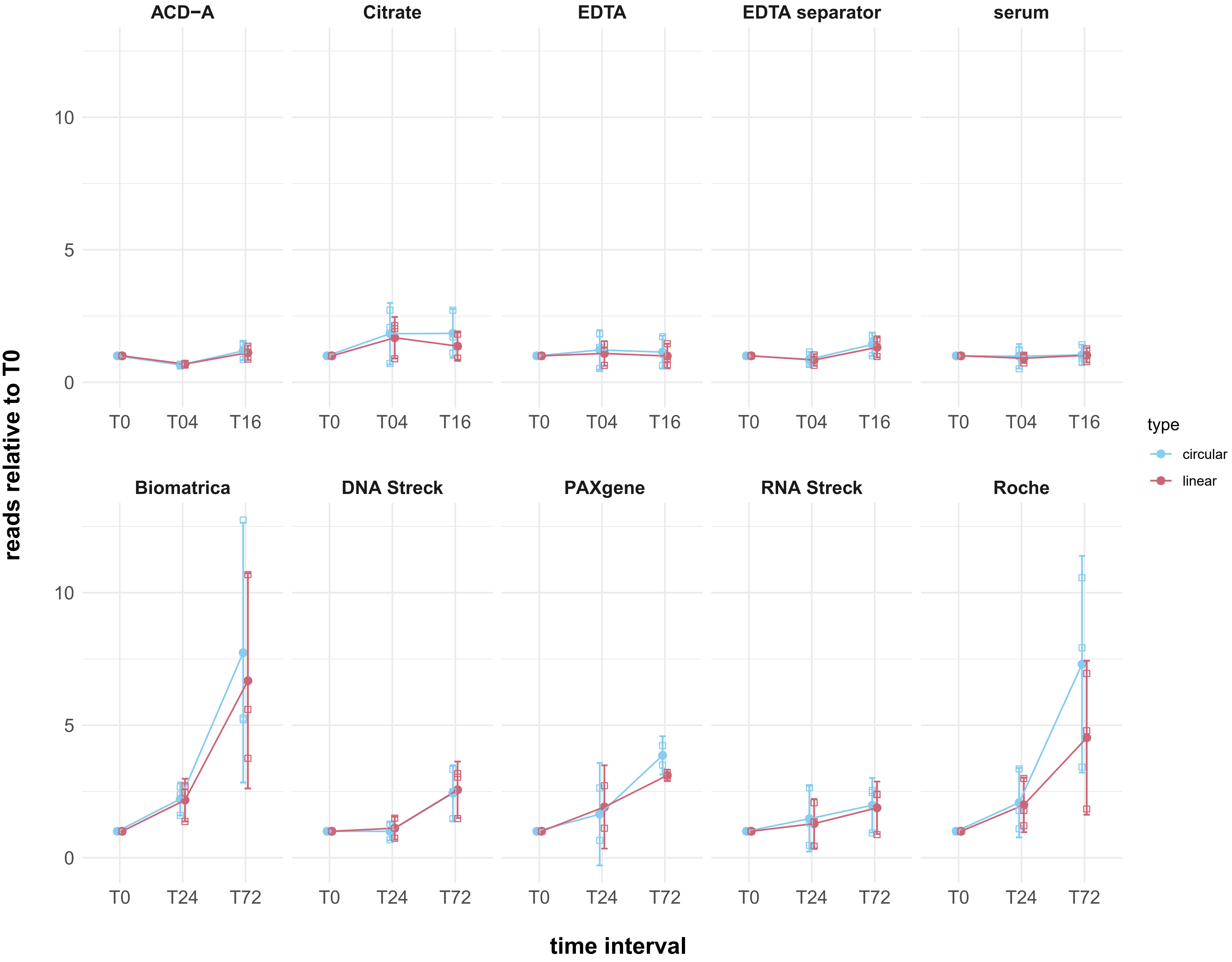


**Supplementary Fig. 12: Total number of circular and linear reads relative to T0.** Each white dot with colored outline represents the number of reads relative to the same donor's T0 tube. The fully colored dots represent the average relative number of reads and 95% confidence intervals are shown. Each plot represents a different tube type. The relative number of linear and circular reads are calculated by multiplying the fraction of the endogenous RNA reads and the Sequin spikes by the fraction of linear or circRNA reads, respectively. T0: plasma prepared immediately after blood draw. T04, T16, T24, T72: plasma prepared 4, 16, 24 and 72 hours after blood draw, respectively. Note that different donors were sampled and that tubes were processed at different time intervals for preservation and non-preservation tubes. ACD-A: BD Vacutainer Glass ACD Solution A tube; Biomatrica: LBgard Blood Tube; Citrate: Vacuette Tube 9 ml 9NC Coagulation sodium citrate 3.2%; DNA Streck: Cell-Free DNA BCT; EDTA: BD Vacutainer Plastic K2EDTA tube; EDTA separator: Vacuette Tube 8 ml K2E K2EDTA Separator; PAXgene: PAXgene Blood ccfDNA Tube; RNA Streck: Cell-Free RNA BCT; Roche: Cell-Free DNA Collection Tube; Serum: BD Vacutainer SST II Advance Tube.
