## Supplementary Fig 13 for "Performance evaluation of RNA purification kits and blood collection tubes in the Extracellular RNA Quality Control (exRNAQC) study"

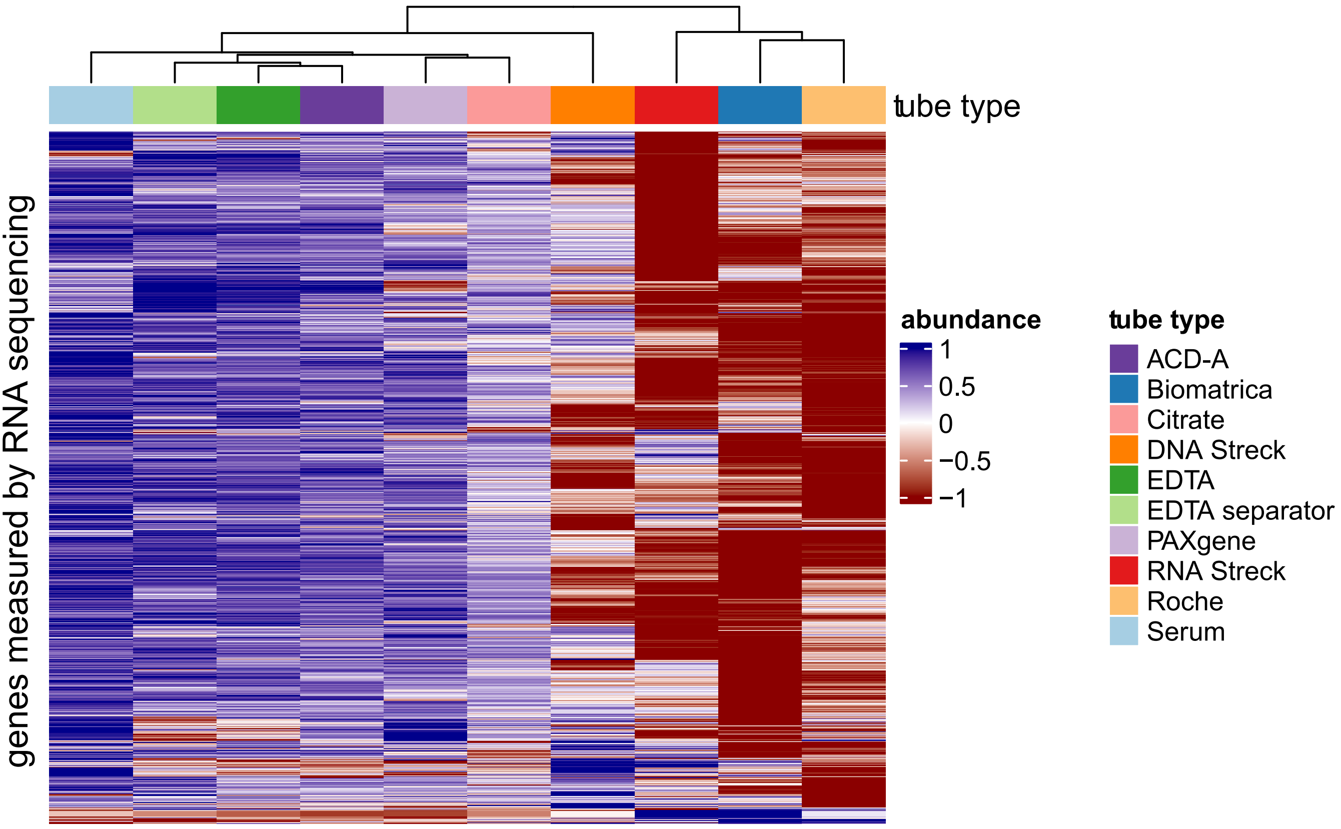


**Supplementary Fig. 13: RNA abundance levels differ across blood collection tubes.** Using normalized and scaled count data, gene abundance levels (i.e., mean abundance of the tube replicates) are shown for each tube type at time interval T0. The normalised counts are scaled between -1 (low abundance) and 1 (high abundance) per gene to make abundance differences across genes comparable. Only genes with ≥ 10 counts in all three replicates of one tube type were included. T0: plasma prepared immediately after blood draw. Note that different donors were sampled for preservation and non-preservation tubes. ACD-A: BD Vacutainer Glass ACD Solution A tube; Biomatrica: LBgard Blood Tube; Citrate: Vacuette Tube 9 ml 9NC Coagulation sodium citrate 3.2%; DNA Streck: Cell-Free DNA BCT; EDTA: BD Vacutainer Plastic K2EDTA tube; EDTA separator: Vacuette Tube 8 ml K2E K2EDTA Separator; PAXgene: PAXgene Blood ccfDNA Tube; RNA Streck: Cell-Free RNA BCT; Roche: Cell-Free DNA Collection Tube; Serum: BD Vacutainer SST II Advance Tube.
