## Supplementary Fig 14 for "Performance evaluation of RNA purification kits and blood collection tubes in the Extracellular RNA Quality Control (exRNAQC) study"

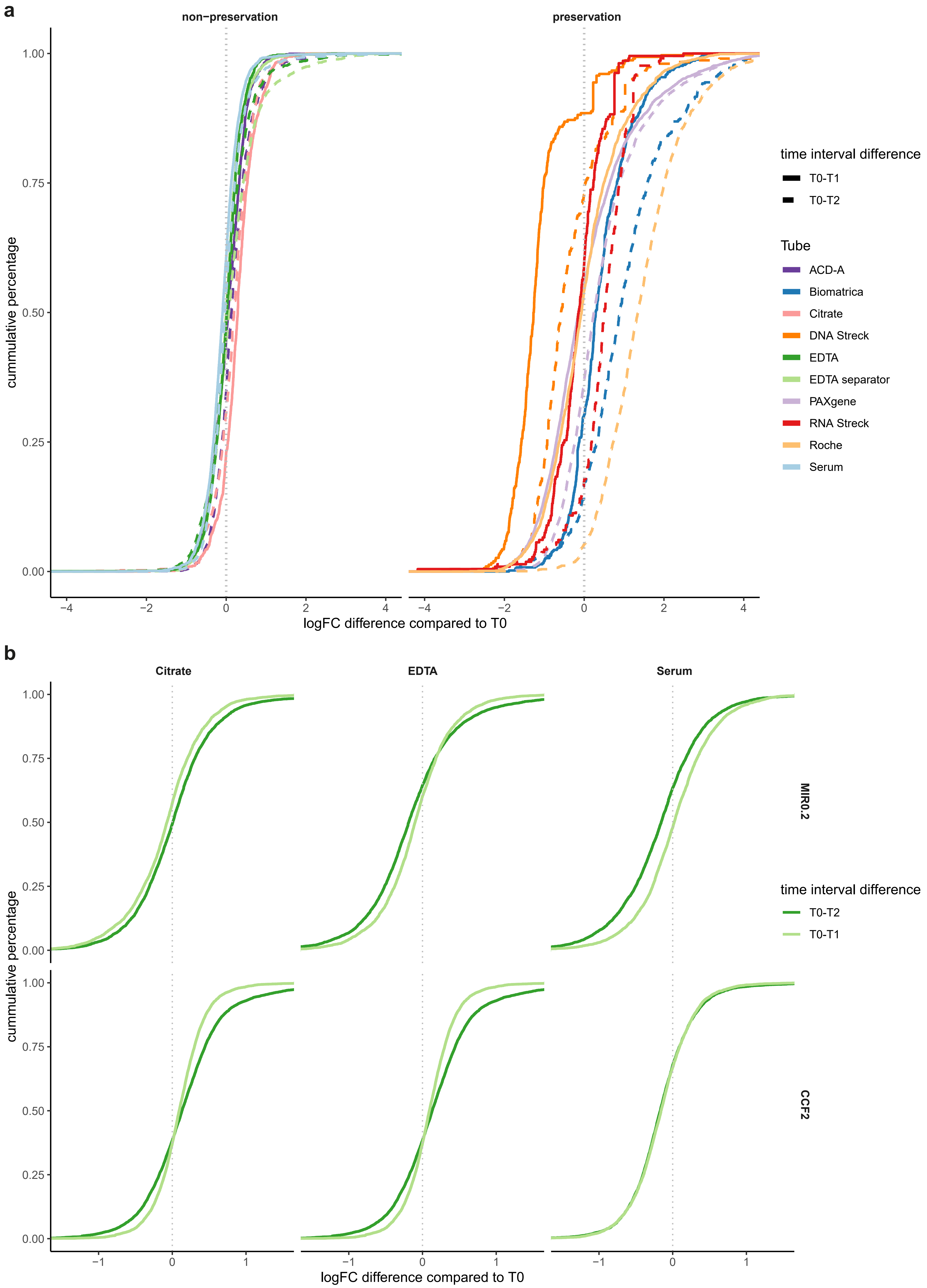


**Supplementary Fig. 14: RNA abundance levels differ across time intervals.** For each blood collection tube in exRNAQC phase 1 **(a)** and 2 **(b)**, distributions of log2 fold changes between time interval 1 and 0 (T0), and time interval 2 and 0 (T0) are shown. For the non-preservation tubes, time interval 1 corresponds to T04 and time interval 2 to T16. For the preservation tubes, time interval 1 corresponds to T24 and time interval 2 to T72. T0: plasma prepared immediately after blood draw. T04, T16, T24, T72: plasma prepared 4, 16, 24 and 72 hours after blood draw, respectively. Note that different donors were sampled and that tubes were processed at different time intervals for preservation and non-preservation tubes. ACD-A: BD Vacutainer Glass ACD Solution A tube; Biomatrica: LBgard Blood Tube; Citrate: Vacuette Tube 9 ml 9NC Coagulation sodium citrate 3.2%; DNA Streck: Cell-Free DNA BCT; EDTA: BD Vacutainer Plastic K2EDTA tube; EDTA separator: Vacuette Tube 8 ml K2E K2EDTA Separator; PAXgene: PAXgene Blood ccfDNA Tube; RNA Streck: Cell-Free RNA BCT; Roche: Cell-Free DNA Collection Tube; Serum: BD Vacutainer SST II Advance Tube.
