## Supplementary Fig 15 for "Performance evaluation of RNA purification kits and blood collection tubes in the Extracellular RNA Quality Control (exRNAQC) study"

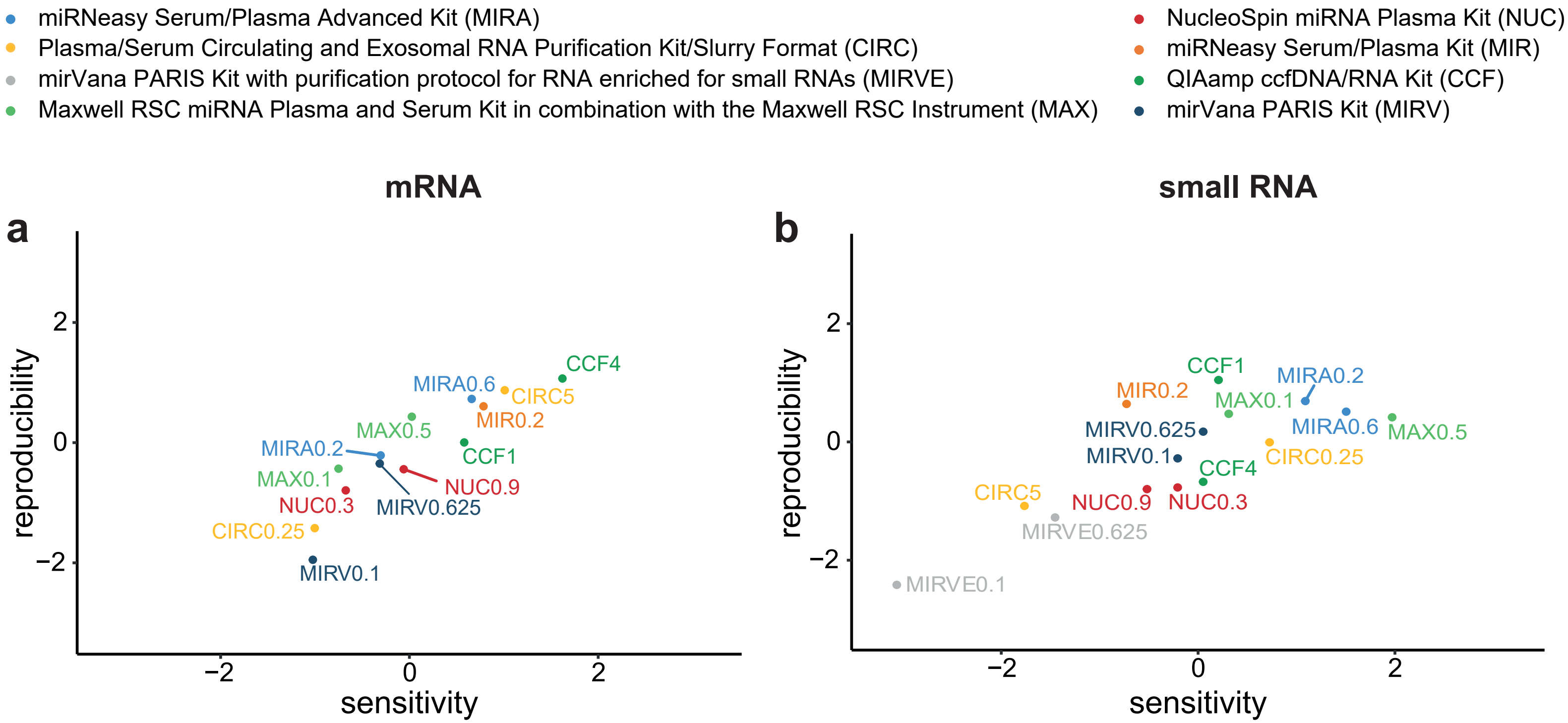


**Supplementary Fig. 15: Kit selection for exRNAQC phase 2 for mRNA capture (a) and small RNA (b) sequencing.** Median robust z-score (see Methods) per kit-input volume combination (13 in a, 15 in b) shown for sensitivity and reproducibility metrics; Number that follows the abbreviation of the purification kit is the plasma input volume (in ml).
