## Supplementary Fig 16 for "Performance evaluation of RNA purification kits and blood collection tubes in the Extracellular RNA Quality Control (exRNAQC) study"

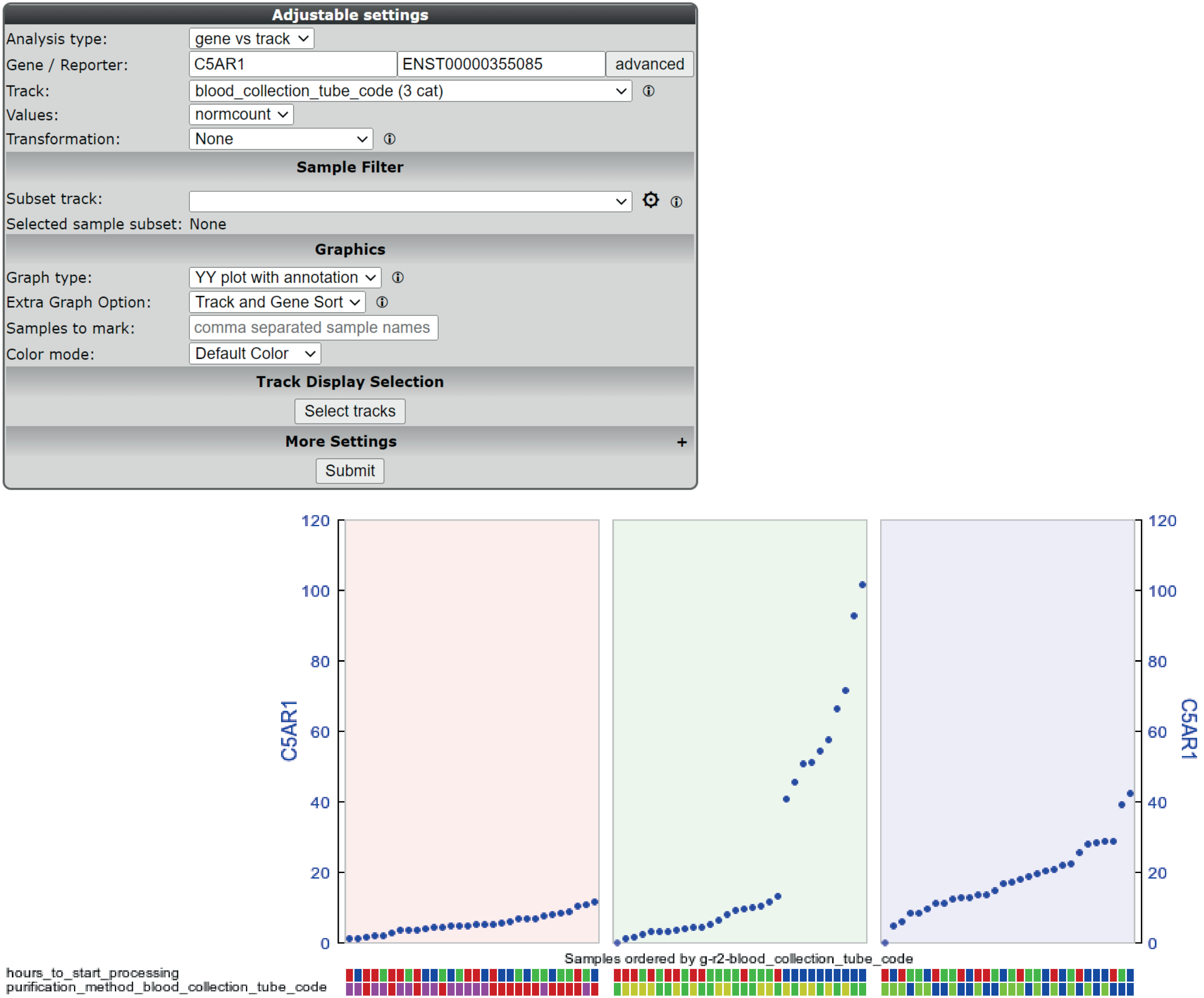


**Supplementary Fig. 16: Data access through the R2 Genomics Analysis and Visualization Platform enables browsable result access for any researcher to mine and analyze the exRNAQC data.** Shown are abundance levels of a gene (C5AR1), identified by the gene set enrichment analyses on the data of exRNAQC phase 2 as differentially abundant between time interval T0 and T16 in EDTA. Following the settings in the upper panel to 'View a gene in groups', this can be nicely visualized in R2 (lower panel). Red panel is Citrate, green is EDTA, purple is Serum. In the annotation track 'hours_to_start_processing', red represents T0, green T04, and blue T16. Mouse hover actions enable to visualize these annotations on the online platform.
