## Supplementary Information for "Performance evaluation of RNA purification kits and blood collection tubes in the Extracellular RNA Quality Control (exRNAQC) study"

**1. Donor material and biofluid preparation procedure** **3**

*1.1. Blood draws and biofluid preparations for exRNAQC study phase 1* *3*

1.1.1. Blood draws and plasma preparations for evaluation of the different exRNA purification methods 3

1.1.2. Blood draws and biofluid preparations for evaluation of the different blood collection tubes 4

*1.2. Blood draws and biofluid preparations for exRNAQC study phase 2* *8*

**2. Spike-in controls** **11**

*2.1. Sequin and External RNA Control Consortium (ERCC) spike-in controls for mRNA capture sequencing* *11*

*2.2. Capture probes for Sequin and ERCC spike-in controls* *12*

*2.3. RNA extraction Control (RC) and Library Prep Control (LP) spike-ins for small RNA sequencing* *12*

**3. RNA purification methods** **14**

*3.1. The miRNeasy Serum/Plasma Kit (abbreviated to MIR; Qiagen, 217184)* *14*

*3.2. The miRNeasy Serum/Plasma Advanced Kit (abbreviated to MIRA; Qiagen, 217204)* *15*

*3.3. The mirVana PARIS Kit (abbreviated to MIRV (and MIRVE); Life Technologies, AM1556)* *16*

*3.4. The NucleoSpin miRNA Plasma Kit (abbreviated to NUC; Macherey-Nagel, 740981.50)* *17*

*3.5. The QIAamp ccfDNA/RNA Kit (abbreviated to CCF; Qiagen, 55184)* *18*

*3.6. The Plasma/Serum Circulating and Exosomal RNA Purification Kit/Slurry Format (abbreviated to CIRC; Norgen Biotek Corp., 42800)* *19*

*3.7. The Maxwell RSC miRNA Plasma and Serum Kit (Promega, custom catalog AX5740, AS1680) in combination with the Maxwell RSC Instrument (abbreviated to MAX; Promega, AS4500)* *20*

3.7.1. Protocol AX5740 21

3.7.2. Protocol AS1680 21

*3.8. The MagNA Pure 24 Total NA Isolation Kit (Roche, 07658036001) in combination with the MagNA Pure 24 instrument (abbreviated to MAP; Roche, 07290519001)* *21*

**4. RNA concentration measurements** **22**

**5. References** **22**

### 1. Donor material and biofluid preparation procedure

#### *1.1. Blood draws and biofluid preparations for exRNAQC study phase 1*

##### *1.1.1. Blood draws and plasma preparations for evaluation of the different exRNA purification methods*

To evaluate eight different exRNA purification methods, separate blood draws were performed for each RNA sequencing workflow (mRNA or small RNA). The blood was drawn at once from each healthy donor and immediately processed to plasma. More specifically, 25 and 26 BD Vacutainer Plastic K2EDTA tubes (Becton Dickinson and Company, 367525) were collected in a single elbow venipuncture using a BD Vacutainer Push Button Blood Collection Set (Becton Dickinson and Company, 367326) for mRNA capture sequencing (experiment exRNAQC004) and small RNA sequencing (experiment exRNAQC011), respectively. All tubes were inverted 5 times to mix the anti-coagulant with the blood, immediately transported to the lab for further processing, and inverted 5 times before taking an aliquot to measure the number of platelets present in whole blood (using an XN-1000 Hematology Analyzer (Sysmex)). Next, centrifugation was performed in a Centrifuge 5804 (Eppendorf, 5804000013) with Rotor A-4-44 (Eppendorf, 5804709004) and appropriate adapters (Eppendorf, 5804753003) at acceleration and braking ramp 0. In a first centrifugation step, blood collection tubes were spun for 20 min at 400 g at room temperature, and the obtained plasma was carefully pooled, leaving ± 0.5 cm above the buffy coat. An aliquot of 7.3 ml (for mRNA capture sequencing) or 6.9 ml (for small RNA sequencing) pooled plasma was set aside and the remaining volume of plasma equally distributed across 15 ml tubes (Greiner Bio-One International, 188271) to be centrifuged for 10 min at 800 g at room temperature. After this second spin, all plasma was pooled, leaving ± 0.5 cm above the pellets and again equally distributed across 15 ml tubes. Subsequently, a third spin of 15 min at 2500 g at room temperature was applied. The plasma was again pooled, leaving ± 0.5 cm above the pellets. In total, 66 ml (for mRNA capture sequencing) or 69 ml (for small RNA sequencing) of this plasma was mixed with the corresponding plasma aliquot previously set aside. This plasma mixture was aliquoted into Safe-Lock cup DNA LoBind 2 ml PCR clean tubes (Eppendorf, 0030108078), snap frozen in liquid nitrogen and stored at -80 °C. Platelets were counted and the degree of hemolysis was determined by measuring levels of free haemoglobin by spectral analysis using a NanoDrop 1000 Spectrophotometer (Thermo Fisher Scientific).

##### *1.1.2. Blood draws and biofluid preparations for evaluation of the different blood collection tubes*

For the evaluation of ten different blood collection tubes (experiments exRNAQC005 and exRNAQC013), three separate blood draw experiments (one for each class of tubes) were planned with different donors. In each of these three experiments, three healthy donors were sampled to collect blood in either non-preservation serum tubes, non-preservation plasma tubes or preservation plasma tubes (Table 1). For collection of the serum tubes, a single puncture of an antecubital vein was performed for each of the three donors using the BD Vacutainer Push Button Blood Collection Set with pre-attached tube holder (Becton Dickinson and Company, 368657). For collection of the non-preservation plasma tubes, two punctures (one in each antecubital vein) were performed to collect 6 tubes from each arm using the BD Vacutainer Push Button Blood Collection Set with pre-attached tube holder (Becton Dickinson and Company, 368657). For collection of the preservation plasma tubes, also two punctures were performed to collect 7 tubes from one antecubital vein and 8 tubes from the contralateral antecubital vein, using a BD Vacutainer Push Button Blood Collection Set with pre-attached tube holder (Becton Dickinson and Company, 367355). Note that for donor PNL-6AJP, three DNA Streck tubes needed to be redrawn at the end of the second vein puncture, due to loss of vacuum of the first three DNA Streck tubes that were used (Table 1). Each blood draw from each puncture started with collecting 2-3 ml blood in a waste tube, followed by collection of the different tube types. The collection order of these tubes was randomized per donor. Blood tubes were filled to the volume recommended by the manufacturer and plasma tubes were inverted 5 times to mix the anti-coagulant with the blood (Table 1). Subsequently, blood collection tubes were processed immediately (T0), or 4 h (T04), 16 h (T16), 24 h (T24) or 72 h (T72) at room temperature after blood collection in order to prepare plasma or Table 1).

**Table 1. Overview of the blood draws for evaluation of the 10 different blood collection tubes.** For each of the three blood draw experiments, the draw order of the tubes per donor is given. serum: BD Vacutainer SST II Advance Tube (Becton Dickinson and Company, 366444); EDTA: BD Vacutainer Plastic K2EDTA tube (Becton Dickinson and Company, 367525); EDTA separator: Vacuette Tube 8 ml K2E K2EDTA Separator (Greiner Bio-One, 455040); ACD-A: BD Vacutainer Glass ACD Solution A tube (Becton Dickinson and Company, 366645); citrate: Vacuette Tube 9 ml 9NC Coagulation sodium citrate 3.2% (Greiner Bio-One, 455322); Roche: Cell-Free DNA Collection Tube (Roche, 07785666001); Qiagen: PAXgene Blood ccfDNA Tube (Qiagen, 768115); Biomatrica: LBgard Blood Tube (Biomatrica, M68021-001); DNA Streck: Cell-Free DNA BCT (Streck, 218996); RNA Streck: Cell-Free RNA BCT (Streck, 230248).

| blood draw experiment | donor | puncture | tube order | time to process |
|---|---|---|---|---|
| non-preservation serum | PNL-41H8 | 1 | (1) serum | T0 |
| | | | (2) serum | T16 |
| | | | (3) serum | T04 |
| | PNL-5TTC | 1 | (1) serum | T16 |
| | | | (2) serum | T04 |
| | | | (3) serum | T0 |
| | PNL-WUVM | 1 | (1) serum | T04 |
| | | | (2) serum | T16 |
| | | | (3) serum | T0 |
| non-preservation plasma | PNL-5AP8 | 1 | (1) EDTA | T0 |
| | | | (2) EDTA separator | T16 |
| | | | (3) EDTA separator | T0 |
| | | | (4) EDTA | T16 |
| | | | (5) ACD-A | T16 |
| | | | (6) ACD-A | T0 |
| | | 2 | (1) EDTA | T04 |
| | | | (2) ACD-A | T04 |
| | | | (3) citrate | T16 |
| | | | (4) citrate | T04 |
| | | | (5) citrate | T0 |
| | | | (6) EDTA separator | T04 |
| | PNL-BRCV | 1 | (1) ACD-A | T0 |
| | | | (2) EDTA | T04 |
| | | | (3) EDTA separator | T04 |
| | | | (4) EDTA | T16 |
| | | | (5) EDTA separator | T0 |
| | | | (6) citrate | T16 |
| | | 2 | (1) citrate | T0 |
| | | | (2) citrate | T04 |
| | | | (3) EDTA | T0 |
| | | | (4) ACD-A | T16 |
| | | | (5) EDTA separator | T16 |
| | | | (6) ACD-A | T04 |
| | PNL-E6AU | 1 | (1) citrate | T04 |
| | | | (2) EDTA | T04 |
| | | | (3) EDTA | T0 |
| | | | (4) EDTA separator | T0 |
| | | | (5) ACD-A | T0 |
| | | | (6) citrate | T16 |

| blood draw experiment | donor | puncture | tube order | time to process |
|---|---|---|---|---|
| | | 2 | (1) citrate | T0 |
| | | | (2) ACD-A | T16 |
| | | | (3) EDTA separator | T04 |
| | | | (4) ACD-A | T04 |
| | | | (5) EDTA | T16 |
| | | | (6) EDTA separator | T16 |
| preservation plasma | PNL-KJ6S | 1 | (1) Roche | T72 |
| | | | (2) Qiagen | T24 |
| | | | (3) Biomatrica | T0 |
| | | | (4) Roche | T0 |
| | | | (5) DNA Streck | T24 |
| | | | (6) RNA Streck | T24 |
| | | | (7) DNA Streck | T0 |
| | | 2 | (1) Qiagen | T72 |
| | | | (2) Biomatrica | T72 |
| | | | (3) Biomatrica | T24 |
| | | | (4) RNA Streck | T72 |
| | | | (5) RNA Streck | T0 |
| | | | (6) Roche | T24 |
| | | | (7) Qiagen | T0 |
| | | | (8) DNA Streck | T72 |
| | PNL-6AJP | 1 | (1) RNA Streck | T0 |
| | | | (2) Qiagen | T0 |
| | | | (3) DNA Streck | not processed |
| | | | (4) Biomatrica | T0 |
| | | | (5) Roche | T24 |
| | | | (6) Biomatrica | T24 |
| | | | (7) DNA Streck | not processed |
| | | 2 | (1) Roche | T0 |
| | | | (2) Biomatrica | T72 |
| | | | (3) RNA Streck | T24 |
| | | | (4) Roche | T72 |
| | | | (5) Qiagen | T72 |
| | | | (6) Qiagen | T24 |
| | | | (7) RNA Streck | T72 |
| | | | (8) DNA Streck | not processed |
| | | | (9) DNA Streck | T0 |
| | | | (10) DNA Streck | T24 |
| | | | (11) DNA Streck | T72 |
| | PNL-IBXE | 1 | (1) Qiagen | T0 |
| | | | (2) RNA Streck | T0 |
| | | | (3) DNA Streck | T0 |
| | | | (4) RNA Streck | T72 |
| | | | (5) Roche | T24 |
| | | | (6) Roche | T0 |
| | | | (7) RNA Streck | T24 |
| | | 2 | (1) Qiagen | T72 |
| | | | (2) Qiagen | T24 |
| | | | (3) DNA Streck | T72 |
| | | | (4) Biomatrica | T0 |
| | | | (5) DNA Streck | T24 |
| | | | (6) Roche | T72 |
| | | | (7) Biomatrica | T24 |
| | | | (8) Biomatrica | T72 |

*Non-preservation serum tubes* were processed at T0 (i.e. 30 min upon blood collection to enable full blood coagulation), T04 or T16 according to the following protocol. Until processing, the tubes were stored upright at room temperature. Tubes were spun for 10 min at 1300 g at room temperature using a Centrifuge 5804 (Eppendorf, 5804000013) with Rotor A-4-44 (Eppendorf, 5804709004) and appropriate adapters (Eppendorf, 5804753003) at acceleration and braking ramp 0. For each tube, the obtained serum was carefully pipetted into a 15 ml tube (Greiner Bio-One International, 188271), leaving ± 0.5 cm above the separator. Serum was then aliquoted into Safe-Lock cup DNA LoBind 2 ml PCR clean tubes (Eppendorf, 0030108078), snap frozen in liquid nitrogen and stored at -80 °C. Platelets were counted (only for T0) and the degree of hemolysis was determined by measuring levels of free haemoglobin by spectral analysis using a NanoDrop 1000 Spectrophotometer (Thermo Fisher Scientific).

*Non-preservation plasma tubes* were processed at T0 (i.e. immediately), T04 or T16, and *preservation plasma tubes* at T0, T24 or T72, according to the following protocol. Until processing, the tubes were stored upright at room temperature. Right before centrifugation, tubes were inverted 5 times and an aliquot to measure the number of platelets present in full blood (using an XN-1000 Hematology Analyzer (Sysmex)) was taken. Tubes were spun on a Centrifuge 5804 (Eppendorf, 5804000013) with Rotor A-4-44 (Eppendorf, 5804709004) and appropriate adapters (Eppendorf, 5804753003) at acceleration and braking ramp 0. In a first centrifugation step, blood collection tubes were spun for 20 min at 400 g at room temperature, and for each tube, the obtained plasma was carefully pipetted into a 15 ml tube (Greiner Bio-One International, 188271), leaving ± 0.5 cm above the buffy coat. Subsequently, these tubes were centrifuged for 10 min at 800 g at room temperature. After this second spin, the plasma was pipetted into new 15 ml tubes, leaving ± 0.5 cm above the pellets. Finally, a third spin of 15 min at 2500 g at room temperature was applied. The plasma was again pipetted into new 15 ml tubes, leaving ± 0.5 cm above the pellets, and aliquoted into Safe-Lock cup DNA LoBind 2 ml PCR clean tubes (Eppendorf, 0030108078), snap frozen in liquid nitrogen and stored at -80 °C. Platelets were counted and the degree of hemolysis was determined by measuring levels of free haemoglobin by spectral analysis using a NanoDrop 1000 Spectrophotometer (Thermo Fisher Scientific).

#### *1.2. Blood draws and biofluid preparations for exRNAQC study phase 2*

For each sequencing workflow, a separate blood draw experiment with five donors was performed (Table 2). Blood was collected in either serum (BD Vacutainer SST II Advance Tube; Becton Dickinson and Company, 367953), EDTA (BD Vacutainer Plastic K2EDTA tube; Becton Dickinson and Company, 367525) or citrate (Vacuette Tube 9 ml 9NC Coagulation sodium citrate 3.2%; Greiner Bio-One, 455322) tubes. For collection, a single puncture of an antecubital vein was performed for each donor using the BD Vacutainer Push Button Blood Collection Set with pre-attached tube holder (Becton Dickinson and Company, 368657), except for donor PNL-QJMM. For this donor, blood collection was briefly paused after filling six tubes and continued using a second puncture in the contralateral antecubital vein (Table 2). Each blood draw from each puncture, except the second puncture of donor PNL-QJMM, started with collecting 2-3 ml blood in a waste tube, followed by collection of the different tube types. The collection order of these tubes was randomized per donor. Blood tubes were filled to the volume recommended by the manufacturer and plasma tubes were inverted 5 times to mix the anti-coagulant with the blood. Subsequently, blood collection tubes were processed immediately (T0), or 4 h (T04) or 16 h (T16) at room temperature after blood collection in order to prepare plasma or serum (Table 2).

**Table 2. Overview of the blood draws for exRNAQC study phase 2.** For each sequencing workflow, the blood draw order of the tubes per donor is given. serum: BD Vacutainer SST II Advance Tube (Becton Dickinson and Company, 367953); EDTA: BD Vacutainer Plastic K2EDTA tube (Becton Dickinson and Company, 367525); citrate: Vacuette Tube 9 ml 9NC Coagulation sodium citrate 3.2% (Greiner Bio-One, 455322).

| blood draw experiment | donor | puncture | tube order | time to process |
|---|---|---|---|---|
| mRNA capture sequencing | PNL-RM7B | 1 | (1) citrate | T16 |
| | | | (2) EDTA | T0 |
| | | | (3) citrate | T04 |
| | | | (4) serum | T0 |
| | | | (5) citrate | T0 |
| | | | (6) EDTA | T16 |
| | | | (7) serum | T04 |
| | | | (8) EDTA | T04 |
| | | | (9) serum | T16 |
| | PNL-QJMM | 1 | (1) citrate | T0 |
| | | | (2) EDTA | T0 |
| | | | (3) serum | T16 |
| | | | (4) EDTA | T16 |

| blood draw experiment | donor | puncture | tube order | time to process |
|---|---|---|---|---|
| | | | (5) serum | T0 |
| | | | (6) EDTA | T04 |
| | | 2 | (1) citrate | T04 |
| | | | (2) citrate | T16 |
| | | | (3) serum | T04 |
| | PNL-2AAR | 1 | (1) citrate | T04 |
| | | | (2) citrate | T16 |
| | | | (3) EDTA | T0 |
| | | | (4) serum | T0 |
| | | | (5) EDTA | T16 |
| | | | (6) EDTA | T04 |
| | | | (7) serum | T04 |
| | | | (8) serum | T16 |
| | | | (9) citrate | T0 |
| | PNL-XNID | 1 | (1) serum | T0 |
| | | | (2) EDTA | T04 |
| | | | (3) citrate | T04 |
| | | | (4) EDTA | T0 |
| | | | (5) citrate | T0 |
| | | | (6) citrate | T16 |
| | | | (7) EDTA | T16 |
| | | | (8) serum | T04 |
| | | | (9) serum | T16 |
| | PNL-ZT37 | 1 | (1) EDTA | T0 |
| | | | (2) citrate | T0 |
| | | | (3) serum | T16 |
| | | | (4) serum | T04 |
| | | | (5) serum | T0 |
| | | | (6) EDTA | T16 |
| | | | (7) EDTA | T04 |
| | | | (8) citrate | T16 |
| | | | (9) citrate | T04 |
| small RNA sequencing | PNL-7DEN | 1 | (1) citrate | T16 |
| | | | (2) citrate | T04 |
| | | | (3) EDTA | T04 |
| | | | (4) serum | T04 |
| | | | (5) citrate | T0 |
| | | | (6) EDTA | T16 |
| | | | (7) EDTA | T0 |
| | | | (8) serum | T16 |
| | | | (9) serum | T0 |
| | PNL-8ZI1 | 1 | (1) EDTA | T16 |
| | | | (2) citrate | T16 |
| | | | (3) citrate | T0 |
| | | | (4) serum | T04 |
| | | | (5) EDTA | T0 |
| | | | (6) serum | T0 |
| | | | (7) serum | T16 |
| | | | (8) citrate | T04 |
| | | | (9) EDTA | T04 |
| | PNL-NLID | 1 | (1) citrate | T16 |
| | | | (2) serum | T04 |
| | | | (3) citrate | T04 |
| | | | (4) EDTA | T04 |
| | | | (5) serum | T0 |
| | | | (6) citrate | T0 |
| | | | (7) EDTA | T16 |
| | | | (8) serum | T16 |
| | | | (9) EDTA | T0 |
| | PNL-UCH7 | 1 | (1) EDTA | T0 |
| | | | (2) citrate | T0 |
| | | | (3) citrate | T16 |

| blood draw experiment | donor | puncture | tube order | time to process |
|---|---|---|---|---|
| | | | (4) serum | T04 |
| | | | (5) serum | T16 |
| | | | (6) citrate | T04 |
| | | | (7) EDTA | T16 |
| | | | (8) EDTA | T04 |
| | | | (9) serum | T0 |
| | PNL-XNID | 1 | (1) EDTA | T0 |
| | | | (2) serum | T0 |
| | | | (3) EDTA | T16 |
| | | | (4) citrate | T04 |
| | | | (5) serum | T04 |
| | | | (6) EDTA | T04 |
| | | | (7) citrate | T16 |
| | | | (8) citrate | T0 |
| | | | (9) serum | T16 |

*Serum tubes* were processed at T0 (i.e. 30 min upon blood collection to enable full blood coagulation), T04 or T16 according to the following protocol. Until processing, the tubes were stored upright at room temperature. Tubes were spun for 10 min at 1300 g at room temperature using a Centrifuge 5804 (Eppendorf, 5804000013) with Rotor A-4-44 (Eppendorf, 5804709004) and appropriate adapters (Eppendorf, 5804753003) at acceleration and braking ramp 0. For each tube, the obtained serum was carefully pipetted into a 15 ml tube (Greiner Bio-One International, 188271), leaving ± 0.5 cm above the separator. Serum was then aliquoted into Safe-Lock cup DNA LoBind 2 ml PCR clean tubes (Eppendorf, 0030108078), snap frozen in liquid nitrogen and stored at -80 °C. Platelets were counted and the degree of hemolysis was determined by measuring levels of free haemoglobin by spectral analysis using a NanoDrop 1000 Spectrophotometer (Thermo Fisher Scientific).

*Plasma tubes* were processed at T0 (i.e. immediately), T04 or T16 according to the following protocol. Until processing, the tubes were stored upright at room temperature. Right before centrifugation, tubes were inverted 5 times and an aliquot to measure the number of platelets present in whole blood (using an XN-1000 Hematology Analyzer (Sysmex)) was taken. Tubes were spun on a Centrifuge 5804 (Eppendorf, 5804000013) with Rotor A-4-44 (Eppendorf, 5804709004) and appropriate adapters (Eppendorf, 5804753003) at acceleration and braking ramp 0. In a first centrifugation step, blood collection tubes were spun for 20 min at 400 g at room temperature, and for each tube, the obtained plasma was carefully pipetted into a 15 ml tube (Greiner Bio-One International, 188271), leaving ± 0.5 cm above the buffy coat. Subsequently, these tubes were centrifuged for 10 min at 800 g at room temperature. After this second spin, the plasma was pipetted into new 15 ml tubes, leaving ± 0.5 cm above the pellets. Finally, a third spin of 15 min at 2500 g at room temperature was applied. The plasma was again pipetted into new 15 ml tubes, leaving ± 0.5 cm above the pellets, and aliquoted into Safe-Lock cup DNA LoBind 2 ml PCR clean tubes (Eppendorf, 0030108078), snap frozen in liquid nitrogen and stored at -80 °C. Platelets were counted and the degree of hemolysis was determined by measuring levels of free haemoglobin by spectral analysis using a NanoDrop 1000 Spectrophotometer (Thermo Fisher Scientific).

### 2. Spike-in controls

Adding spike-in controls is key for exRNA profiling as the RNA sequencing input is volume-based. To correct for both RNA input (either induced by the original sample or RNA isolation efficiency) and library preparation variation, two sets of spike-ins are used. RC and/or Sequin spike-ins are added during RNA isolation (upon sample lysis), and LP and/or ERCC spike-ins are added to the RNA eluate (before gDNA removal and library preparation). Depending on the amount of platelet RNA present in plasma, different spike-in concentrations are used, aiming for ± 5% reads going to the total amount of spike-ins.

#### *2.1. Sequin and External RNA Control Consortium (ERCC) spike-in controls for mRNA capture sequencing*

For profiling using the TruSeq RNA Exome Library Prep kit, Sequin spike-in controls (Garvan Institute of Medical Research, https://www.sequinstandards.com) are added to the lysate during RNA isolation. Aiming to obtain 2.5% sequencing reads aligning to the spike-in controls, a 1/1,000,000 Sequin spike-in dilution in RNase-free water was used. Per 100 µl liquid biopsy input volume, 1 µl Sequin spike-in controls (Garvan Institute of Medical Research27) was added to the lysate (see main manuscript).

The second spike-in set for profiling using the TruSeq RNA Exome Library Prep kit that we add after RNA isolation are ERCC spikes (ThermoFisher Scientific, 4456740). As for the Sequin spike-ins, we aimed to have 2.5% of the reads going to these RNA molecules. Here, a 1/500,000 dilution in RNase-free water was used. ERCC spikes were added before gDNA removal (see main manuscript for volumes).

#### *2.2. Capture probes for Sequin and ERCC spike-in controls*

To detect the spike-in controls, the capture probes of the TruSeq RNA Exome kit are complemented with capture probes for both the Sequin and ERCC spike-in sets in the first and second hybridization step of the library preparation protocol. These probes are 80-mers designed by tiling the spike-in sequences and do not map to the human genome. To this purpose, only 80-mers with a GC content between 25-70%, a GC-based Tm between 60-80 °C and a $\Delta$G larger than -7 (calculated by UNAfold (version 3.8) settings: hybrid-ss-min -E -n DNA -t 54 -T 54) were retained, and further filtered to end up with 3560 probes, i.e. the minimal number of probes needed to obtain optimal spike-in coverage. The sequences of these oligos are provided in Supplemental table 11. Here, a 70.4 ng/µl stock concentration of biotinylated 80-mer capture probes (Twist Biosciences) was used, resulting in a concentration of 4 nM per probe.

#### *2.3. RNA extraction Control (RC) and Library Prep Control (LP) spike-ins for small RNA sequencing*

For profiling using the TruSeq Small RNA Library Prep Kit, RNA extraction Control (RC) spike-ins (custom order at IDT) are added to the lysate during RNA isolation, and Library Prep Control (LP) spike-ins to the RNA eluate.

RC spike-ins are a pool of RNA extraction dynamic range controls (25-mer oligoribonucleotides; RC1-01 - RC1-12) and RNA extraction size controls (25-, 28- or 34-mer oligoribonucleotide; RC2-25 - RC2-34) selected from literature (Locati et al., Nucleic Acids Res., 2015). RC spike-in IDs and sequences are listed in Table 3.

**Table 3. Different RNA extraction Control (RC) spike-ins are used.** For each RC spike-in, the ID, sequence and relative concentration in the 10 pM RC spike-in pool are shown.

| RC spike-in IDs | sequence | relative concentration |
|---|---|---|
| RC1-01 | ACUCAUCUACGUACGCAUCUAGUCU | 0.01 x |
| RC1-03 | UGCUAUCAUAUCACAGUACGCGAGC | 0.01 x |
| RC1-04 | UAGAUGAGAUACUGAUAGCGACGUA | 0.01 x |
| RC1-06 | AUCGUCUCGUCAUCUCAUAUCUACA | 0.1 x |
| RC1-07 | UAUGCAUAUGAUCACGAGACUCAGU | 1 x |
| RC1-09 | GCUCUACACUCUACUCGUCAGCUGU | 1 x |
| RC1-10 | CGAUGCUAUAGACUCUCACGUGAUG | 1 x |
| RC1-11 | CGCAUCAGUCGUCAUCUAGAUACAG | 0.1 x |
| RC1-12 | CUGAUGAUAGAUACGCGCACACAGU | 0.1 x |
| RC2-25 | AUGCUGAUGAUAGACGCUACUGACU | 0.1 x |
| RC2-28 | CGUAUCGCUGCUCUGAGUCACUAUCUAC | 0.1 x |
| RC2-34 | AGAUAGUACUGAUCUGCUGCGACGAGUGACUGUC | 0.1 x |

LP spike-ins are a pool of small RNA sequencing library prep controls (22-mer oligoribonucleotides; LP1-01 - LP1-12) selected from literature (Hafner et al., RNA, 2011). LP spike-in IDs and sequences are listed in Table 4.

**Table 4. Different Library Prep Control (LP) spike-ins are used.** For each LP spike in, the ID, sequence and relative concentration in the 10 pM LP spike-in pool are shown.

| LP spike-in IDs | sequence | relative concentration |
|---|---|---|
| LP1-01 | GUCCCACUCCGUAGAUCUGUUC | 1 x |
| LP1-02 | GAUGUAACGAGUUGGAAUGCAA | 0.01 x |
| LP1-03 | UAGCAUAUCGAGCCUGAGAACA | 0.1 x |
| LP1-04 | CAUCGGUCGAACUUAUGUGAAA | 0.01 x |
| LP1-06 | UCUUAACCCGGACCAGAAACUA | 1 x |
| LP1-07 | AGGUUCCGGAUAAGUAAGAGCC | 1 x |
| LP1-10 | UGAUACGGAUGUUAUACGCAGC | 0.1 x |
| LP1-11 | CCUGGAACUUAGGACGUGAAUC | 0.1 x |
| LP1-12 | UCAUGAGUCCGUACCUUGAUUG | 0.01 x |

RC and LP spike-ins concentrations were optimized, aiming to obtain 2.5% sequencing reads aligning to the spike-in controls. To this purpose, RC and LP spike-ins were dissolved to 200 µM stock solutions using nuclease-free water (Sigma-Aldrich, W4502), and equimolarly pooled to 333 nM. Subsequently, 6.25 pM, 625 fM and 62.5 fM pools were created using a 500 nM carrier oligo (TCGAAGTATTC; diluted in nuclease-free water) to dilute the initial 333 nM pool. These three pools were spiked into plasma during RNA isolation, by adding 2 µl to the lysate, followed by TruSeq Small RNA Library Prep sequencing. Based on these sequencing data, a separate RC spike-in and LP spike-in pool was created, in which each RNA control is diluted at a different concentration, in order to correct for adaptor ligation bias during library preparation. To this purpose, the RC and LP spike-in stock solutions were diluted to 5 µM using nuclease-free water, and pooled into a ligation bias-corrected 10 pM RC spike-in pool and 10 pM LP spike-in pool, respectively, using 500 nM carrier oligo. The indicated 10 pM concentration of these pools corresponds to the concentration of RC2-34, which has the highest ligation efficiency. Concentrations of the remaining spikes are relative to the RC2-34 concentration (Table 3). Finally, using these ligation-bias corrected 10 pM pools and 500 nM carrier oligo, RC and LP spike-in pools were made. The final ligation bias-corrected concentration of RC spike-in pool was 1259 fM for the kit comparison study (exRNAQC011), and 191 fM for the tube comparison study (exRNAQC013) and phase 2 (exRNAQC017 small RNA sequencing). The final ligation bias-corrected concentration of LP spike-in pool was 486 fM for the kit comparison study (exRNAQC011), and 34 fM for the tube comparison study (exRNAQC013) and phase 2 (exRNAQC017 small RNA sequencing). Per 100 µl liquid biopsy input volume, 1 µl RNA extraction Control (RC) spike-ins was added to the lysate during RNA purification. LP spikes were added before gDNA removal (see main manuscript for volumes).

### 3. RNA purification methods

#### *3.1. The miRNeasy Serum/Plasma Kit (abbreviated to MIR; Qiagen, 217184)*

All RNA purifications throughout the study are performed using the miRNeasy Serum/Plasma Kit, unless specified otherwise. For evaluation of the different exRNA purification methods, RNA purifications were performed in triplicate and 200 µl plasma was used per RNA purification, as the manufacturer’s manual indicates a maximum recommended biofluid input volume of 200 µl; required minimum volumes are not mentioned.

Plasma is thawed on ice and 1000 µl QIAzol Lysis Reagent is added to each sample. Samples are vortexed and incubated for 5 min at room temperature, followed by the addition of Sequin control spike-ins and/or RC RNA extraction Control (RC) spike-ins. Subsequently, samples are vortexed and 200 µl chloroform is added, followed by vortexing of the lysates for 15 s. After a 2 min incubation at room temperature, samples are centrifuged for 15 min at 12000 g at 4 °C. Next, 600 µl of the upper aqueous phase is transferred to a new collection tube on ice, to which 900 µl ethanol is pipetted. Samples are mixed by pipetting up and down, loaded (up to 700 µl) on an RNeasy MinElute spin column and centrifuged for 15 s at 10000 g at room temperature. The flowthrough is discarded, and loading and centrifugation repeated using the remainder of the samples. Afterwards, 700 µl RWT buffer is added to the column and samples are centrifuged for 15 s at 10000 g. Flowthroughs are discarded and 500 µl RPE buffer is pipetted onto the columns, followed by centrifugation for 15 s at 10000 g. Again, flowthroughs are discarded, and 500 µl 80 % ethanol is loaded onto the columns. Samples are centrifuged for 2 min at 10000 g, and afterwards, the columns are placed into a new collection tube (with open lid) and dried for 5 min at full speed (16900 g). Finally, the columns are placed in a new collection tube, 14 µl RNase-free water (Sigma, W4502) is added to the center of the column membrane, and RNA is eluted by centrifugation for 1 min at full speed.

#### ***3.2. The miRNeasy Serum/Plasma Advanced Kit (abbreviated to MIRA; Qiagen, 217204)***

For evaluation of the different exRNA purification methods, RNA purifications were performed in triplicate, and a minimum and maximum input volume of 200 µl and 600 µl plasma was used, respectively. In phase 2, 600 µl plasma input volume was used.

Plasma is thawed on ice and 60 µl Buffer RPL is added per 200 µl of plasma input volume. Samples are vortexed for 5 s and left at room temperature for 3 min, followed by the addition of Sequin control spike-ins and/or RC RNA extraction Control (RC) spike-ins. Per 200 µl plasma input volume, 20 µl RPP Buffer is added. Samples are vortexed for >20 s, incubated at room temperature for 3 min, and centrifuged at 12000 g for 3 min. Per 200 µl plasma input volume, 220 µl of the clear and colourless supernatant is transferred to a new tube and 1 volume of isopropanol is added, and tubes are vortexed. The entire sample (up to 700 µl) is transferred to a RNeasy UCP MinElute column and centrifuged for 15 s at 10000 g, and loading and centrifugation repeated with the remainder of the samples (only for the maximum plasma input volume). Flowthroughs are discarded and 700 µl Buffer RWT is added onto the column. Columns are again centrifuged for 15 s at 10000 g and flowthroughs discarded. Next, 500 µl buffer RPE is added onto the column and samples are centrifuged for 15 s at 10000 g. Flowthroughs are discarded and 500 µl 80 % ethanol is added to the column. Columns are centrifuged for 2 min at 10000 g and placed in a new collection tube. Columns are centrifuged at full speed (16900 g) for 5 min, with open lid to dry the membrane. Dry columns are placed in a new 1.5 ml collection tube and 20 µl RNase-free water (Sigma, W4502) is added directly to the center of the spin column membrane and incubated for 1 min. Next, columns are centrifuged for 1 min at full speed (16900 g) to elute the RNA.

#### *3.3. The mirVana PARIS Kit (abbreviated to MIRV (and MIRVE); Life Technologies, AM1556)*

For evaluation of the different exRNA purification methods, RNA purifications were performed in triplicate, and a minimum and maximum biofluid input volume of 100 µl and 625 µl was used, respectively. Although not explicitly stated by the manufacturer, the minimum input volume was set on 100 µl based on the manufacturer's indication that smaller sample volumes need to be diluted to 100 µl with Cell Disruption Buffer.

Plasma is thawed on ice and added to an equal volume of 2x Denaturing Solution at room temperature. This mixture is incubated for 5 min on ice and Sequin control spike-ins and/or RC RNA extraction Control (RC) spike-ins are added, followed by adding a volume of Acid-Phenol:Chloroform equal to the total lysate volume. Samples are mixed by vortexing for 60 s and centrifuged for 5 min at 10000 g at room temperature to separate the mixture into aqueous and organic phases. The aqueous phase (i.e. 140 µl and 650 µl for the minimum and plasma input volume, respectively) is recovered and transferred to a fresh tube. Subsequently, 1.25 volumes of 100 % ethanol are added to the aqueous phase, and the mixed sample (up to 700 µl) is pipetted onto a Filter Cartridge and centrifuged for 30 s (all centrifugation steps are at 10000 g). Flowthroughs are discarded, and loading on the Filter Cartridge and centrifugation repeated using the remainder of the samples. Filter Cartridges are washed by applying 700 µl miRNA Wash Solution 1 and centrifuging for 15 s. The flowthrough is discarded. Next, samples are washed twice by applying 500 µl Wash Solution 2/3 and centrifuging for 15 s. After discarding the flowthrough from the last wash, the Filter Cartridge is replaced in the Collection Tube and spun for 1 min to remove residual fluid from the filter. To elute the RNA, 100 µl of preheated Elution Solution is pipetted to the center of the filter, placed in a new Collection Tube, and samples are centrifuged for 30 s.

For evaluation of the different exRNA purification methods for small RNA sequencing, also an alternative protocol claiming to enrich for small RNAs (abbreviated to MIRVE) was tested. The first steps are identical to the purification protocol described above. After recovering the aqueous phase, 1/3 volume of 100 % ethanol is added and mixed with the lysate. Then, the mixture (up to 700 µl) is pipetted onto a Filter Cartridge and centrifuged for 30 s. The filtrate is transferred to a fresh tube. These steps are repeated with the remainder of the sample. Filtrates are pooled and the total volume of filtrate is determined. Next, 2/3 volume of room temperature 100 % ethanol is added to the filtrate, and the sample is mixed thoroughly. This mixture is passed through a second Filter Cartridge. This time, the flowthrough is discarded, and the Filter Cartridge is washed and RNA eluted as described for the purification protocol above (i.e. MIRV).

#### *3.4. The NucleoSpin miRNA Plasma Kit (abbreviated to NUC; Macherey-Nagel, 740981.50)*

For evaluation of the different exRNA purification methods, RNA purifications were performed in triplicate, and a minimum and maximum biofluid input volume of 300 µl and 900 µl was used, respectively.

Plasma is thawed on ice and 90 µl MLP Buffer per 300 µl input volume is added. Samples are vortexed for 5 s and incubated for 3 min at room temperature. Sequin control spike-ins and/or RC RNA extraction Control (RC) spike-ins are added to the lysate. Next, 30 µl MPP Buffer per 300 µl plasma input volume is added. Samples are vortexed for 5 s, incubated for 1 min at room temperature and centrifuged for 3 min (all centrifugation steps are at 11000 g). The clear supernatant (i.e. 250 µl and 1100 µl for the minimum and maximum input volume, respectively) is transferred into a new Collection Tube and per 300 µl plasma input volume 400 µl isopropanol is added. Samples are vortexed for 5 s, loaded onto a NucleoSpin miRNA Column, incubated for 2 min at room temperature and centrifuged for 30 s. Flowthroughs are discarded and loading repeated with the remainder of the samples. Next, the columns are washed by adding 100 µl Buffer MW1 and centrifuging for 30 s. Flowthroughs are discarded and columns washed a second and third time by adding 700 µl Buffer MW2 and centrifuging for 30 s, and adding 250 µl Buffer MW2 and centrifuging for 2 min, respectively. Subsequently, the column is placed into a new Collection Tube, and 30 µl RNase-free water (Sigma, W4502) pipetted onto the silica membrane. Samples are incubated for 1 min at room temperature and centrifuged for 1 min to elute the RNA.

#### ***3.5. The QIAamp ccfDNA/RNA Kit (abbreviated to CCF; Qiagen, 55184)***

For evaluation of the different exRNA purification methods, RNA purifications were performed in triplicate, and a minimum and maximum input volume of 1000 µl and 4000 µl plasma was used, respectively. In phase 2, 2000 µl plasma input volume was used.

Plasma is thawed on ice, transferred to a 15 ml collection tube and 300 µl Buffer RPL is added for each 1000 µl of plasma. Samples are vortexed for 5 s and left at room temperature for 3 min, followed by the addition of Sequin control spike-ins and/or RC RNA extraction Control (RC) spike-ins. Per 1000 µl plasma input volume, 100 µl Buffer RPP is added. Samples are vortexed for >20 s and incubated on ice for 3 min. Proteins are precipitated by centrifuging the samples at 3000 g for 10 min. The clear and colourless supernatant (i.e. 1100 µl and 4400 µl for the minimum and maximum plasma input volume, respectively) is transferred to a new tube (on ice), 1 volume of ice-cold isopropanol is added and the tubes are vortexed. Up to 4000 µl sample is transferred to an RNeasy Midi spin column and the column is centrifuged at room temperature for 1 min at 3000 g. Flowthroughs are discarded, and loading repeated with the remainder of the samples. Next, 4000 µl Buffer RWT is added to the column. Columns are centrifuged for 1 min at 3000 g and flowthroughs are discarded, followed by the addition of 2500 µl Buffer RPE and centrifugation for 5 min at 3000 g. After placing the columns into a new 15 ml collection tube, 200 µl RNase-free water (Sigma, W4502) is added directly to the center of the membrane and columns are incubated for 1 min. Next, columns are centrifuged for 1 min at full speed (4500 g) to elute the RNA. Subsequently, 200 µl Buffer RPL and 800 µl 100 % ethanol are added to the eluate and samples are mixed by pipetting up and down. Up to 700 µl sample is pipetted onto an RNeasy MinElute spin column. Columns are centrifuged at 10000 g for 15 s at room temperature, flowthroughs discarded and loading repeated with the remainder of the samples. Subsequently, 500 µl Buffer RPE is pipetted onto the column, followed by centrifuging the columns for 15 sec at 10000g. Flowthroughs are discarded and 500 µl 80 % ethanol is added to the columns. The columns are again centrifuged for 15 s at 10 000 g and placed in a fresh collection tube, followed by centrifugation for 5 min at full speed (16900 g), with open lid to dry the membrane. Then, the columns are placed in a new 1.5 ml collection tube and 14 µl RNase free water (Sigma, W4502) is added directly to the center of the spin column membrane. Finally, columns are centrifuged for 1 min at full speed (16900 g) to elute the RNA.

#### *3.6. The Plasma/Serum Circulating and Exosomal RNA Purification Kit/Slurry Format (abbreviated to CIRC; Norgen Biotek Corp., 42800)*

For evaluation of the different exRNA purification methods, RNA purifications were performed in triplicate, and a minimum and maximum biolfuid input volume of 250 µl and 5000 µl was used, respectively. Note that depending on the plasma input volume that is used, the manufacturer's manual instructs to use different volumes of Lysis Buffer A and 100 % ethanol. Here, we provide the protocol to specifically process 250 µl plasma. Adjusted volumes to process 5000 µl plasma are indicated between brackets. In addition, note that in the manufacturer's manual centrifugation speeds are quoted in rpm. The indicated speeds thus depend on the radius of the centrifuge rotor. Here, a Centrifuge 5804 (Eppendorf, 5804000013) with Rotor A-4-44 (Eppendorf, 5804709004) was used at the start of the protocol. As soon as samples were loaded onto spin columns (see protocol below), centrifugation steps were performed using a Centrifuge 5424R (Eppendorf, 5404000618) with Rotor FA-45-24-11 (Eppendorf, 5424700004).

Plasma is thawed on ice and 200 µl Slurry C2 and 300 µl (9800 µl) Lysis Buffer A is added. Samples are mixed by vortexing for 15 s. After incubation for 10 min at 60 °C, Sequin control spike-ins and/or RC RNA extraction Control (RC) spike-ins are added, as well as 750 µl (15000 µl) 100 % ethanol, and samples are vortexed for 15 s, followed by centrifugation for 30 s at 1000 RPM (all centrifugation steps are at room temperature). The supernatant is carefully decanted and 300 µl Lysis Buffer A is added to the pellet. Samples are mixed well by vortexing for 15 s and incubated for 10 min at 60 °C. Then, 300 µl 100 % ethanol is added and the mixture is vortexed for 15 s. Next, the samples (up to 650 µl) are loaded onto a Mini Filter Spin column and centrifuged for 1 min at 14000 RPM. Flowthroughs are discarded. This loading and centrifugation step is repeated with the remainder of the samples. Subsequently, 400 µl Wash Solution A is applied to the column, followed by centrifugation for 1 minute at 14000 RPM and discarding the flowthrough. This wash step is repeated two more times, for a total of three washes. Columns are spun empty, for 3 min at 14000 RPM and transferred to a fresh Elution tube. To elute the RNA, 100 µl Elution Solution A is applied to the column and samples are centrifuged for 2 min at 2000 RPM, followed by 3 min at 14000 RPM.

#### *3.7. The Maxwell RSC miRNA Plasma and Serum Kit (Promega, custom catalog AX5740, AS1680) in combination with the Maxwell RSC Instrument (abbreviated to MAX; Promega, AS4500)*

For evaluation of the different exRNA purification methods, RNA purifications were performed in triplicate, and a minimum and maximum biofluid input volume of 100 µl and 500 µl was used, respectively. At the time the exRNAQC study was set up, the Maxwell RSC miRNA Plasma and Serum Kit was not yet commercially available, and purifications were performed using custom catalog number AX5740, received from the company. To test the interactions between pre-analytics, the commercially available kit (AS1680) was used. Note that these two versions of the kit have similar components, except for the Maxwell RSC cartridges. The difference between cartridges is that the commercially available cartridge (AS1680) uses a newer magnetic purification cellulose resin and seal stock material (e-mail communication Promega).

##### *3.7.1. Protocol AX5740*

Plasma is thawed on ice and 80 µl Proteinase K and 230 µl Binding Buffer is added. Samples are vortexed for 10 s and incubated for 15 min at 37 °C. During this incubation step, the RSC Cartridges are prepared as follows. The cartridges are placed in the RSC deck trays with well #1 facing away from the Elution Tubes, and snapped into position by pressing down on the cartridges. Seals are removed and a RSC Plunger is placed into well #8 of each cartridge. Sequin control spike-ins and/or RC RNA extraction Control (RC) spike-ins are added to well #1 and 50 µl Nuclease-Free Water to each Elution Tube. After the incubation step, the lysate is added to well #1. Subsequently samples are loaded onto the instrument and the automated purification run is started according to the Maxwell RSC miRNA method.

##### *3.7.2. Protocol AS1680*

Plasma is thawed on ice and 80 µl Proteinase K and 230 µl Lysis Buffer C is added. Samples are vortexed for 5 s and incubated for 15 min at 37 °C. During this incubation step, the RSC Cartridges are prepared as follows. The cartridges are placed in the RSC deck trays with well #1 facing away from the Elution Tubes, and snapped into position by pressing down on the cartridges. Seals are removed and a RSC Plunger is placed into well #8 of each cartridge. Sequin control spike-ins and/or RC RNA extraction Control (RC) spike-ins are added to well #1 and 50 µl Nuclease-Free Water to each Elution Tube. After the incubation step, the lysate is added to well #1. Subsequently samples are loaded onto the instrument and the automated purification run is started according to the miRNA Plasma and Serum method.

#### ***3.8. The MagNA Pure 24 Total NA Isolation Kit (Roche, 07658036001) in combination***

***with the MagNA Pure 24 instrument (abbreviated to MAP; Roche, 07290519001)***

For evaluation of the different exRNA purification methods, a minimum and maximum biofluid input volume of 2000 µl and 4000 µl was used, respectively. As recommended by the manufacturer, we made use of the cfNA ss 2000 protocol for 2000 µl samples and the cfNA ss 4000 protocol for 4000 µl samples.

Plasma is thawed on ice and aliquoted in volumes of 1050 µl into 1.5 ml microcentrifuge tubes. To each tube, 105 µl proteinase K is added and samples are incubated for 20 min at 37 °C. After incubation, 1000 µl of each microcentrifuge tube is pooled into a Falcon round bottomed test tube (VWR, 734-0446) to obtain 2000 µl and 4000 µl input volumes. Next, cfNA buffer mix is prepared in bulk by mixing 1750 µl Cell-Free Nucleic Acid Enhancement Buffer (CELB) with 300 µl Isopropanol (IPA) per 2000 µl sample. Of this cfNA buffer mix, 2000 µl and 4000 µl is added to the 2000 µl and 4000 µl input samples, respectively, followed by the addition of Sequin control spike-ins and/or RC RNA extraction Control (RC) spike-ins. Samples are thoroughly mixed by dispensing and aspirating the liquid 8 times to produce a homogeneous mixture, and centrifuged at 1400 g for 1 minute. Remaining bubbles were removed with the back of a tip. The MagNA Pure 24 instrument was loaded as described in the manufacturer's manual and samples were eluted in 50 µl.

### 4. RNA concentration measurements

Eluate RNA concentrations are measured using the Femto Pulse system (Agilent Technologies, M5330AA) with the Ultra Sensitivity RNA Kit (Agilent Technologies, FP-1201-0275) according to the manufacturer's instructions.
